# Advancing motion denoising of multiband resting-state functional connectivity fMRI data

**DOI:** 10.1101/860635

**Authors:** John C. Williams, Philip N. Tubiolo, Jacob R. Luceno, Jared X. Van Snellenberg

## Abstract

Simultaneous multi-slice (multiband) accelerated functional magnetic resonance imaging (fMRI) provides dramatically improved temporal and spatial resolution for resting-state functional connectivity (RSFC) studies of the human brain in health and disease. However, multiband acceleration also poses unique challenges for denoising of subject motion induced data artifacts, the presence of which is a major confound in RSFC research that substantively diminishes reliability and reproducibility. We comprehensively evaluated existing and novel approaches to volume censoring-based motion denoising in the Human Connectome Project (HCP) dataset. We show that assumptions underlying common metrics for evaluating motion denoising pipelines, especially those based on quality control-functional connectivity (QC-FC) correlations and differences between high- and low-motion participants, are problematic, and appear to be inappropriate in their current widespread use as indicators of comparative pipeline performance and as targets for investigators to use when tuning pipelines for their own datasets. We further develop two new quantitative metrics that are instead agnostic to QC-FC correlations and other measures that rely upon the null assumption that no true relationships exist between trait measures of subject motion and functional connectivity, and demonstrate their use as benchmarks for comparing volume censoring methods. Finally, we develop and validate quantitative methods for determining dataset-specific optimal volume censoring parameters prior to the final analysis of a dataset, and provide straightforward recommendations and code for all investigators to apply this optimized approach to their own RSFC datasets.

## 1. Introduction

### 1.1. Background

The study of resting-state functional connectivity (RSFC) with functional Magnetic Resonance Imaging (fMRI) has become the dominant approach to studying the connectome of the human brain, a key priority of the National Institute of Mental Health for achieving their strategic objective to define the mechanisms of complex behaviors (NIMH, 2020). However, it is widely recognized that participant motion represents a serious confound in RSFC research, and that aggressive steps must be taken to mitigate its impact (Ciric et al., 2017; Parkes et al., 2018; Power et al., 2012; Power et al., 2014; Power et al., 2015; Power et al., 2019b; Satterthwaite et al., 2013; Satterthwaite et al., 2012; Van Dijk et al., 2012; Yan et al., 2013). The success of this ongoing effort will be crucial for the reliability and reproducibility of RSFC fMRI studies, and the use of fMRI as a tool for understanding human connectomics in both health and disease.

In recent years, simultaneous multi-slice (multiband) acceleration of fMRI sequences (Moeller et al., 2010; Ugurbil et al., 2013) has gained prominence, and has been adopted by several large-scale studies of human brain function, including the Human Connectome Project (HCP; Ugurbil et al., 2013), the UK Biobank study (Miller et al., 2016), and the Adolescent Brain Cognitive Development (ABCD) study (Casey et al., 2018; Hagler et al., 2019; Jernigan et al., 2018; Volkow et al., 2018). Multiband acceleration provides substantial improvements in both spatial and temporal resolution of fMRI data. However, data acquired with sub-second repetition times (TRs), as enabled by multiband acceleration, appear to contain not just traditionally recognized markers of motion artifact (e.g., net elevations in observed RSFC correlations, with both spatially-dependent and spatially-independent components), but also novel artifacts not observed in traditional, higher-TR single-band data. It has been previously observed in both the ABCD and HCP studies that both true respiratory motion and pseudomotion (factitious head motion, observed primarily in the phase-encode direction as a result of tissue changes due to lung expansion impacting the B_0_ field) are evident in multiband fMRI due to the higher sampling rate of these sequences relative to traditional single-band fMRI (Burgess et al., 2016; Fair et al., 2020; Glasser et al., 2018; Power et al., 2019a), findings which parallel others’ observations as well as our own (see **Figure S1**).

The discovery of novel forms of motion and pseudomotion artifact (hereafter, “motion artifact,” except where this distinction is of note) in fast-TR multiband data underscores a critical need to comprehensively evaluate methods for characterizing and removing motion artifact in these data. Previously, Burgess and colleagues (Burgess et al., 2016) evaluated the performance of volume censoring, alone and in tandem with other methods, in the HCP multiband dataset using a single combined set of framewise displacement (FD) and DV (temporal derivative root-mean-squared variance over voxels) censoring thresholds. It was noted that while volume censoring reduced spatially specific artifacts, its performance was significantly impaired in the context of multiband data, at least in part due to the aforementioned presence of respiratory motion (Power et al., 2019a). However, notch filtering of motion estimates prior to calculation of the subject-level quality control (QC) measure FD has been shown in other work to lead to improved performance of volume censoring pipelines (Fair et al., 2020; Power et al., 2019a), although this work did not evaluate similarly modified measures of DV.

Thus, while it seems clear that volume censoring behaves differently in fast-TR multiband datasets than it does in singleband datasets, and some improvement in volume censoring methods has been made over existing singleband methods (Fair et al., 2020; Power et al., 2019a), we suspected that substantive improvement in these methods may still be possible. For example, while band-stop (notch) filtering of motion parameters (MPs) improves FD-based volume censoring (Fair et al., 2020; Power et al., 2019a), such filters have not been attempted for DV-based censoring (widely used alongside FD censoring), and filtering changes the magnitude of FD values, raising the possibility that common FD censoring thresholds (e.g., of 0.2 or 0.5 mm; see (Power et al., 2014) may no longer be appropriate.

Critically, it should be noted that volume censoring is not the only widely used method for addressing motion-related confounds in RSFC datasets, with ICA-based methods such as ICA-FIX (Griffanti et al., 2014), ICA-AROMA (Pruim et al., 2015), and temporal ICA (Glasser et al., 2018) making up the other primary class of methods used in the literature. We do not attempt to address here whether volume censoring, as a class of approaches, outperforms ICA-methods either individually or as a group; rather, our work is focused on optimizing the application of volume censoring for those investigators who opt for this approach to mitigating the impact of participant motion and related confounds on RSFC datasets.

### 1.2. Overview

We began this work with the goal of evaluating the enhancement in denoising performance produced by two potential improvements to volume censoring methods for use with multiband fMRI data: 1) low-pass filtering MPs and voxel time series at 0.2 Hz prior to calculating FD and DV, to produce what we term LPF-FD and LPF-DV, respectively; and 2) adaptively setting LPF-DV thresholds within each run separately to accommodate differences in baseline and central tendency (Afyouni and Nichols, 2018), by fitting a generalized extreme value (GEV) distribution to each run’s LPF-DV values and rejecting the upper tail (GEV-DV censoring, demonstrated in **Figure S2**). Although prior work has proposed the use of notch-filtered FD as discussed in *Section 1.1* above, we expected *a priori* that low-pass filtering would outperform these methods when a standard band-pass filter is applied to RSFC timeseries data (e.g., 0.009 – 0.08 Hz or 0.01 – 0.1 Hz), because the high-frequency signals retained by a notch filter would be removed from RSFC data by the bandpass filter. Nonetheless, we evaluate notch filtering approaches to FD here as well.

Next, we aimed to quantify improvements in RSFC data quality using traditional RSFC data quality metrics (hereafter, DQMs) that have been employed throughout the RSFC literature (Burgess et al., 2016; Ciric et al., 2017; Muschelli et al., 2014; Parkes et al., 2018; Power et al., 2012; Power et al., 2014; Power et al., 2015; Pruim et al., 2015; Satterthwaite et al., 2013; Satterthwaite et al., 2012; Van Dijk et al., 2012; Yan et al., 2013) in the Human Connectome Project 500 Subjects Release (HCP500) dataset (Smith et al., 2013; Van Essen et al., 2013). As it was unclear which volume censoring threshold is “optimal” for any given dataset, we calculated these DQMs across the full range of potential thresholds for each method, intending to use them to first determine optimal volume censoring parameters for each method, and then comparing across methods.

However, this investigation revealed substantive issues with nearly all evaluated DQMs that precludes their use as criteria, either alone or in combination, for establishing optimal volume censoring parameters within methods, or as clearly interpretable arbiters between methods, mirroring concerns that have been raised previously (Glasser et al., 2018). These issues are particularly pronounced in DQMs that depend on associations between subject-level QC metrics (such as mean or median FD; hereafter, SQMs), and the observed magnitude of RSFC correlations (e.g., QC-FC correlations), which we further demonstrate are likely due to confounding third-variables influencing both SQMs and true differences in brain connectivity across research participants. These issues, which we detail in section 3.1, led us to develop a novel DQM intended for comparing between censoring methods that quantifies a common graphical approach to quality assessment used in prior work (Power et al., 2012; Power et al., 2014), which we call mean absolute change in resting state functional connectivity (MAC-RSFC).

Next, we developed and evaluated a quantitative, empirical, method for determining the optimal value of volume censoring parameters for removing motion artifacts from RSFC data by optimizing the trade-off between the reduction motion-induced bias and loss of power that result from data removal due to censoring, which we call ΔMSE-RSFC. This approach uses a bias-variance decomposition of mean squared error (MSE) to estimate the total error in sample mean RSFC correlations. We then applied a global multiparameter optimization procedure to this measure to determine optimal parameters for simultaneous LPF-FD- and GEV-DV-based censoring in the HCP500 dataset by minimizing ΔMSE-RSFC. In addition, we present a MATLAB tool for applying a computationally simplified version of this method, such that investigators can readily identify near-optimal censoring thresholds for their own datasets, and demonstrate its use in an independent dataset from our own laboratory.

### 1.3. Summary

This manuscript describes our efforts to benchmark volume censoring methods and to optimize volume censoring parameters (i.e., censoring thresholds). First, we evaluated existing data quality metrics (DQMs) and found substantial concerns with their use in the optimization of volume censoring methods. Second, we developed novel DQMs that ameliorate some of these concerns: 1) MAC-RSFC, for comparing between censoring methods; and 2) ΔMSE-RSFC, for optimizing censoring parameters within censoring methods. We subsequently evaluate these metrics over a wide range of censoring parameter values (i.e., over the full range of “strictness” for motion denoising). Finally, we present a software tool that will allow investigators to extend these results to other datasets and evaluate this approach in another, independently acquired dataset. These findings, and their associated figures, are summarized in **Table 1**, which readers may wish to use as a roadmap to the many analyses reported herein.

**Table 1.**
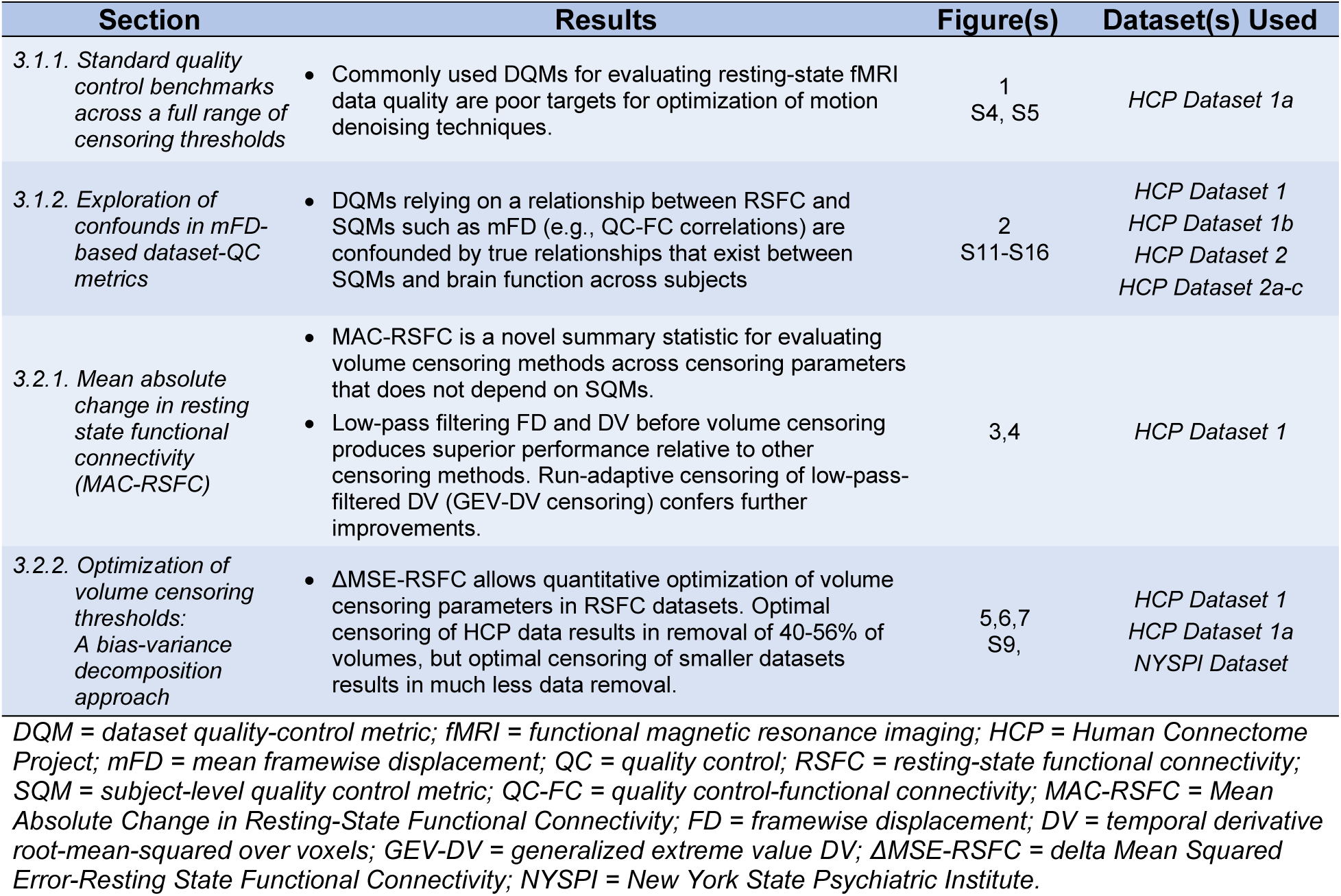
Summary of major findings and associated figures and datasets used in analyses.

## 2. Materials and Methods

### 2.1. General methods

#### 2.1.1. Functional magnetic resonance imaging datasets

##### 2.1.1.1. Human Connectome Project (HCP) datasets

In our primary analyses, we employed minimally preprocessed resting-state data (Glasser et al., 2013) from the Human Connectome Project (Smith et al., 2013; Van Essen et al., 2013) 500 Subjects Release (HCP500; n = 501), with 4 runs of resting-state fMRI data per subject (3 subjects had 3 runs, and 17 had 2 runs). Each run is composed of a maximum of 1200 volumes (14 minutes 24 seconds) of data (mean = 1174.12 volumes), collected over two sessions. We also produced two subsets of *HCP Dataset 1*, denoted *HCP Dataset 1a* and *HCP Dataset 1b*, to support specific analyses detailed in forthcoming sections. In *HCP Dataset 1a* (n = 475; shown in *Section 3.1.1* and *Section 3.2.2.1*), we removed high-motion subjects who were dropped from analyses prior to reaching 50% of volumes censored using LPF-FD censoring, to control for subject removal when evaluating DQMs. In *HCP Dataset 1b* (n = 314; shown in *Section 3.1.2.1*), we removed 187 subjects with a family history of neurological or psychiatric disease (FH+), or evidence of substance use during the course of the study (SU+). All HCP data were acquired with 2 mm^3^ isotropic voxels, 720 ms TR, and multiband acceleration factor of 8, as detailed elsewhere (Smith et al., 2013; Ugurbil et al., 2013).

*HCP Dataset 2* (n = 403, including 186 subjects from *Dataset 1*; shown in *Supplementary Section S2.1*) is a subset of minimally preprocessed resting-state data from the HCP 1200 Subjects Release (HCP1200), comprising only unrelated subjects with 4 full (1200 volume) runs of resting-state fMRI data, with no runs flagged with QC issue A (anatomical anomalies), B (segmentation and surface QC), and C (some data acquired during periods of head coil instability), and only the first sibling from each family (based on numerical subject identifier), to eliminate non-independence due to familial relationships. This dataset includes 161 FH+ subjects and 242 FH- subjects. *HCP Dataset 2* was used to produce three subsets (*Datasets 2a, 2b, and 2c*) consisting of 161 FH+ and 161 FH- (in each, total n = 322; all shown in Supplementary *Section S2.1*), selected by optimally matching FH- to FH+ subjects using three different procedures described in *Supplementary Section S1.4.* We employed the full HCP 1200 Subjects release for *HCP Dataset 2* because the HCP 500 Subjects Release would not have provided a sufficient number of unrelated FH+ and FH- subjects.

Additionally, as described in *Supplementary Section S1.1.1*, we further generated two additional subsets of the minimally preprocessed HCP500 dataset, comprising: 1) both resting-state and working memory task data (Barch et al., 2013) from 330 subjects with a complete set of working memory task and resting-state data (*Supplementary Section S1.1.1.2*), and 2) resting-state data from 478 subjects with 4 complete runs of resting-state data (*Supplementary Section S1.1.1.3*).

##### 2.1.1.2. New York State Psychiatric Institute (NYSPI) Dataset

To validate ΔMSE-RSFC-based optimization of volume censoring parameters (see *section 3.2.2.2*), we employed a sample of resting state data acquired from 26 healthy participants (8 female, 18 male, mean age = 33.58) using a General Electric (GE; Boston, MA) MR750 3T scanner (multiband factor 6, TR = 850 ms, voxel size = 2 mm isotropic) scanner at the New York State Psychiatric Institute (NYSPI; New York, NY), over 4 runs of 15 minutes each. Further details of this dataset are provided in the *Supplementary Section S1.1.2,* with demographic information shown in **Table S1**.

All procedures associated with the collection of the NYSPI dataset were approved by the New York State Psychiatric Institute Institutional Review Board. Written informed consent was obtained from each participant prior to their entering into the study. Participants were compensated monetarily for their participation.

#### 2.1.2. Resting-state functional connectivity preprocessing

Data from NYSPI were first preprocessed using the Human Connectome Project (HCP) Minimal Preprocessing Pipelines (Glasser et al., 2013) version 3.20.0 (https://github.com/Washington-University/HCPpipelines). All Human Connectome Project resting-state data were downloaded in minimally-preprocessed form from the “Resting State fMRI 1 Preprocessed” and “Resting State fMRI 2 Preprocessed” packages for each subject on db.humanconnectome.org (Hodge et al., 2016). Note that these data do not include the ICA-FIX cleanup procedure implemented by the HCP consortium. Resting-state blood-oxygen-level-dependent (BOLD) data were obtained from the files, *rfMRI_REST1_LR.nii.gz*, *rfMRI_REST1_RL.nii.gz, rfMRI_REST2_LR.nii.gz,* and *rfMRI_REST2_RL.nii.gz*, and motion parameters were obtained from the file, *Movement_Regressors.txt* corresponding to each run.

In all HCP datasets, we discarded the first 10 volumes of each run to allow for magnetic resonance signal equilibration prior to any further preprocessing. Data from NYSPI had two calibration volumes discarded prior to any preprocessing (these images are attached to the functional run for this GE sequence, but not for HCP datasets or other Siemens multiband sequences), and then the first 7 volumes were discarded after minimal preprocessing for magnetic resonance signal equilibration.

In all datasets, signals from 264 regions of interest (ROIs) were obtained as the average fMRI timeseries within 10mm diameter spheres from prior work (Power et al., 2011). Nuisance signals from white matter, cerebrospinal fluid, and global signal were obtained following published methods (Power et al., 2014), described in detail in *Section 2.1.2.1*. ROI signals were mode 1000 normalized (Power et al., 2012), demeaned, detrended, and 0.009 – 0.08 Hz band-pass filtered using a second-order zero-phase Butterworth filter (note that detrending and band-pass filtering ROI timeseries after spatial averaging over voxels within each ROI mask is equivalent to detrending and band-pass filtering voxel timeseries prior to spatial averaging). The first and last 30 volumes of each run were then discarded due to filter edge effects.

##### 2.1.2.1. Calculation of nuisance signals

Global signal (GS) was calculated as the mean signal across in-brain voxels, determined from the *brainmask_fs.2.nii.gz* files for each subject. Nuisance signals from white matter and cerebrospinal fluid were calculated as the average signal in all compartment voxels remaining after an iterative erosion procedure, in which masks were eroded up to four times as long as at least one voxel remained in the mask following erosion. GS, white matter, and cerebro-spinal fluid signals were all calculated prior to the removal of other nuisance signals (see *Section 2.1.2.2*).

##### 2.1.2.2. Calculation of resting-state functional connectivity (RSFC) correlations

RSFC correlations were calculated as partial correlations between each pairwise ROI, controlling for nuisance parameters (but not controlling for the timeseries of any other ROI), after volume censoring (Power et al., 2012; Power et al., 2014; Smyser et al., 2011). In cases where multiple volume censoring methods are used together (e.g., when using LPF-FD and GEV-DV censoring in tandem), volumes exceeding either threshold were censored.

Although the use of partial correlations (as opposed to residualizing timeseries data against nuisance parameters using regression) is somewhat uncommon, it ensures that the effects of nuisance parameters on RSFC correlations are always orthogonal, which protects against the accidental reintroduction of nuisance signals when nuisance regressions are conducted in a stepwise manner (Lindquist et al., 2019). Because the calculation of partial correlations and sequential linear regressions are all linear operations, our approach should not produce different results from sequential regressions, if the sequential regressions are conducted correctly (i.e., in a way that avoids reintroducing signals removed in prior steps; see Lindquist et al., 2019). Finally, this approach also has the advantage of requiring substantially less disk space, as only the ROI timeseries (*Section 2.1.2*) and nuisance signals (see *Section 2.1.2.1* and the following paragraph) need to be stored, rather than a fully processed whole-brain volume.

The following nuisance regressors were controlled for when calculating RSFC correlations: band-pass filtered MPs (using the same filter as was applied to the RSFC timeseries) and the filtered MPs squared, the derivatives of the band-pass filtered MPs and the squares of those derivatives, the white matter signal and its derivative, and the cerebro-spinal fluid signal and its derivative. All derivatives were calculated by backwards differences. Analyses both with and without GSR (including its first derivative) were conducted and are presented side by side throughout. All RSFC correlations were Fisher’s *r*-to-*Z* transformed immediately after their calculation, prior to their use in any further computations or analyses, and are reported in this form throughout this manuscript. In all cases, (Z-transformed) RSFC correlations for each subject were calculated separately over runs and averaged together.

#### 2.1.3 Volume censoring methods

##### 2.1.3.1 Standard censoring

For standard (FD and DV) volume censoring, volumes were identified for removal when their FD or DV exceeded a threshold set for each metric. FD was calculated as the sum of the absolute value of all six motion parameters, following conversion of rotation parameters to the translation of a point on the circumference of a 50 mm radius sphere (Power et al., 2012); DV was calculated as the root-mean-square (over voxels) of the first derivative (by backwards differences) of the timeseries across all brain voxels (Power et al., 2012; Smyser et al., 2011) after mode 1000 normalization, without band-pass filtering or removal of nuisance regressors (which are conducted later in the pipeline, prior to calculating RSFC correlations in a partial correlation model with nuisance regressors). Calculation of standard DV values was carried out following volume smoothing with a 4 mm FWHM gaussian kernel in SPM 12, to produce values closer to those reported for prior datasets (Power et al., 2012; Power et al., 2014), as unsmoothed data produces much higher values of DV. Smoothed data were not used for any other purpose, including calculation of LPF-DV values. Prior to band-pass filtering of timeseries, linear interpolation was used to replace censored time series data before discarding these time points from analysis. Runs for which less than 2 minutes of uncensored data (167 volumes) remained after censoring were excluded from analysis. Subjects were retained for analysis provided they had at least 1 run remaining after censoring.

##### 2.1.3.2. Low pass filtered framewise displacement (LPF-FD) and DV (LPF-DV)

LPF-FD was obtained by calculating FD as normal, but on MPs that were first low-pass filtered at 0.2 Hz with a second-order zero-phase Butterworth filter. LPF-DV was calculated by applying the same filter to voxel timeseries data prior to calculation of DV. The aggressiveness of LPF-FD and LPF-DV was set by selecting threshold LPF-FD and LPF-DV values, denoted *Φ_F_* and *Φ_D_* respectively.

##### 2.1.3.3. Band-stop (notch) filtered framewise displacement (Notch-FD) and DV (Notch-DV)

Band-stop (notch) filtered framewise displacement (Notch-FD) and DV (Notch-DV) were calculated identically to LPF-FD and LPF-DV, respectively, except using a second-order zero-phase band-stop filter in place of a low-pass filter, as has been previously developed (Fair et al., 2020; Power et al., 2019a). Two versions of Notch-FD and Notch-DV were evaluated: one with a 0.31 Hz – 0.43 Hz band-stopband as suggested by Fair and associates (Fair et al., 2020), and another with a 0.2 Hz – 0.5 Hz stopband as suggested by Power and associates (Power et al., 2019a).

##### 2.1.3.4. Adaptive DV censoring using generalized extreme value fitting (GEV-DV)

We also examined using run-specific LPF-DV censoring thresholds that adapt to the amount of motion in each run by fitting a generalized extreme value (GEV) distribution (Prescott and Walden, 1980) to the LPF-DV values within each run by maximum likelihood estimation (using the MATLAB function *fitdist*), and setting the threshold LPF-DV value *Φ_D_* separately within each run such that the empirical cumulative distribution function at *Φ_D_* is equal to 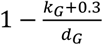 (i.e., the area under the curve of the GEV to the right of the cutoff is equal to 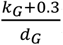), where *k_G_* is the shape parameter obtained from the GEV fit, and *d_G_* is a free parameter. The shape parameter *k_G_* is greater in runs containing more extreme DV values (i.e., a thicker right tail), causing a greater proportion of the data to be excluded when more data has high LPF-DV values relative to the central tendency for that run. The +0.3 in the numerator ensures that the numerator cannot take on a negative value, while the free parameter *d_G_* is a tunable free parameter that sets the aggressiveness of volume censoring across the dataset (placing the free parameter in the denominator instead of as a multiplier preserves the intuition that a higher value results in a more lenient censoring threshold, as with other censoring parameters). We term this censoring method run-adaptive LPF-DV censoring, or GEV-DV censoring.

### 2.2. Evaluation of commonly used data-quality metrics (DQMs)

#### 2.2.1. Standard quality control benchmarks across a full range of censoring thresholds

The following SQMs and DQMs were calculated on *HCP Dataset 1* using published techniques: QC-FC (Quality Control – Functional Connectivity; Power et al., 2012; Power et al., 2014; Power et al., 2015; Satterthwaite et al., 2013; Satterthwaite et al., 2012; Van Dijk et al., 2012; Yan et al., 2013); median absolute QC-FC (Ciric et al., 2017); QC-FC distance correlation (the correlation between QC-FC and ROI pair distance; Ciric et al., 2017; Muschelli et al., 2014; Parkes et al., 2018; Power et al., 2012; Satterthwaite et al., 2013); QC-FC null rejection rate (proportion of significant QC-FC correlations, after false discovery rate correction; Ciric et al., 2017); and high-low null rejection rate (proportion of significantly different RSFC correlations between high- and low-motion terciles of the sample; Burgess et al., 2016; Power et al., 2014; Pruim et al., 2015; Satterthwaite et al., 2013; Van Dijk et al., 2012). Additionally, the mean and distance-dependence of QC-FC correlations and RSFC correlations were evaluated using a two-step generalized linear model (GLM) method (Burgess et al., 2016). The precise details of the computation of each of these measures is presented in *Supplementary Section S1.2.* Briefly, QC-FC (which, in addition to being a DQM itself, is also used in the calculation of many other DQMs), refers to the correlation between an SQM such as the mean FD value for a participant and their RSFC correlations.

#### 2.2.2. Exploration of confounds in mFD-based dataset-QC metrics

##### 2.2.2.1. Comparison of high- to low-motion participants after censoring

We determined the number of significant differences in RSFC correlations between high- and low-motion participants in *HCP Dataset 1*, following prior work (Parkes et al., 2018; Power et al., 2014; Pruim et al., 2015; Satterthwaite et al., 2013; Van Dijk et al., 2012), by splitting the sample into terciles based on median FD for each participant. We also removed participants with a family history of any psychiatric or neurological condition (FH+), or who engaged in drug use or had elevated blood-alcohol content (BAC) over the course of the study (SU+), which resulted in exclusion of 154 FH+ and 32 SU+ participants, respectively, to produce *HCP Dataset 1b*, and repeated this process.

To determine whether removal of these participants for *HCP Dataset 1b* resulted in a greater-than-expected reduction in the number of differences in RSFC correlations, which would indicate a confounding effect of FH and SU status on the relationship between FD and RSFC correlations, we also generated 1000 Monte Carlo resamplings of the dataset that excluded an equal number of subjects, but by excluding subjects who were instead FH- and SU-, and who still had approximately the same average motion as the FH+ and SU+ subjects, as detailed in *Supplementary Section S1.3*.

Significant differences between high- and low-motion participants (respectively, the upper and lower tercile of the median FD for each participant across all volumes in all runs) were then determined for each ROI pair across a range of uncorrected significance thresholds, from α = 0.0001 to α = 0.1, in increments of 10^-5^. This was done for the full dataset, for a dataset excluding all family history or substance use positive participants, and for each of the 1000 Monte Carlo samples.

##### 2.2.2.2. Evaluation of trait confounds in DQMs using datasets balanced by subject motion

In order to address the possibility that the Monte Carlo procedure described above in *Section 2.2.2.1* and *Supplementary Section S1.3* failed to match the *distribution* of motion within subjects, given that it focused exclusively on average motion, we also produced three datasets in which FH+ and FH- participants were matched on a one-to-one basis based on the temporal dynamics and overall motion characteristics across all of their data. We divided each of three datasets (see *Supplementary Section S1.4* for details on the construction of these datasets) into terciles based on mean FD to create high- and low-motion groups, similar to the procedure described in *Section 2.2.2.1*. For each of these datasets, we calculated the number of significantly different RSFC correlations between high- and low-motion terciles after removing the 161 FH+ subjects, and again after instead removing the 161 FH- subjects matched by motion. We then evaluated whether the number of significant high-low differences obtained after removing each subset differed using Fisher’s exact test.

### 2.3. Development and evaluation of volume censoring with novel dataset-quality metrics (DQMs)

#### 2.1.1. Mean absolute change in resting state functional connectivity (MAC-RSFC)

Mean Absolute Change in Resting State Functional Connectivity (MAC-RSFC) is obtained by first calculating the mean within-subject RSFC correlations in all ROI pairs after targeted volume censoring, and then subtracting the mean RSFC correlations resulting from random censoring (averaged over 10 random censoring vectors), to obtain Δ*Z_i,k_* for the *i*th subject and *k*th ROI pair.

That is, for subject *i*, run *j*, ROI pair *k*, and random permutation vector *p*:

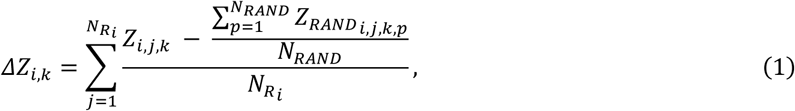

where *Z_i,j,k_* is the RSFC correlation (Z-transformed Pearson’s product moment correlation) for subject *i,* run *j,* and ROI pair *k,* after targeted volume censoring. 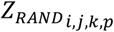 is the RSFC correlation for subject *i,* run *j*, and ROI pair *k* after volume censoring using random permutation vector *p*, 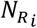 is the number of runs that exist after censoring for subject *i*, and *N_RAND_* is the number of random permutations used per run (here, 10). MAC-RSFC is calculated for the full sample as:

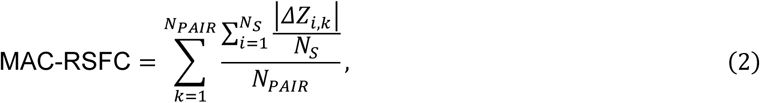

in which *N_S_* is the number of subjects in the sample after volume censoring and N_PAIR_ is the number of ROI pairs used for analysis (here, 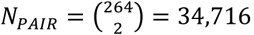).

95% confidence intervals for MAC-RSFC were approximated for each parameter value using the bias-corrected and accelerated bootstrap with 10,000 bootstrap samples (Efron and Tibshirani, 1993). Note that MAC-RSFC has a nonzero null expectation, making it unsuitable for evaluating whether censoring using any given method has a significant effect on RSFC correlations. However, comparisons between two or more methods for any given quantity of volumes removed by censoring do have a null expectation of zero (provided neither method has a differential effect on the removal of true neural signals; see *Section 3.2.3* for an in-depth discussion of this issue). That is, in the absence of any true between-method (or between-parameter) differences in the impact of targeting volumes for removal on RSFC estimates, MAC-RSFC would be expected to be identical across methods. Note further that, while *ΔMSE-RSFC* (see *Section 2.3.2.2*) decomposes the effects of volume censoring on sample RSFC estimates into bias and variance components, MAC-RSFC measures both in aggregate.

#### 2.3.2. Optimization of volume censoring thresholds: a bias-variance decomposition approach

##### 2.3.2.1. Quantitative methods for selecting censoring thresholds

To our knowledge, the only formalized procedure for quantitatively selecting a censoring threshold in the literature involves calculating the correlation between RSFC estimates from a sliding window and the lowest-motion sliding windows in a given subject, and binning these estimates by the maximum FD value observed in the window (Fair et al., 2020; Power et al., 2014). The correlation of each window to the reference (low-motion) windows is then compared to randomly-ordered (permuted) data, and a threshold is selected such that sliding windows with higher FD values than the threshold show a significantly lower correlation to the reference data than is expected by chance.

Unfortunately, this approach has multiple drawbacks that make it unsuitable for selecting an optimal censoring threshold, either within a single study or in the literature more broadly. First, although threshold selection is based on a null-hypothesis statistical test (NHST), it is a misapplication of NHST to an optimization problem. There is no *a priori* reason why P = 0.05 should lead to an optimal censoring parameter, as compared to any other P value. Indeed, one could argue that the top 5^th^ percentile, rather than the bottom, should be used (i.e., P = 0.95, rather than P = 0.05, in the original 1-tailed formulation of the test), which would reflect statistically significant evidence that a given time-window is *more similar* to the most motion-free data in a single subject than is an average (and thus more heavily motion-contaminated) time-window. Ultimately, any P value threshold could be used to select a cutoff because the goal of setting a censoring threshold has no direct relationship to any NHST in this context. Thus, while this method is quantitative, the choice of cutoff remains arbitrary. Moreover, this approach has the additional drawback of identifying different thresholds for different subjects, and will paradoxically select more lenient thresholds for higher motion subjects (because the null reference distribution generated from random timepoints should have more contamination from high-motion data, the null distribution in these subjects should have less similarity to the lowest-motion data and will thus set the P < 0.05 cutoff at a higher FD). Indeed, work using this approach in the ABCD dataset (see Fair et al., 2020, Figure 7B) suggests an FD cutoff anywhere between approximately 0.05 and 0.3 mm depending on whether one desires a censoring threshold below the P < 0.05 threshold for *all* subjects or a threshold below this threshold in only the highest-motion subject (or somewhere in between).

Thus, we instead developed a novel method of determining optimal volume censoring thresholds by balancing the tradeoff between reducing the impact of motion and other artifacts on RSFC and minimizing the loss of temporal degrees of freedom (and thus power) that occurs when data is discarded. Our goal was to optimize the ability of investigators to resolve true sample-level mean RSFC correlations by simultaneously minimizing both a) motion-induced bias, defined as the change in sample mean RSFC correlations due to motion-targeted volume censoring; and b) the increase in RSFC confidence interval widths resulting from loss of high-motion volumes, runs, and subjects.

To this end, we developed a novel DQM, described in full in *Section 2.3.2.2* through *Section 2.3.2.4*, which we term ΔMSE-RSFC. ΔMSE-RSFC measures overall improvement in RSFC estimates resulting from motion denoising by employing a mean-squared error (MSE) calculation from a bias-variance decomposition (i.e., an MSE calculation that includes bias, in addition to variance), divided by the sample size (number of subjects remaining after volume censoring). MSE is divided by sample size because we are attempting to optimize the ability to identify the true values of RSFC correlations at the group level, with minimum bias and error. Because the error of this estimate will scale proportionally to the inverse root of the sample size, the square of the error will increase proportionally to the inverse of N. Thus, we include N in the denominator of our optimization target, so that censoring thresholds will be penalized for causing the removal of subjects from the dataset, as doing so increases the error of group level estimates.

##### 2.3.2.2. Calculation of ΔMSE-RSFC

We developed *ΔMSE-RSFC* as a quantitative benchmark for volume censoring performance that could be used as an optimization target to determine optimal free parameter values for each volume censoring method (i.e., optimal LPF-FD *Φ_F_* and GEV-DV *d_G_*). *ΔMSE-RSFC* was calculated in each ROI pair *k* and averaged over all ROI pairs, i.e.,

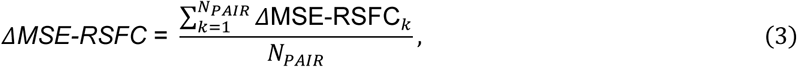

where N_PAIR_ is the number of ROI pairs (34,716), and

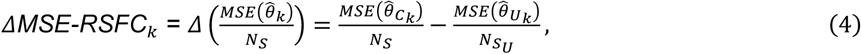

where 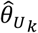 is the estimated RSFC correlation in ROI pair *k* from uncensored data, 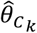 is the same estimator for volume censored data, and *N_S_U__* is the number of subjects in the dataset in the uncensored dataset. 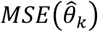 is the mean squared error in the RSFC correlation estimate for ROI pair *k*, accounting for both the variance (observed across subjects) and bias produced by motion artifact:

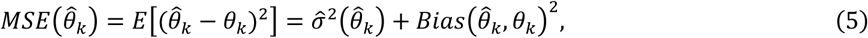

in which *θ_k_* is the true population mean RSFC correlation for a single ROI pair *k* and 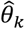 is its estimator, 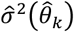 is the observed between-subjects variance in 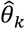 and 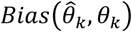 is the bias in 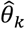, here defined as the total motion-induced bias that is removable by volume censoring (for general details on the bias-variance decomposition of *MSE,* see Wackerly et al., 2008). Thus, **Equation 4** can also be written as:

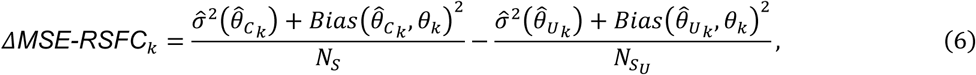

where 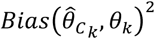 is the motion-induced bias in RSFC correlation *k* after censoring; 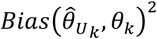 is the bias in RSFC correlation *k* before censoring, estimated as described in *Section 2.3.2.4*.

**Equation 6** may be used to calculate ΔMSE-RSFC in ROI pair *k* as a function of volume censoring parameter values, as shown in **Figure 5.** 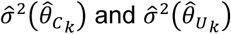 may be computed directly as variance across subjects in ROI pair correlation *k* after censoring and before censoring, respectively; *N_S_* is determined by observing the number of subjects remaining in the dataset for analysis after censoring, *N_S_U__* the number of subjects in the dataset before censoring. 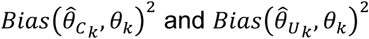 are respectively estimated as described in *Section 2.3.2.3* and *Section 2.3.2.4*, below.

##### 2.3.2.3. Estimation of bias after censoring

As it is not possible to measure either 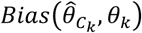 or its square, 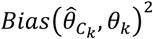, directly, we instead first estimated the change in bias due to volume censoring, 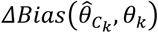, in each ROI pair by measuring the mean sample-wide magnitude of the change in its RSFC correlation due to volume censoring and taking the additive inverse, i.e.,

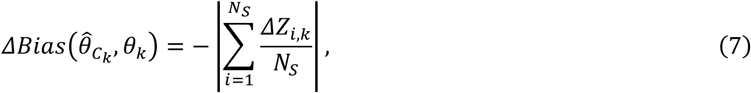

where Δ*Z_i,k_* is the change in in observed RSFC correlation *k* for subject *i* due to targeted censoring, over and above the change due to random censoring, as defined in **Equation 1** (see *Section 2.3.1*).

We can write the bias present in ROI pair correlation *k* as the bias present in uncensored data, plus the change in bias produced by volume censoring:

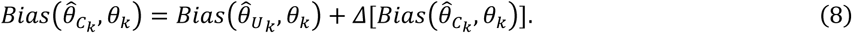

It follows from **Equation 8** that the square of the bias present after censoring in ROI pair correlation *k*, 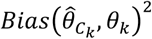, can be written as the squared bias present in uncensored data, plus the change in squared bias produced by volume censoring:

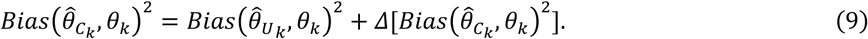

The change in squared bias, 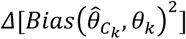, in **Equation 9**, is distinct from the square of the change in bias (which would be instead denoted as 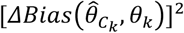. Due to this distinction, 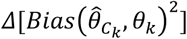] may not be obtained simply by squaring the estimate of 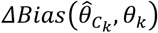 obtained from **Equation 7** (as exponentiation is a nonlinear operation).

The change in squared bias due to volume censoring in ROI pair *k* may be expressed alternatively as the difference therein between censored and uncensored data, which also follows from **Equation 9**:

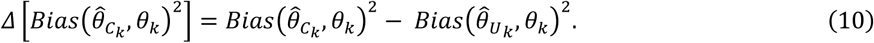

Next, using **Equation 8**, we can then rewrite **Equation 10** as follows:

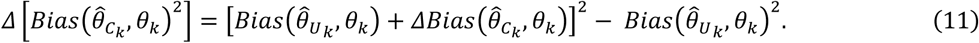

The artifact-induced bias remaining in the estimate for ROI pair *k* after censoring may now be calculated from estimates of bias in the uncensored data and the observed change in bias due to censoring (Equation 11):

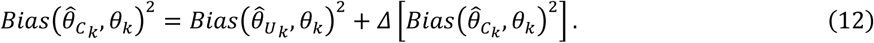

**Equation 7** may be used to calculate 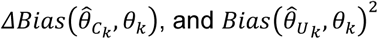 is calculated as described in *Section 2.3.2.4*, below. The resulting estimate of 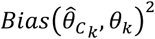 from **Equation 12** can then be used in **Equation 6** to calculate *ΔMSE-RSFC*_*k*_ for each ROI pair *k*.

##### 2.3.2.4. Estimation of total bias in uncensored data

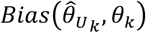 is the total sample-wide motion-induced bias in RSFC correlations that is removable by volume censoring, and is necessarily unknown. However, we estimate this value for each ROI pair by measuring the additive inverse of the change in mean RSFC correlations due to volume censoring, 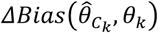, as a function of percent of frames removed (that is, estimation occurs after re-parameterizing censoring thresholds as percent frames removed and resampling to a resolution of 0.01% using linear interpolation). We then estimate the slope using robust regression (bisquare weight function, tuning constant of 4.685, and an intercept of 0), and extrapolating to estimate the value at 100% frames removed. This was done to provide a reasonable estimate of total bias that is robust to the instability in RSFC correlation estimates as the number of frames removed begins to approach 100%. As a demonstration, this method is shown for the average RSFC correlation across all ROI pairs, using LPF-FD based censoring, without GSR, in **Figure S3.** The estimate of 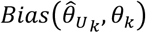 that is obtained through this procedure allows for the calculation of 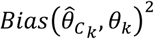 using **Equation 12**, which is in turn used in the calculation of *ΔMSE-RSFC_k_* using **Equation 6.**

##### 2.3.2.5. Volume censoring optimization

When volume censoring parameters for LPF-FD and GEV-DV were optimized separately (i.e., when using one censoring method at a time), *Φ_F_* and *d_G_* were respectively varied such that the percentage of volumes censored across the HCP500 data set varies from 0% to 100%. For each censoring threshold, ΔMSE-RSFC was calculated as described in *Section 2.3.2.2*. After censoring, the minimum value of ΔMSE-RSFC was obtained in the range of 0% to 80% volumes censored (to exclude areas of instability caused by excessive removal of data), and the corresponding volume censoring parameter was taken as the optimal value for single-method volume censoring.

##### 2.3.2.6. Optimization of volume censoring parameters in arbitrary datasets

As previously stated, volume censoring is routinely performed using criteria from both estimated framewise motion and signal fluctuation in tandem, respectively implemented in this work using LPF-FD and GEV-DV. Optimizing this dual censoring technique using ΔMSE-RSFC as an objective function requires optimizing both censoring parameters (*Φ_F_* and *d_G_*) simultaneously. As described in *Supplementary Section S1.5,* we successfully performed a two-dimensional global optimization procedure in the HCP500 dataset. However, direct application of this procedure in arbitrary datasets may not be practical in all cases due to two major limitations. First, finely searching a 2-dimensional parameter space as required to avoid local minima is computationally expensive, even when using modern stochastic global optimization techniques. Second, datasets with less data than the HCP500 would be expected to have a greater degree of instability, which would impair 2-dimensional optimization procedures considerably more than those optimizing parameters in one dimension only.

As a result, we developed a one-dimensional optimization procedure for combined LPF-FD and GEV-DV volume censoring simultaneously, fixed in the ratio of LPF-FD *Φ_F_* to GEV-DV *d_G_* found to be optimal in the HCP500 dataset (see *section 3.2.2*). This provides the advantage of reducing the order of time complexity of the computations required for ΔMSE-RSFC optimization and reducing the number of free parameters to be optimized, increasing robustness to noise in light of relatively low data quantity in the case of smaller datasets. We employed this technique to obtain optimal volume censoring parameters for combined LPF-FD and GEV-DV volume censoring in the *NYSPI Dataset* (n = 26; see *section 2.1.1.2*). We additionally have released software allowing investigators to implement this procedure in their own RSFC datasets, as described in *Section 5.3*.

#### 2.3.3 Positive controls for RSFC volume censoring

We additionally conducted two analyses aimed at determining whether the censoring methods suggested by the analyses described above may remove (desirable) true neural signal from RSFC, which is a potential pitfall of the MAC-RSFC and ΔMSE-RSFC methods described above (see *Section 3.2.3* for discussion). These analyses are described in detail in *Supplementary Section S1.6*.

## 3. Results and Discussion

### 3.1. Evaluation of commonly used data-quality metrics (DQMs)

#### 3.1.1. Standard quality control benchmarks across a full range of censoring thresholds

Our first step was to calculate widely used QC metrics for evaluating the success of motion denoising of RSFC datasets (i.e., DQMs) across a comprehensive range of potential censoring thresholds, ranging from no censoring to thresholds that approach removal of all data in the dataset. Although extremely computationally intensive, we view this as a critical step in establishing which DQMs are best suited for determining the success or failure of volume censoring techniques, as well as for optimizing such techniques.

We show several commonly-used DQMs in **Figure 1**, calculated after either standard FD or LPF-FD based volume censoring, and presented as a function of the percentage of volumes in the entire dataset that are removed as a result of censoring (to allow direct comparison between methods despite differing threshold values), with identified “optima” in **Table S2**. Results for DV-based measures are shown in **Figure S4** (Standard DV and LPF-DV) and **Figure S5** (GEV-DV), with “optima” shown in **Table S3** and **Table S4**. Taken together, the results shown in **Figure 1** raise serious questions as to whether several of these DQMs are appropriate benchmarks for evaluating the denoising of RSFC datasets. For example, each of: a) the median absolute value of all QC-FC correlations (**Figure 1A;** see Ciric et al., 2017), b) the proportion of ROI pairs with statistically significant QC-FC correlations (**Figure 1B;** see Ciric et al., 2017), and c) the proportion of ROI pairs showing group differences between high- and low-motion terciles of the dataset (high-low null rejection rate; **Figure 1D;** see Burgess et al., 2016; Power et al., 2014; Pruim et al., 2015; Satterthwaite et al., 2013; Van Dijk et al., 2012) show nearly immediate increases, rather than the expected decreases, as censoring becomes more aggressive for both standard FD and LPF-FD when GSR is employed. A similar effect is seen with standard FD without GSR, but after a clearer “initial dip.” While this dip may suggest an optima, the properties of the data that drive the dramatic rise in these metrics after this optima are unclear. At face value, these results suggest that with GSR virtually no censoring need be performed, and that without GSR, LPF-FD achieves better data quality than standard FD, but at the cost of censoring 4-5 times as much data. However, as we argue in more detail below, our view is that these findings instead reflect fundamental issues with this set of DQMs, and that alternative DQMs should be employed instead.

**Figure 1.**
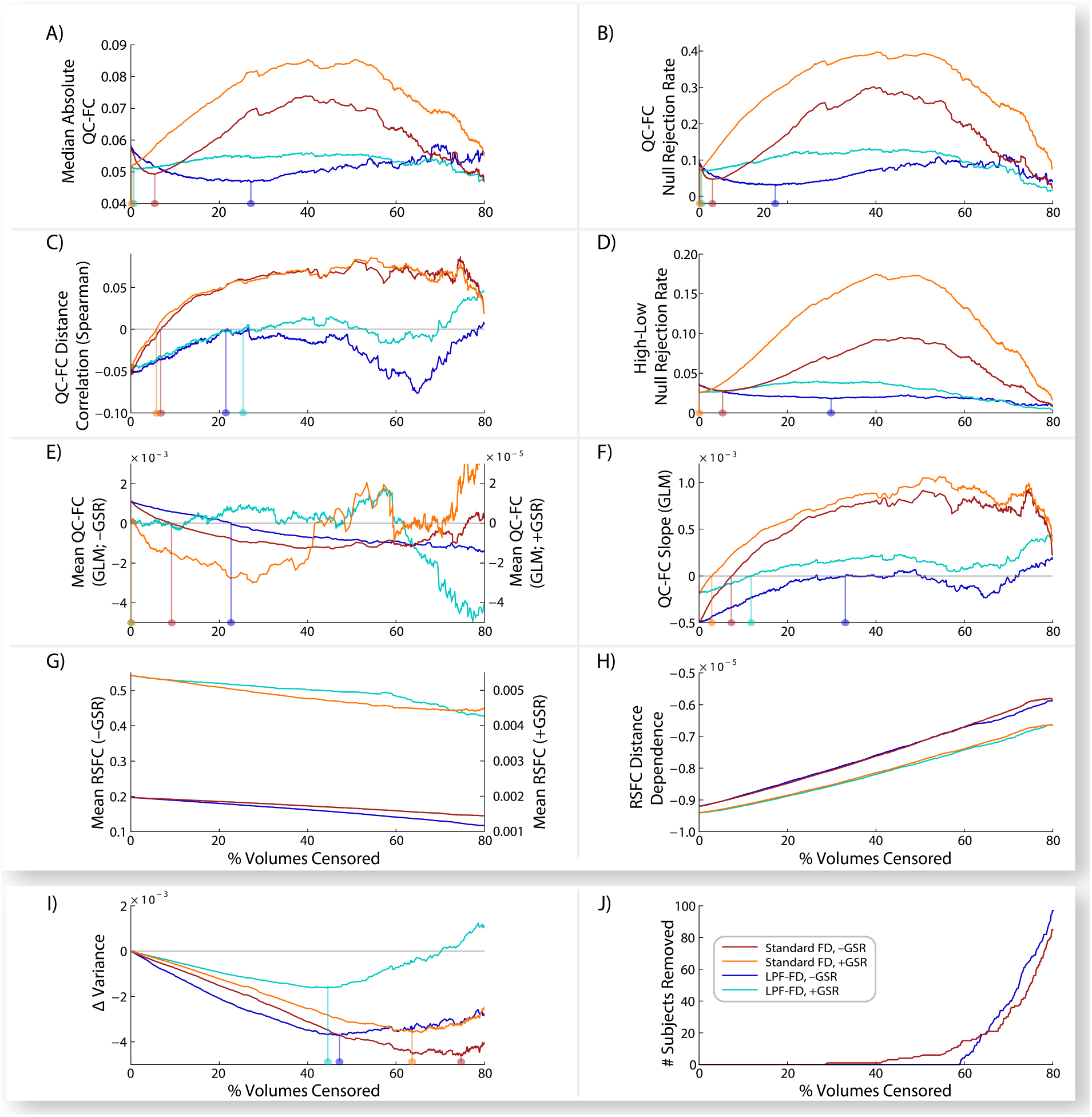
Commonly used dataset quality control metrics (DQMs) as a function of the percent volumes censored in a subset of the HCP500 dataset (*HCP Dataset 1a*; n = 475). DQMs calculated after volume censoring using either standard framewise displacement (FD) or low-pass-filtered FD (LPF-FD). **A)** Median absolute QC-FC across ROI pairs, **B)** proportion of ROI pairs with significant QC-FC correlations (null rejection rate), **C)** correlation of QC-FC with the distance between ROI pairs, and **D)** proportion of ROI pairs showing significant differences between high-motion and low-motion participants. Also shown are DQMs calculated using a two-step generalized linear model (GLM), including: **E)** mean QC-FC correlation across the dataset, **F)** distance-dependence (slope) of QC-FC correlations, **G)** mean RSFC correlation across the dataset, and **H)** mean distance-dependence (slope) of RSFC correlations. Also shown are **I)** change in between-subjects variance of RSFC correlations due to censoring and **J)** the number of subjects removed due to volume censoring. Vertical lines show “optima” for each DQM, when identifiable. Results with and without global signal regression (GSR) are denoted +GSR and -GSR; where these identifiers appear in a Y-axis label (panels E and G), it indicates that -GSR and +GSR results are presented at different scales. Data in panel **J** are identical for +GSR and -GSR.

The rank-order correlation between QC-FC correlations and ROI pair distance (**Figure 1C;** see Ciric et al., 2017; Muschelli et al., 2014; Parkes et al., 2018; Power et al., 2012; Satterthwaite et al., 2013) raises a distinct set of concerns. While standard FD censoring (with or without GSR) results in an apparent rapid “improvement” to an absence of distance-dependence (i.e., the complete removal of the typically observed effect that short-distance RSFC correlations have a stronger relationship to QC measures such as mFD than do long-distance RSFC correlations), further censoring with standard FD reverses the distance-dependence effect. This challenges the notion that standard FD censoring is simply removing the distance-dependent effect of motion artifact, because in that case a reverse-distance-dependence effect should not be possible – instead, distance-dependence should reach or approach 0 and then plateau. It is unclear how this reversal should be interpreted. While LPF-FD requires substantially more censoring to reach a QC-FC distance correlation of 0 (**Figure 1C**), the fact that it does *not* result in QC-FC distance correlations greater than 0 may indicate that it is a preferable censoring method, although censoring past approximately 50% of frames removed causes this metric to worsen again. Regardless, the behavior of this metric across these censoring approaches undermines clear interpretation of results employing this DQM.

We also evaluated four DQMs from a generalized linear model (GLM) approach developed by Burgess and colleagues (Burgess et al., 2016) that estimates a) the mean QC-FC value over all ROI pair correlations **(Figure 1E;** see Ciric et al., 2017; Muschelli et al., 2014; Power et al., 2014; Power et al., 2015; Satterthwaite et al., 2013), b) the slope of QC-FC and distance **(Figure 1F;** see Ciric et al., 2017; Power et al., 2012; Satterthwaite et al., 2012), c) mean RSFC **(Figure 1G),** and d) the mean distance-dependence of RSFC correlations (**Figure 1H;** see Power et al., 2012; Power et al., 2014; Power et al., 2015; Satterthwaite et al., 2012). Results from the slope of QC-FC and distance (**Figure 1F**) almost precisely mirror the correlation between QC-FC and distance (**Figure 1C**), as discussed above. With GSR, mean QC-FC (**Figure 1E**) begins near zero (the theoretically optimal value) and only exhibits small fluctuations with additional censoring. Without GSR, mean QC-FC shows steady reductions with more censoring, but these values become negative, raising a set of questions parallel to those outlined above regarding the QC-FC distance correlation shown in **Figure 1C**; namely, it is unclear how to interpret distance-dependence measures when increasingly aggressive censoring pushes the metric past its theoretically optimal value into a “reversed” or “flipped” state from the uncensored data.

However, the two measures based directly on RSFC correlations (**Figure 1G-H**), rather than on their association with mFD (as in the QC-FC DQMs), suggest that regardless of censoring method increasingly stringent censoring results in continuous reductions in both distance-dependence and the magnitude of RSFC correlations. Unlike the other DQMs, this provides a clearly interpretable result, although it leads to the unfortunate conclusion that even the smallest motions have a measurable impact on RSFC, and that this influence can only be removed by censoring until the entire dataset has been discarded—clearly, not a practicable solution to the issue of motion in RSFC.

Finally, **Figure 1I-J** shows the average (across ROIs) change in between-subject variance in the dataset as a result of FD-based censoring (and for DV-based censoring in **Figure S6**), as well as the number of participants removed from the dataset at each threshold. This demonstrates that the behavior of the DQMs shown in **Figure 1A-F** cannot be attributed to removal of subjects causing changes to the dataset. In addition, increasingly aggressive censoring leads to marked reductions in between-subjects variance, demonstrating that the removal of the highest FD volumes from the dataset makes RSFC values across subjects more similar. Moreover, this effect continues well past the putatively ‘optimal’ censoring thresholds identified by every DQM evaluated here. Thus, there are clear data-quality impacts of volume censoring on RSFC correlations (at least in terms of reducing motion-associated subject-level variability in RSFC) that occur at much higher censoring thresholds than are suggested by commonly used DQMs, or than is widely appreciated in the existing literature.

#### 3.1.2. Exploration of confounds in mFD-based dataset-QC metrics

One possibility raised by the above results is that there is a fundamental problem with DQMs based on QC-FC relationships. Every DQM reviewed above that depends on mFD behaves in a way that is not clearly interpretable, while every DQM that does not depend on subject-level QC metrics shows monotonic changes in the expected direction with additional censoring, and never exceeds a theoretically optimal value. Consequently, we conducted follow-up analyses to examine whether one or more confounds may impact DQMs that rely on SQMs.

We hypothesized that these DQMs could be confounded by true differences in RSFC correlations between high- and low-motion participants. That is, these DQMs rest on the assumption that the true relationship between mFD and RSFC will approach 0 as the motion artifact remaining in a dataset approaches 0. However, there could be true differences in RSFC between high- and low-motion individuals that are unrelated to motion artifact; that is, a “third variable” confound. This is consistent with reports that RSFC differences have been observed between high- and low-motion participants, even when only considering low-motion scans (Zeng et al., 2014), and that the quantity and quality of participant motion is significantly associated with participant demographic characteristics (Bolton et al., 2020; Siegel et al., 2017). Such confounds could call into question not just the assumption that the observed value of QC-FC correlations (or number of significant differences between high- and low-motion subjects) in a dataset should optimally be zero, but also that *any* target value for such metrics can be known a priori.

In light of these concerns, we decided to test whether two likely candidates for a third variable that could impact both RSFC correlations and participant motion may be impacting the QC-FC null rejection rate (the DQM reported in **Figure 1B**). **Figure 2** shows the observed proportion of RSFC correlations that significantly differ between high- and low-motion participants (upper and lower tercile on median FD, respectively) at various uncorrected P value thresholds for both the full HCP 500 dataset (*HCP Dataset 1*) and for a reduced dataset (*HCP Dataset 1b*) that excludes participants who had a parent with any psychiatric or neurological disorder (FH+ subjects), or who used any illicit substance or had a blood-alcohol level above 0.05 during the course of the study (SU+ subjects). As expected, FH+ and SU+ subjects had elevated median FD values (mean = 16.2) relative to FH- and SU- participants (mean = 14.7; *t_490_* = 3.06; *P* = 0.002). **Figure 2** also shows 95% confidence intervals from a Monte Carlo simulation of the effect of removing an equal number of randomly selected participants who exhibited an equivalent amount of motion to FH+ and SU+ subjects, but who were themselves FH- and SU- (see *Supplementary Section S1.3.*).

**Figure 2.**
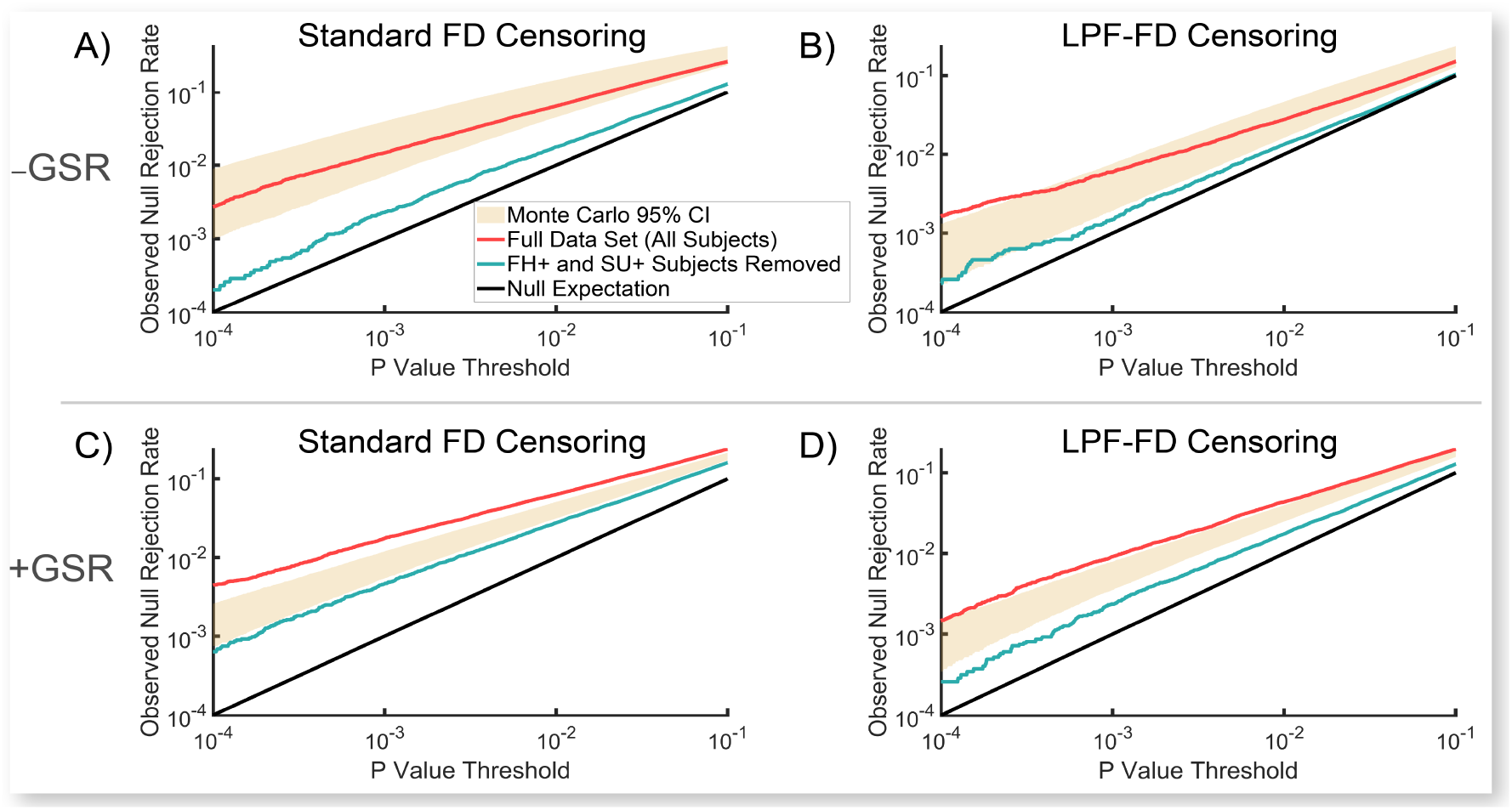
Proportion of significant RSFC correlations between high- and low-motion participants at various uncorrected P value thresholds in the HCP500 dataset (n = 501; *HCP Dataset 1)*. The proportion observed for the full dataset (red) and after removing subjects who tested positive for drug use or elevated blood alcohol content on the day of a scan session (SU+) or have a family history (FH+) of a psychiatric or neurological disorder (blue-green; n = 314; *HCP Dataset 1b*) are shown. 95% confidence intervals from 1,000 Monte Carlo resamplings of the data by instead removing FH- and SU- subjects with motion characteristics matched to the FH+ and SU+ subjects in the original dataset (matched on FD; see *Supplementary Section S1.3*) are shown in the beige area. Black lines represent the null expectation. Results are shown for analyses with standard volume censoring (left) and LPF-FD volume censoring (right), without global signal regression (GSR; –GSR, top) and with GSR (+GSR, bottom). Censoring thresholds selected to ensure an equal number of volumes removed across methods (thresholds for all methods were selected to match the percentage of volumes removed identified as optimal for LP-FD censoring using the methods described in *Section 2.3.2.5* and *Section 3.2.2.1*) in the HCP500 dataset (*HCP Dataset 1;* without GSR: 46.68%; with GSR: 43.32%): **A)** FD > 0.1475 mm **B)** LPF-FD > 0.0318 mm **C)** FD > 0.155 mm **D)** LPF-FD > 0.0337 mm.

We observed that removing FH+ and SU+ participants causes a significantly greater reduction in observed null hypothesis rejection rates than would be expected simply by removing an equivalent number of FH- and SU- participants who exhibit similar levels of motion. Thus, these findings are consistent with the “third-variable” effect hypothesized above, such that FH+ and SU+ participants exhibit *both* true differences in RSFC relative to FH- and SU- individuals, *and* elevated in-scanner motion, thereby producing a *true association* between RSFC and motion that is independent of motion-induced signal artifacts. These results are also consistent with previous findings that trait effects were still detectable in RSFC correlations in the HCP dataset, even after aggressive denoising methods were employed (Siegel et al., 2017).

One potential critique of the analysis reported in section 3.1.2.1 is that matching FH+ and SU+ subjects to FH- and SU- subjects on overall motion (i.e., mFD) may fail to capture important differences in the *type* of motion occurring in these two groups; that is, even if mFD is equivalent in two individuals, one individual may, e.g., exhibit relatively steady motion throughout the scan, while another exhibits lower motion through most of the scan but has several large individual motions that result in equivalent mFD values between these individuals. To evaluate this possibility, we developed a matching algorithm that finds pairs of subjects who have maximal overlap in the empirical cumulative density functions of the derivatives of their motion parameter (MP) traces. Results of this analysis are consistent with the results reported in **Figure 2**, and are reported in detail in *Supplementary Section S2.1*.

Thus, DQMs that depend on relationships between measured BOLD signal and an SQM (e.g., QC-FC measures) are clearly impacted by unmeasured third variables. Moreover, although we demonstrate here that FH status acts on both measures of participant motion and on RSFC values, thus confounding any DQM that depends on the relationship between motion and RSFC, we cannot rule out that other, unmeasured, individual differences could also have this impact. Thus, unless *all* participant characteristics that impact both motion and true RSFC correlations can be identified and modeled, DQMs that implicitly assume that such confounds do not exist should not be employed as objective criteria for evaluating the effectiveness of motion denoising strategies on RSFC data.

### 3.2. Development and evaluation of volume censoring with novel dataset-quality metrics (DQMs)

#### 3.2.1. Mean absolute change in resting state functional connectivity (MAC-RSFC)

##### 3.2.1.1. Visual evaluation of censoring methods

Because of the issues described above with established DQMs, we sought alternative methods of evaluating the impact of denoising on RSFC, beginning with a visual comparison approach employed in prior work (Power et al., 2012; Power et al., 2014). Changes in RSFC correlations resulting from motion-targeted volume censoring are shown for all ROI pairs (Power et al., 2011) in **Figure 3**; these changes are compared to RSFC changes resulting from removal of an identical number of randomly selected volumes within each run. Thresholds were selected such that an equivalent number of volumes were removed by each evaluated method, to allow an apples-to-apples comparison between censoring methods (the values chosen for display were the optimal number of volumes removed for LPF-FD for all FD measures and the optimal values for GEV-DV for all DV measures, determined separately for analyses with and without GSR, as presented in *Section 3.2.2.1*, below).

**Figure 3.**
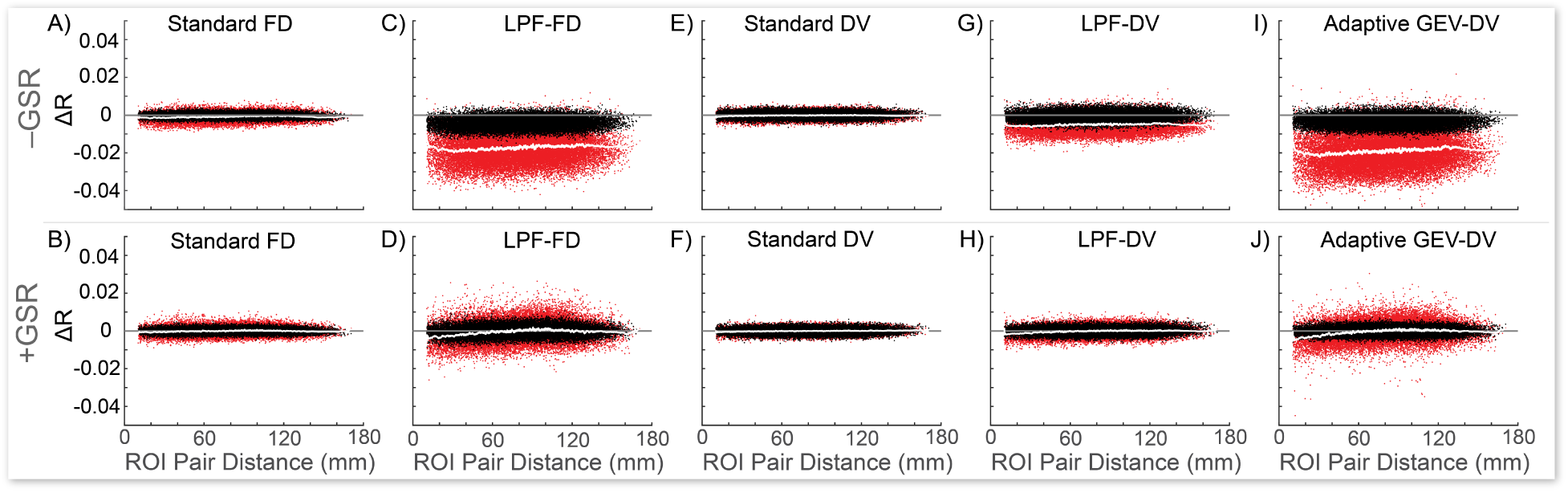
Changes in pairwise ROI RSFC correlations resulting from volume censoring. Results shown after censoring using standard FD **(A,B)** and standard DV **(E,F)**, LPF-FD **(C,D)** and LPF-DV **(G,H)**, and adaptive GEV-DV **(I,J)**, plotted against the Euclidean distance between ROIs on the X-axis, in the HCP500 dataset (n = 501; *HCP Dataset 1*). Red dots indicate the average change across the sample in RSFC correlation resulting from volume censoring for each ROI pair. Black dots indicate the average change resulting from random censoring of an equal number of volumes within each run. White lines indicate a sliding window of the mean of the red dots. Results are shown both without global signal regression (GSR; –GSR, top) and with GSR (+GSR, bottom). Censoring thresholds were selected such that an equal amount of data was removed by each method (without GSR: 46.68% for FD measures and 45.95% for DV measures; with GSR: 43.32% for FD measures and 37.95% for DV measures) as follows. Without GSR: **A)** FD > 0.1475, **C)** LPF-FD > 0.0318, **E)** DV > 61, **G)** LPF-DV > 10.75, and **I)** GEV-DV d_G_ = 1.16. With GSR: **B)** FD > 0.1550, **D)** LPF-FD > 0.0337, **F)** DV > 63, **H)** LPF-DV > 11.25, and **J)** GEV-DV d_G_ = 1.36.

Standard (unfiltered) censoring methods resulted in minimal changes in RSFC compared to random censoring of an equivalent number of frames within each run, consistent with the view that this results in the targeting of respiratory motion and pseudomotion in high-TR datasets (Power et al., 2019a), signal effects that, because of their relatively high frequency, should be effectively removed from the data by standard band-pass filtering of RSFC data, and thus are not ideal targets for volume censoring. Standard methods were outperformed by LPF-based censoring (**Figure 3A-H**), and GEV-DV censoring produced an even greater change in observed RSFC than LPF-DV censoring (**Figure 3G-J**). Finally, in analyses without GSR, all LPF and GEV methods produced an overall downward shift in RSFC magnitude that was not observed in analyses employing GSR (which is expected as GSR causes calculated RSFC correlations to be approximately centered on zero; Aquino et al., 2020; Fox et al., 2009; Glasser et al., 2018; Murphy et al., 2009; Murphy and Fox, 2017; Saad et al., 2012). Given that GSR has been shown to globally reduce the magnitude of RSFC correlations (Ciric et al., 2017; Murphy et al., 2009), this suggests that LPF-based methods may be producing some of the same effect as GSR in minimizing the impact of motion on overall RSFC correlation magnitude.

It is also worth noting that results for standard FD and DV censoring (**Figure 3A, B, E, F**) do not show the distance-dependence typically observed in prior work (e.g., Power et al., 2012; Power et al., 2014), further demonstrating that standard FD and DV censoring do not behave as expected in the HCP dataset, likely because of the effect of respiration-related motion and pseudomotion on motion parameters and RSFC timeseries in multiband data, as discussed in *Section 1.1* and shown in **Figure S1**. However, this cannot explain our failure to reproduce the results of Burgess and colleagues in the HCP dataset (Burgess et al., 2016). Although a thorough empirical investigation of this issue is beyond the scope of the work we present here, we suspect that this discrepancy occurred because Burgess and colleagues did not use a standard band-pass filter of 0.009 – 0.08 Hz (or 0.01 – 0.1 Hz) in preprocessing their data, instead using only a 0.009 Hz high-pass filter. Thus, higher frequency effects that were removed from our data will have remained in the data as analyzed by Burgess and associates, and may have produced the distance-dependent artifacts that standard censoring methods were then able to remove. Moreover, we would note that we did not fail to observe distance-dependent effects more broadly (see, e.g., *Section 3.2.2.1*, below), only that standard FD and DV censoring did not produce these effects.

##### 3.2.1.2. Quantitative evaluation of censoring methods using MAC-RSFC

Next, we attempted to quantify the visual comparison in **Figure 3** to evaluate it across a range of censoring parameters, as we did for other DQMs. Thus, we developed a new DQM based on this visual comparison: we compute the mean absolute value of the within-subject change in RSFC correlations relative to randomly removing an equivalent number of volumes within each run, across all ROI pairs, and across all subjects in the sample, thus effectively quantifying the spread of red dots away from the black dots in **Figure 3** (see *Section 2.3.1* for methodological details). Observed between-method differences in this metric should be specifically associated with differences in targeting of BOLD signal fluctuations associated with the FD and DV values used in the method (i.e., unfiltered, band-pass filtered, or low-pass filtered; and for GEV-DV versus other methods, the difference between a single study-wide DV threshold versus run-specific DV thresholds). We term this measure MAC-RSFC (Mean Absolute Change in Resting State Functional Connectivity). **Figure 4** (which summarizes over 10^16^ partial correlations) shows that LPF-FD and LPF-DV produced larger magnitude changes in RSFC correlations than both standard FD- and DV-based censoring, relative to random removal of volumes, regardless of how much data is removed by each method. That is, across the full range of possible censoring thresholds (from 0% to nearly 100% of data removed), the LPF-based methods we propose here significantly (given the non-overlap of 95% confidence intervals obtained from bootstrapping) outperformed all other censoring methods evaluated here (but see *Section 3.2.3* for a discussion of the critical assumptions underlying this interpretation of these results). In addition, GEV-DV outperformed LPF-DV, suggesting that, in line with our observations in **Figure S2** and **Figure 3G-J**, an adaptive thresholding method is preferable for handling the substantial differences in central tendency of DV measures across runs. Finally, it should be noted that MAC-RSFC is intended only for comparison across volume censoring methods rather than for selecting optimal parameters or thresholds for any given censoring method, as the monotonic increase of MAC-RSFC as a function of increasingly strict censoring parameters observed in **Figure 4** makes it unsuitable for selecting optima within methods (for this purpose, we developed ΔMSE-RSFC, discussed in Section 3.2.2, below).

**Figure 4.**
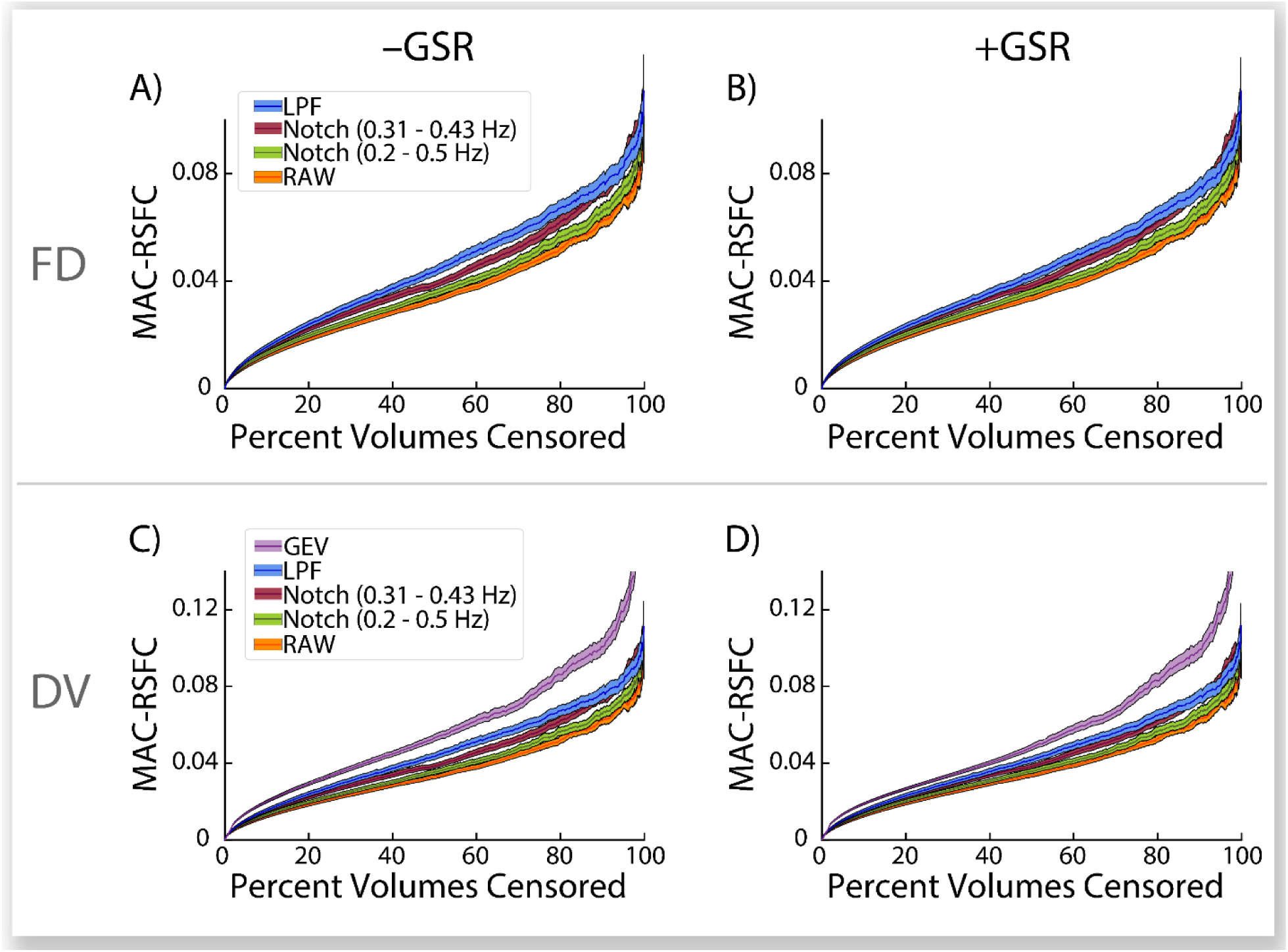
Mean Absolute Change in Resting State Functional Connectivity (MAC-RSFC) and 95% confidence intervals (CIs) as a function of the percent of volumes censored in the HCP500 dataset (n = 501; *HCP Dataset 1*). Results shown after volume censoring using standard FD, Notch-FD, and LPF-FD **(A,B)**, and standard DV, Notch-DV, LPF-DV, and GEV-DV **(C,D)** across a range of parameter values. 95% CIs on MAC-RSFC were estimated via bias corrected and accelerated bootstrapping with 10,000 bootstrap samples. Results are shown both without global signal regression (GSR; –GSR, left) and with GSR (+GSR; right). Notch (band-stop) filter-based censoring methods were evaluated using stopbands of 0.31 Hz – 0.43 Hz and 0.2 Hz – 0.5 Hz.

As noted above, other authors have recently suggested employing band-stop (notch) filters to separate respiration-related motion from other motion in fast-TR data such as the HCP dataset (Fair et al., 2020; Power et al., 2019a). Although they did not directly compare the efficacy of a band-stop filter to other methods (nor did they propose applying such filters to voxelwise data prior to calculation of DV), we additionally show in **Figure 4** that while these methods both outperform standard, unfiltered, approaches to FD-based censoring on MAC-RSFC, they perform more poorly than LPF methods. This result is expected when RSFC preprocessing includes band-pass filtering with a cutoff frequency below that which is used for low-pass or band-stop filtering of MPs (here, 0.009 Hz – 0.08 Hz band-pass filtering), as do the wide majority of published analyses. High-frequency components of MP traces and voxel timeseries that are removed during the calculation of LPF-FD and LPF-DV, respectively, will largely correspond to high-frequency components of motion artifact that are removed from BOLD timeseries when they are band-pass filtered. Thus, censoring methods that use low-pass filtered framewise statistics for targeting volumes are expected to produce superior estimates of which volumes truly contain unremoved artifact relative to methods that use band-stop filtering, at least when the underling timeseries data has been band-pass filtered, as it has been here and in the majority of published RSFC studies.

#### 3.2.2. Optimization of volume censoring thresholds: A bias-variance decomposition approach

It has been noted that, irrespective of the censoring method used, increasingly strict thresholds result in continuous “improvements” in data quality (Caballero-Gaudes and Reynolds, 2017; Power et al., 2014; Power et al., 2015), an observation that is also borne out by all DQMs evaluated here that do *not* depend on SQMs such as mFD (**Figure 4** and **Figure 1G-H**). This raises a critical challenge as to how to balance the tradeoff between the benefits of additional denoising and the costs of discarding additional data. Presently, such thresholds are typically selected from previously established thresholds (e.g., FD > 0.2 mm or FD > 0.5 mm in singleband data), or are sometimes adapted for particular datasets through qualitative visual inspections of a variety of DQMs. However, such approaches do not rest on a rigorous quantitative optimization, and the only existing quantitative approach to selecting FD thresholds has significant flaws (see *Section 2.3.2.1*). Thus, we developed a novel quantitative approach to finding optimal censoring thresholds, termed ΔMSE-RSFC, which is based on an estimated bias-variance decomposition of the mean squared error (MSE) of sample mean RSFC correlations (see *Section 2.3.2.1* through *Section 2.3.2.6*).

##### 3.2.2.1. Optimization of volume censoring parameters in the HCP500 dataset using ΔMSE-RSFC

**Figure 5** shows this approach for LPF-FD and GEV-DV censoring (used separately). **Figure 5A** demonstrates that increasingly aggressive censoring (nearly) continuously reduces motion-induced bias in RSFC, with significantly greater effects in analyses without GSR. Additionally, **Figure 5B** shows that both censoring methods produce a reduction in between-subjects variance that exceeds the increase caused by increasing sampling error until approximately 40-50% of volumes are removed. As variance is reduced before any subjects are removed (**Figure 5C**), the reduction in variance is not due to removal of high-motion participants; rather, it is due to the exclusion of volumes that were impacting individual RSFC correlation estimates in higher-motion subjects. Thus, contrary to our initial expectations, power to resolve sample-level RSFC correlations *increases* with more aggressive censoring, despite the loss of temporal degrees of freedom resulting from reduced between-subject variance that exists because of motion effects on RSFC correlations. Finally, ΔMSE-RSFC, calculated as the change in the ratio of the sum of squared bias and variance to the number of remaining subjects (see *Section 2.3.2.2*), produces a U-shaped curve when plotted against percent frames removed (**Figure 5D**), and is thus suitable for optimization as described in *Section 2.3.2.5*. The global minimum (the greatest magnitude reduction in MSE-RSFC) represents the point at which maximal improvement in data quality is achieved in this dataset for each method: beyond this point, while further removal of data will result in a reduction in bias, that reduction will be accompanied by a *larger* increase in variance that results in a poorer overall estimate of the true value of RSFC correlations in the sample. The optimal single-method censoring parameters for LPF-FD were *Φ_F_* = 0.0318 mm and *Φ_F_* = 0.0337, and for GEV-DV were *d_G_* = 1.16 and *d_G_* = 1.36, respectively for analyses without and with GSR (see **Table 2**).

**Figure 5.**
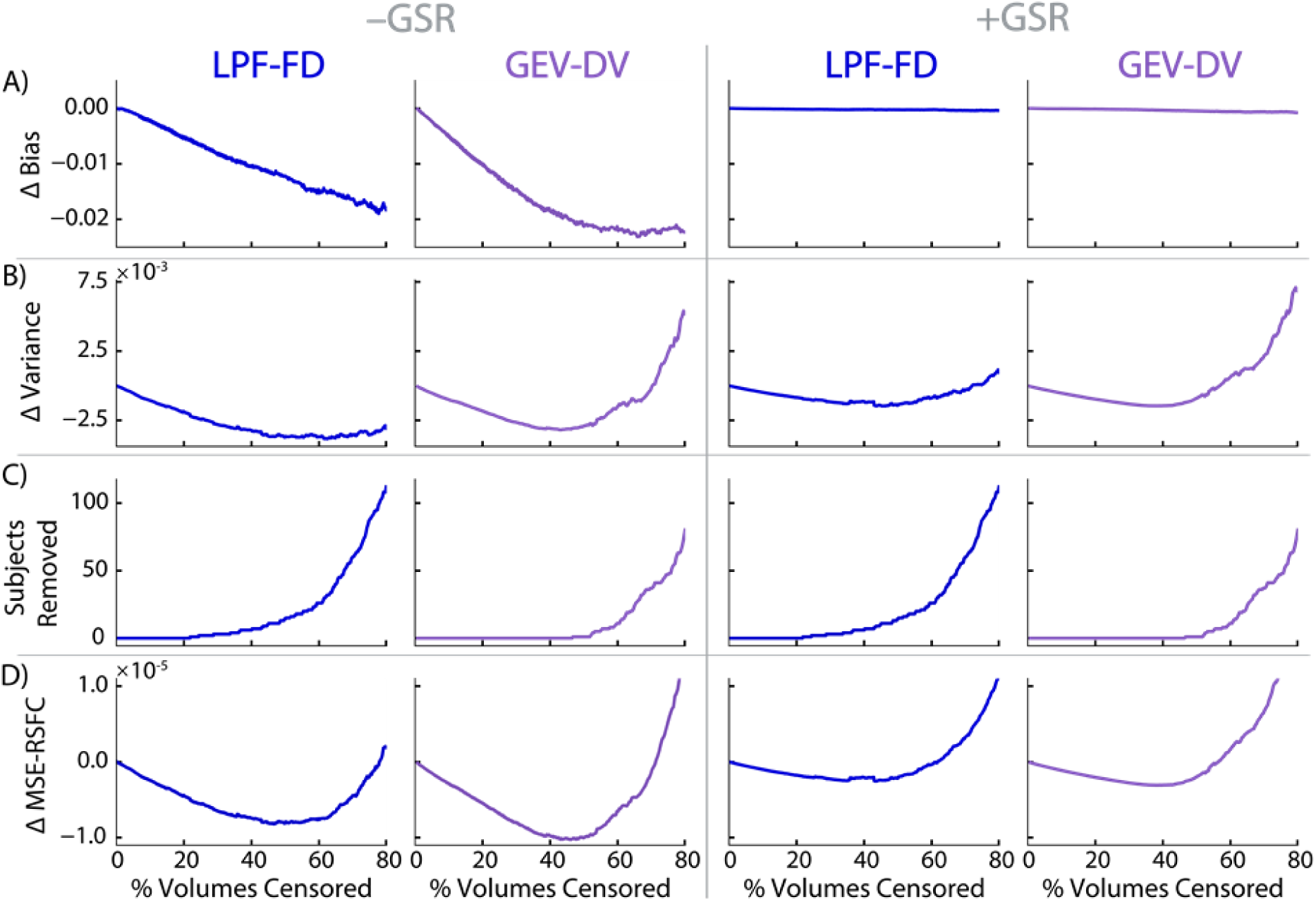
Change in sample statistics used to calculate ΔMSE-RSFC due to volume censoring in the HCP500 dataset (n = 501; *HCP Dataset 1*). A) Mean change in sample average Z-transformed ROI pair correlations, over and above that due to random removal of an equivalent number of randomly selected volumes within each run. **B)** Mean change in between-subjects variance in sample ROI pair correlation estimates. **C)** Number of subjects removed from the sample due censoring. **D)** Change in estimated mean square error (MSE) divided by the remaining number of subjects, ΔMSE-RSFC, averaged across ROI pairs. Analyses without global signal regression (GSR) are denoted by –GSR (left); those with GSR are denoted by +GSR (right).

**Table 2.** Optimal parameters for volume censoring determined by minimizing *ΔMSE-RSFC in the HCP500 dataset*.

| **Optimal Volume Censoring Parameters for the HCP500 Dataset (HCP Dataset 1)** |  |  |
| --- | --- | --- |
|  | **No GSR** | **With GSR** |
| **LPF-FD Volume Censoring Only** |  |  |
| LPF-FD $\Phi_F$ (mm) | 0.0318 | 0.0337 |
| % Frames Removed | 46.68 | 43.32 |
| **GEV-DV Volume Censoring Only** |  |  |
| GEV-DV $d_G$ (dimensionless) | 1.16 | 1.36 |
| % Frames Removed | 45.95 | 37.95 |
| **Combined LPF-FD and GEV-DV Volume Censoring** |  |  |
| LPF-FD $\Phi_F$ (mm) | 0.0339 | 14.336 |
| GEV-DV $d_G$ (dimensionless) | 1.39 | 1.30 |
| $\Phi_F / d_G$ ratio (mm) | 0.02438 | 11.0277 |
| Frames Removed (%) | 56.15 | 40.06 |

*GSR = global signal regression. LPF-FD = low-pass filtered framewise displacement. GEV-DV = generalized extreme value derivative of timecourses, root-mean-squared variance over voxels. All results are from analyses using the HCP 500 dataset (n = 501; HCP Dataset 1).*

To allow for direct comparison between optimal volume censoring thresholds derived from ΔMSE-RSFC and those suggested by traditional DQMs (evaluated in **Figure 1**), we show in **Figure 6** the difference in RSFC correlations produced by optimal LPF-FD censoring using ΔMSE-RSFC over and above the optima suggested by minimizing median absolute QC-FC (Ciric et al., 2017), in *HCP Dataset 1a* (i.e., after removing 26 high-motion subjects from *HCP Dataset 1* in order to control for the effect of subject removal by excessive volume censoring; see *Section 2.1.1.1*). This shows an apparent general global reduction in RSFC correlations (without GSR), as well as removal of distance-dependence when compared to the censoring threshold suggested by mean absolute QC-FC, consistent with a further reduction in motion artifact remaining in the dataset following censoring with the optima suggested by ΔMSE-RSFC. Equivalent plots for all other DQMs also show similar results when using LPF-FD (**Figure S7**) and GEV-DV (**Figure S8)** volume censoring. Plots of ΔMSE-RSFC showing minima for *HCP Dataset 1a* for LPF-FD and GEV-DV censoring are shown in **Figure S9**. In summary, we show that censoring at the more aggressive thresholds provided by ΔMSE-RSFC optimization results in a reduction in the distance-dependence of RSFC correlations when compared to *any* known DQM that relies on SQMs such as mFD (i.e., QC-FC measures).

**Figure 6.**
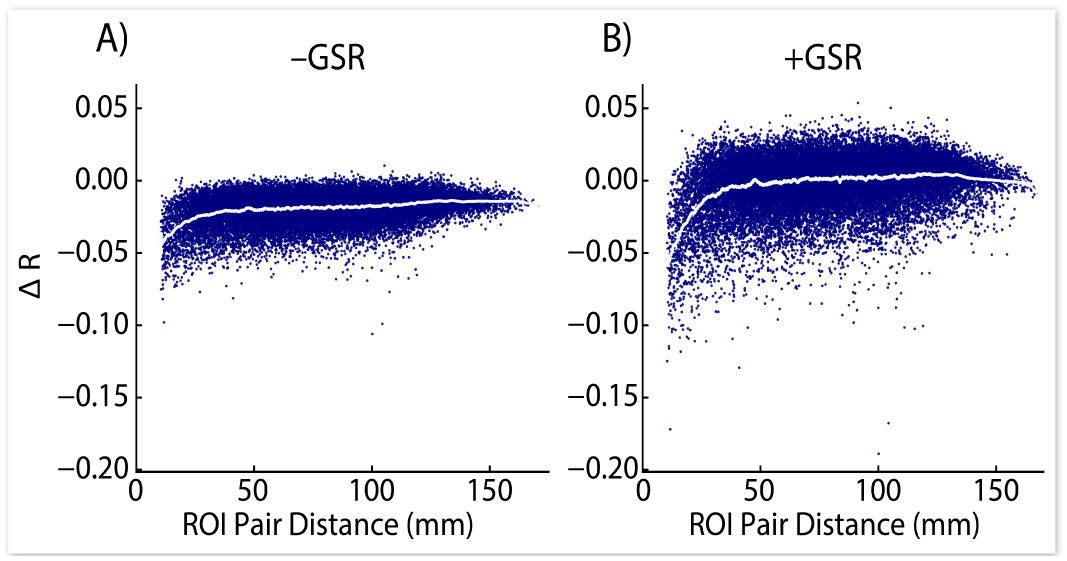
Change in ROI pair correlations resulting from additional censoring as indicated by ΔMSE-RSFC, over and above the optimal threshold indicated by Median Absolute QC-FC, in a subset of the HCP 500 dataset (n = 475; *HCP Dataset 1a*). Results shown for analyses without global signal regression (GSR; –GSR, left) and with GSR (+GSR, right).

Next, we determined optimal combined thresholds for LPF-FD and GEV-DV censoring when used together by seeking the global minimum of *ΔMSE-RSFC* in the space produced by the free parameter for each of the two methods. These results suggest that a relatively restrictive threshold for LPF-FD is required for analysis without GSR, but that it is optimal to rely primarily on GEV-DV censoring for data employing GSR. That is, it is likely that the artifactual signals that are indexed by increasing LPF-FD values, but that are not *also* accompanied by outlying values of LPF-DV at the same timepoint, are largely removed using GSR. Consequently, when used in concert with GSR, LPF-FD values can be largely disregarded so long as relatively aggressive GEV-DV censoring is carried out. Optimal censoring parameters for combined LPF-FD + GEV-DV censoring are: without GSR, *Φ_F_* = 0.0339 mm and *d_G_* = 1.39; with GSR, *Φ_F_* = 14.336 and *d_G_* = 1.30 (**see Table 2**).

##### 3.2.2.2. Optimization of volume censoring parameters in arbitrary datasets

Finally, we tested the ΔMSE-RSFC optimization procedure in a validation set collected at the New York State Psychiatric Institute (NYSPI) as described in *Section 2.3.2.6*, and present the results in **Figure 7**. As in the HCP500 dataset, motion-induced bias in RSFC correlations decreases monotonically (**Figure 7A-B**) with increasing strictness of censoring parameters. Variance initially decreases, reaching minima before 20% of volumes are censored across the dataset, before increasing substantially (**Figure 7C-D**) and becoming unstable as a result of excessive removal of data, including data from entire subjects (**Figure 7E-F**). Calculated ΔMSE-RSFC is shown in **Figure 7G-H.**

**Figure 7.**
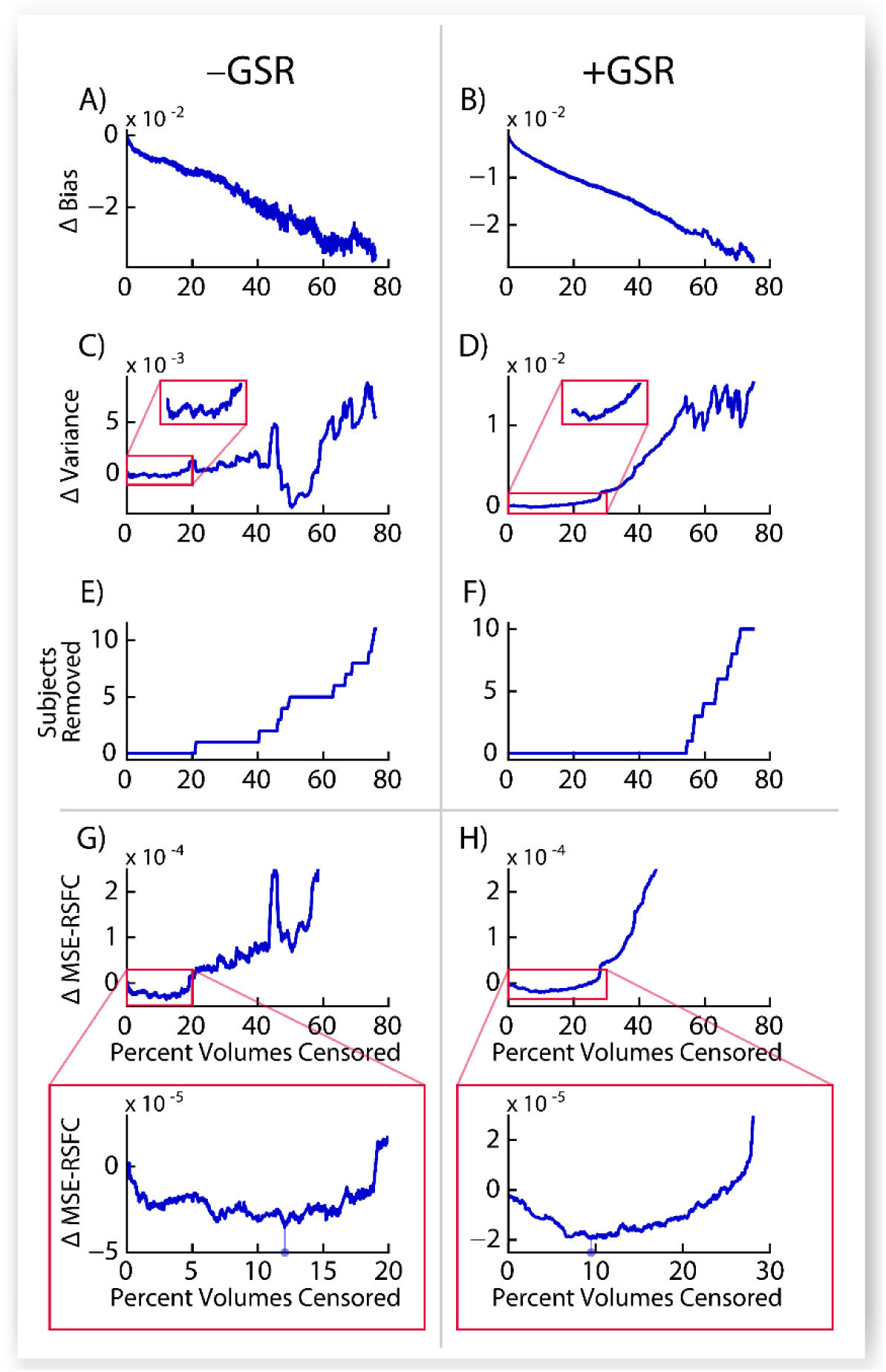
Optimization of volume censoring parameters by minimizing ΔMSE-RSFC in the New York State Psychiatric Institute (NYSPI) dataset (n = 26). Shown are, as a function of percent of frames removed due to volume censoring across the dataset, the change in bias **(A,B)**, change in variance **(C,D)**, number of subjects removed by censoring **(E,F)**, and the resulting ΔMSE-RSFC **(G,H)**. The insets in panels G and H display the region within which the minimum ΔMSE-RSFC was found; note the different axes in the insets of panels G and H. Drop-down lines in panels G and H show the detected ΔMSE-RSFC minima.

The *NYSPI Dataset* shown here is considerably smaller than the HCP500 dataset, with approximately 5% the number of subjects. In spite of this, ΔMSE-RSFC appears to be a sufficiently stable DQM with clearly identifiable minima that are suitable targets for optimization in analyses both with and without GSR (**Figure 7G-H**; see inset). Identified optimal censoring parameters in the NYSPI Dataset (**Table 3**) are considerably more lenient than those in the HCP500 dataset (see **Table 2**), likely resulting from the greater “penalty” associated with data loss in a smaller dataset. Critically, the inclusion of *N* in the denominator for calculation of ΔMSE-RSFC accounts for the loss of statistical power associated with a reduction in number of subjects remaining in the dataset after censoring, even if the removal of these subjects reduces total variance across the sample. This property makes ΔMSE-RSFC robust even in datasets in which minimum variance is achieved in a region of instability (due to extensive removal of subjects unsupportable by the dataset’s sample size, e.g., **Figure 7C**), instead selecting censoring parameters that retain substantially more data (e.g., **Figure 7G**).

**Table 3.** Optimal parameters for volume censoring *in the NYSPI Dataset*.

|  | No GSR | With GSR |
| --- | --- | --- |
| LPF-FD $\Phi_F$ (mm) | 0.1158 | 47.5294 |
| GEV-DV $d_G$ (dimensionless) | 4.74 | 4.32 |
| Frames Removed (%) | 12.063 | 9.495 |
| Subjects Removed | 0 | 0 |

*GSR = global signal regression. LPF-FD = low-pass filtered framewise displacement. GEV-DV = generalized extreme value derivative of timecourses, root-mean-squared variance over voxels. All results are from analyses using the NYSPI dataset ( $n = 26$ ).*

#### 3.2.3. Caveats and limitations of MAC-RSFC and ΔMSE-RSFC

The methods described throughout this work—and more generally any methods that ultimately rely upon summarizing functional connectivity between voxels or ROIs using a temporally invariant set of values (e.g., Pearson’s correlations)—necessarily reduce all dynamic components of brain activity into temporally invariant descriptors, which represent an average of all constituent brain states captured over the course of a BOLD acquisition. Concerns have been raised by Glasser and colleagues (2018) that non-selective fMRI denoising techniques (e.g., volume censoring and GSR) may be removing true, non-artifactual, variation in functional connectivity that occurs over the course of fMRI timeseries, and that DQMs used to optimize denoising methods cannot distinguish the removal of artifact from the removal of true neural signal. Certainly, in the case of volume censoring, brain states that co-occur with motion will disproportionately be captured by volumes that are flagged for removal by framewise statistics (e.g., those based on forms of FD and DV), and thus will be disproportionately removed from timeseries prior to the calculation of connectivity statistics. In the (likely) event that motion is associated with entering or exiting sleep-states (which are common during RSFC; see Tagliazucchi and Laufs, 2014) this may not be especially problematic, as many investigators may prefer to minimize the quantity of sleep impacting their data. We cannot rule out, however, that other brain-states exist that are associated with both motion and true differences in RSFC, or, more broadly, that DQMs evaluating the effectiveness of volume censoring are favoring the removal of true neural signal rather than removal of artifactual signals. We focus most heavily on the former issue here, because it is most germane to volume censoring and related approaches that selectively remove specific *timepoints* from a dataset. However, it should be noted that the latter concern regarding removal of true neural signal could apply to any DQM employed in evaluating any of a wide variety of denoising techniques, ranging from volume censoring (as in this work) to other methods that are applied to entire timeseries, including (but not limited to) GSR and ICA-based approaches, unless a “positive control” of some kind is employed in order to demonstrate that desirable neural signal is not being removed.

Fundamentally, the logic of volume censoring as a method of artifact mitigation rests on the assumption that the metric used to identify volumes impacted by an artifact is not systematically associated with brain-states of interest. If it becomes evident that motion occurs more frequently during particular brain-states that are relevant to the hypotheses under consideration, then volume censoring should either not be employed in favor of other denoising techniques such as temporal independent components analysis (tICA; Glasser et al., 2018), or otherwise used with extreme care to avoid confounding. Ultimately, however, these issues are beyond the scope of the work presented here, which instead attempts to adjudicate among several proposed censoring methods and produce a broad set of recommendations for investigators who intend to use volume censoring with multiband RSFC data. We raise these issues here because the logic of MAC-RSFC and ΔMSE-RSFC follow directly from them.

If an investigator accepts the view that FD and DV (or any variant thereof, such as LPF-FD and GEV-DV) is not positively associated with any brain-state of interest, then because MAC-RSFC is calculated as a subtraction from random censoring, changes in MAC-RSFC as a result of censoring *must be* beneficial changes that result from removing volumes impacted by artifact. This follows because any impact on RSFC of removing volumes that is not a direct result of removing the highest FD or DV frames in a volume is accounted for by the comparison to random removal of volumes (averaged over 10 randomizations to reduce noise in the estimate). In contrast, however, if an investigator does *not* accept the assumption above, then MAC-RSFC could easily increase, and make a censoring method appear “better,” as a result of removing true neural signals from the data if such neural signals are simultaneously highly distinct from the temporal average and correlated with periods of greater than baseline subject motion (FD) or signal change (DV). Additionally, it should be further emphasized the MAC-RSFC is intended solely for quantitatively comparing volume censoring methods to one another (e.g., Standard FD censoring to LPF-FD censoring), rather than selecting optimal parameters for any given method (in contract with ΔMSE-RSFC). This is because, as noted at the outset of *Section 3.2.2*, if the maximal value of MAC-RSFC is taken to be optimal, there would be no data remaining in the RSFC dataset. This was our primary motivation for developing ΔMSE-RSFC, which instead tries to find an optimal balance between reduction of motion-induced bias and data loss due to excessive volume censoring.

Likewise, in calculating ΔMSE-RSFC, reduced variance and “bias” in RSFC correlations observed to result from volume censoring could be due to not just a reduction in motion artifact, but also the disproportionate removal of motion-associated brain states that may occur more frequently in some individuals than in others. That is, if a substantial fraction of these brain states are themselves correlated with motion events (i.e., if dynamic fluctuations in functional connectivity are correlated with FD- and/or DV-based measures), and if motion-associated neural activity varies considerably across individuals, then one could observe a reduction in variance resulting from increasingly aggressive volume censoring, as in **Figure 1I** and **Figure S6B,** due to the disproportionate removal of data during these states. Similarly, motion-associated neural events that produce a true net change in RSFC correlations across a sample would manifest as a reduction in bias when performing censoring. In this case, volume censoring would be removing true neural signal along with motion artifact. However, as stated previously, this is a general limitation of motion denoising techniques that rely on the removal of motion-contaminated volumes, such as volume censoring and spike regression, and others including GSR (and mean greyordinate timeseries regression), as well as any RSFC statistics that analyze timeseries as stationary processes.

Nonetheless, we undertook two “positive control” analyses that had the potential to demonstrate that increasingly aggressive censoring with LPF-FD and GEV-DV, as we recommend here based on results from MAC-RSFC and ΔMSE-RSFC, resulted in the removal of true neural signal. These analyses are described in *Supplementary Section S1.6* and *Supplementary Section S2.2*. Although these analyses cannot conclusively demonstrate that no neural signal is being removed by volume censoring, they nonetheless represent an opportunity for falsification of the claim that neural signal is not being removed. As discussed in *Supplementary Section S2.2*, neither analysis suggests that increasingly aggressive volume censoring results in substantive removal of true neural signal until a large proportion of data are removed, well beyond that which our methods consider to be optimal in any of the datasets analyzed here. In either case, while the selection of volume censoring as a component of motion denoising pipelines is beyond the scope of this manuscript, we encourage investigators to be fully cognizant of the assumptions inherent to denoising methods employed in their work, as well as the relationships between these assumptions and the hypotheses under investigation.

## 4. Conclusions

### 4.1. Overview

The work presented here provides several significant advances in preprocessing strategies for reducing the impact of participant motion with volume censoring in RSFC studies employing acquisitions with multiband acceleration and fast repetition times (TRs), as well as in evaluating the success or failure of volume censoring-based denoising pipelines. Specifically, we evaluated a number of dataset-quality metrics (DQMs) that have been widely used in the literature, and show that DQMs that rely on QC-FC measures, such as the correlation between RSFC correlations and subject-level QC measures (SQMs), systematically exhibit erratic behavior when assessed over a comprehensive range of censoring thresholds, raising serious questions as to their appropriateness in this context. Expanding on this observation, we further show that at least one “third variable,” family history of psychiatric or neurological disorder, critically confounds these DQMs as it leads to both true differences in RSFC and higher estimates of motion. In conclusion, these findings argue strongly that DQMs that depend on the relationship between RSFC and SQMs such as mean framewise displacement (mFD), including those that have been widely employed throughout the RSFC fMRI literature, should not be used as objective, quantitative benchmarks in the assessment and optimization of motion denoising pipelines. Notably, although we did not directly employ single-band RSFC data in this work, the concerns we raise regarding third variable confounds could potentially extend to single-band or otherwise lower-TR RSFC datasets.

Next, we developed two new DQMs, MAC-RSFC and ΔMSE-RSFC, to serve in place of standard DQMs. The first, MAC-RSFC, simply quantifies the average change in RSFC estimates resulting from motion-targeted volume censoring, over-and-above random (not motion-targeted) removal of an equivalent number of volumes within each run, and is intended for use in quantitatively comparing across volume censoring methods. We used MAC-RSFC to show that the LPF-FD and GEV-DV methods proposed here result in larger motion-targeted changes in RSFC correlations across all censoring thresholds than both unfiltered FD and DV and notch filtered FD methods proposed elsewhere (Fair et al., 2020; Power et al., 2019a). As discussed in *Section 3.2.1.2*, the improved performance relative to notch filtered FD is likely a result of the fact that timeseries data is band-pass filtered prior to the calculation of RSFC correlations, which means the high frequency signals retained by notch filtering methods necessarily cannot impact RSFC correlations, as they have already been filtered out in the underlying data. The second DQM, ΔMSE-RSFC, is designed explicitly for optimization of denoising methods (e.g., selecting volume censoring parameters or thresholds), in that it attempts to minimize the error in estimates of true RSFC correlation values in a sample. Finally, we use this latter DQM to estimate optimal censoring parameters for the HCP 500 dataset and provide publicly available code that will allow investigators to estimate optimal parameters for other fMRI datasets using a simplified version of the ΔMSE-RSFC optimization methods described herein. We expect these methods to considerably improve the reliability and reproducibility of resting-state fMRI studies by facilitating improved data quality through motion denoising.

### 4.2. Limitations and Further Considerations

The novel methods presented here were specifically designed for high-TR multiband data, and we would advise against generalizing these findings without further evaluation to singleband datasets, although low-pass filtering motion estimates has indeed been found to improve denoising in single-band data (Gratton et al., 2020). However, although these methods were developed specifically on the publicly available HCP500 data release, they were further validated on an independent dataset, and we do expect that they will generalize broadly to other multiband datasets of similar TR. Notably, the optimal parameters we obtained for the NYSPI dataset were substantially more lenient than those obtained for the HCP500 dataset, indicating that optimal censoring parameters should not be naively applied across datasets without conducting the dataset-specific optimization procedure described herein, which can be carried out using the publicly-available code accompanying this article (see *Section 5.3*).

While throughout this manuscript we present results from analyses with and without GSR side-by-side, this procedure for optimizing censoring parameters should additionally be repeated for denoising pipelines that utilize volume censoring together with other methods, including RETROICOR (Glover et al., 2000), ICA-FIX (Griffanti et al., 2014), ICA-AROMA (Pruim et al., 2015), temporal ICA (Glasser et al., 2018), and DiCER (Aquino et al., 2020), before these results can be extended to those contexts.

Denoising techniques that utilize physiological recordings of respiration, including RETROICOR (Glover et al., 2000), may be used in particular to partly mitigate the impact of respiratory motion and pseudomotion (Caballero-Gaudes and Reynolds, 2017; Kassinopoulos and Mitsis, 2019; Xifra-Porxas et al., 2021), though incompletely (Glasser et al., 2018; Power et al., 2020; Power et al., 2019a; Power et al., 2017). To maximize generalizability to studies in which physiological data are of variable quality or are unavailable, as is often the case (Power et al., 2019a), we did not consider these methods here. Future work should seek to determine optimal volume censoring parameters across a wider variety of denoising pipelines.

As a final note, we discuss in *Supplementary Section S2.3* and show in **Figure S10** that the distribution of motion artifact systematically varies across runs within each session, and over time within each run, with the highest-quality data acquired near the start of each run, and during the first run of each session. Notably, a substantial “rebound” towards higher quality data occurs at the beginning of the second run, and thus it is advisable to employ a larger number of shorter runs rather than a smaller number of long runs. Specifically, **Figure S10** strongly suggests that employing twice as many runs at half the length employed in the HCP study could have resulted in substantially less participant motion than was observed.

## 5. Acknowledgements

### 5.1 Assistance

Drs. Mark Slifstein, Hongshik Ahn, Joseph Schwartz, Yuefan Deng, and Anissa Abi-Dargham provided helpful commentary on various aspects of the methods presented here. Zu Jie Zheng, Alexander Eichert, Eilon Silver-Frankel, Hung-Wei (Bernie) Chen, and Sameera Abeykoon assisted with deploying analysis software to the SeaWulf cluster and generating figures. Mutahira Bhatti, Umaimah Nawaz, Justin Beutel, and Ayman A. Khan assisted with editing figures and tables. **T**he authors thank Stony Brook Research Computing and Cyberinfrastructure, and the Institute for Advanced Computational Science (IACS) at Stony Brook University for access to the high-performance SeaWulf computing system, IACS staff also provided technical assistance critical for the completion of this work, with notable support from Firat Coşkun.

### 5.2. Funding

Research reported in this publication was supported by the National Institute of Mental Health of the National Institutes of Health (NIH) under award numbers K01MH107763 and R01MH120293 to JXVS, and F30MH122136 and to JCW. JCW was also supported by the Stony Brook University Medical Scientist Training Program (T32GM008444; PI: Michael A. Frohman). PNT was supported by the Stony Brook University Undergraduate Research and Creative Activities (URECA) Summer Research Program. The content is solely the responsibility of the authors and does not necessarily represent the official views of the NIH.

### 5.3. Data and code availability

The data employed here are, in part, publicly available from the Human Connectome Project (HCP) at https://db.humanconnectome.org (Hodge et al., 2016), and include all resting-state fMRI data from the *HCP 500 Subjects Release*, a subset of the working memory task data from the *HCP 500 Subjects Release,* and a subset of the resting-state fMRI data from the *HCP 1200 Subjects Release* (Barch et al., 2013; Glasser et al., 2013; Smith et al., 2013; Ugurbil et al., 2013; Van Essen et al., 2013), as described in *Section 2.1.1.1* and *Supplementary Section S1.1.1*. Data were provided by the Human Connectome Project, WU-Minn Consortium (Principal Investigators: David Van Essen and Kamil Ugurbil; U54MH091657) funded by the 16 NIH Institutes and Centers that support the NIH Blueprint for Neuroscience Research; and by the McDonnell Center for Systems Neuroscience at Washington University.

The software package accompanying this article allows a user to perform the LPF-FD and GEV-DV volume censoring methods described herein, and determine optimal volume censoring parameters in their own datasets. This code is publicly available under the terms of the GNU General Public License Version 3 MathWorks File Exchange at https://www.mathworks.com/matlabcentral/fileexchange/98949-mcot and GitHub at https://github.com/MMTI/MCOT.

### 5.4. Author contributions

Conceptualization, J.C.W. and J.X.V.S.; Data Curation, J.C.W. and J.X.V.S.; Formal Analysis, J.C.W., P.N.T., and J.X.V.S.; Funding Acquisition, J.C.W. and J.X.V.S.; Investigation, J.C.W., P.N.T., and J.X.V.S; Methodology, J.C.W., P.N.T., J.R.L., and J.X.V.S.; Project Administration, J.C.W. and J.X.V.S.; Resources, J.X.V.S.; Software, J.C.W., P.N.T., J.R.L., and J.X.V.S.; Supervision, J.X.V.S.; Validation, J.C.W., P.N.T., J.R.L., and J.X.V.S.; Visualization, J.C.W., P.N.T., and J.X.V.S.; Writing – Original Draft, J.C.W., P.N.T., J.R.L., and J.X.V.S.; Writing – Review & Editing, J.C.W., P.N.T., J.R.L., and J.X.V.S.

## Abbreviations

ABCD: Adolescent Brain Cognitive Development
BOLD: blood-oxygen-level-dependent
DQM: data quality metric;
DV: temporal derivative root-mean-squared over voxels
FD: framewise displacement
FH+: positive for a reported family history of psychiatric or neurological disease
FH-: negative for a reported family history of psychiatric or neurological disease
fMRI: functional magnetic resonance imaging
GEV: generalized extreme value
GEV-DV: generalized extreme value low-pass filtered temporal derivative root-mean-squared over voxels
GLM: generalized linear model
GSR: global signal regression
HCP: Human Connectome Project
HCP500: Human Connectome Project 500 Subjects Release
LPF: low-pass filtered
LPF-DV: low-pass filtered temporal derivative root-mean-squared over voxels
LPF-FD: low-pass filtered framewise displacement
MAC-RSFC: mean absolute change in resting state functional connectivity
mFD: mean framewise displacement
MP: motion parameter
MSE: mean squared error
NYSPI: New York State Psychiatric Institute
QC: quality control
QC-FC: quality control-functional connectivity
ROI: region of interest
RSFC: resting state functional connectivity
SQM: subject-level quality control metric
SU+: tested positive for substance (drug or alcohol) use on the day of a scan session
SU-: tested negative for substance (drug or alcohol) use on the day of a scan session
ΔMSE-RSFC: delta mean squared error-resting state functional connectivity

## References

Afyouni, S., Nichols, T.E., 2018. Insight and inference for DVARS. Neuroimage 172, 291–312.

Aquino, K.M., Fulcher, B.D., Parkes, L., Sabaroedin, K., Fornito, A., 2020. Identifying and removing widespread signal deflections from fMRI data: Rethinking the global signal regression problem. Neuroimage 212, 116614.

Barch, D.M., Burgess, G.C., Harms, M.P., Petersen, S.E., Schlaggar, B.L., Corbetta, M., Glasser, M.F., Curtiss, S., Dixit, S., Feldt, C., Nolan, D., Bryant, E., Hartley, T., Footer, O., Bjork, J.M., Poldrack, R., Smith, S., Johansen-Berg, H., Snyder, A.Z., Van Essen, D.C., Consortium, W.U.-M.H., 2013. Function in the human connectome: task-fMRI and individual differences in behavior. Neuroimage 80, 169–189.

Bolton, T.A.W., Kebets, V., Glerean, E., Zoller, D., Li, J., Yeo, B.T.T., Caballero-Gaudes, C., Van De Ville, D., 2020. Agito ergo sum: Correlates of spatio-temporal motion characteristics during fMRI. Neuroimage 209, 116433.

Burgess, G.C., Kandala, S., Nolan, D., Laumann, T.O., Power, J.D., Adeyemo, B., Harms, M.P., Petersen, S.E., Barch, D.M., 2016. Evaluation of Denoising Strategies to Address Motion-Correlated Artifacts in Resting-State Functional Magnetic Resonance Imaging Data from the Human Connectome Project. Brain Connect 6, 669–680.

Caballero-Gaudes, C., Reynolds, R.C., 2017. Methods for cleaning the BOLD fMRI signal. Neuroimage 154, 128–149.

Casey, B.J., Cannonier, T., Conley, M.I., Cohen, A.O., Barch, D.M., Heitzeg, M.M., Soules, M.E., Teslovich, T., Dellarco, D.V., Garavan, H., Orr, C.A., Wager, T.D., Banich, M.T., Speer, N.K., Sutherland, M.T., Riedel, M.C., Dick, A.S., Bjork, J.M., Thomas, K.M., Chaarani, B., Mejia, M.H., Hagler, D.J., Jr., Daniela Cornejo, M., Sicat, C.S., Harms, M.P., Dosenbach, N.U.F., Rosenberg, M., Earl, E., Bartsch, H., Watts, R., Polimeni, J.R., Kuperman, J.M., Fair, D.A., Dale, A.M., Workgroup, A.I.A., 2018. The Adolescent Brain Cognitive Development (ABCD) study: Imaging acquisition across 21 sites. Dev Cogn Neurosci 32, 43–54.

Ciric, R., Wolf, D.H., Power, J.D., Roalf, D.R., Baum, G.L., Ruparel, K., Shinohara, R.T., Elliott, M.A., Eickhoff, S.B., Davatzikos, C., Gur, R.C., Gur, R.E., Bassett, D.S., Satterthwaite, T.D., 2017. Benchmarking of participant-level confound regression strategies for the control of motion artifact in studies of functional connectivity. Neuroimage 154, 174–187.

Efron, B., Tibshirani, R., 1993. An introduction to the bootstrap. Chapman & Hall, New York.

Fair, D.A., Miranda-Dominguez, O., Snyder, A.Z., Perrone, A., Earl, E.A., Van, A.N., Koller, J.M., Feczko, E., Tisdall, M.D., van der Kouwe, A., Klein, R.L., Mirro, A.E., Hampton, J.M., Adeyemo, B., Laumann, T.O., Gratton, C., Greene, D.J., Schlaggar, B.L., Hagler, D.J., Jr., Watts, R., Garavan, H., Barch, D.M., Nigg, J.T., Petersen, S.E., Dale, A.M., Feldstein-Ewing, S.W., Nagel, B.J., Dosenbach, N.U.F., 2020. Correction of respiratory artifacts in MRI head motion estimates. Neuroimage 208, 116400.

Fox, M.D., Zhang, D., Snyder, A.Z., Raichle, M.E., 2009. The global signal and observed anticorrelated resting state brain networks. J Neurophysiol 101, 3270–3283.

Glasser, M.F., Coalson, T.S., Bijsterbosch, J.D., Harrison, S.J., Harms, M.P., Anticevic, A., Van Essen, D.C., Smith, S.M., 2018. Using temporal ICA to selectively remove global noise while preserving global signal in functional MRI data. Neuroimage 181, 692–717.

Glasser, M.F., Sotiropoulos, S.N., Wilson, J.A., Coalson, T.S., Fischl, B., Andersson, J.L., Xu, J., Jbabdi, S., Webster, M., Polimeni, J.R., Van Essen, D.C., Jenkinson, M., Consortium, W.U.-M.H., 2013. The minimal preprocessing pipelines for the Human Connectome Project. Neuroimage 80, 105–124.

Glover, G.H., Li, T.Q., Ress, D., 2000. Image-based method for retrospective correction of physiological motion effects in fMRI: RETROICOR. Magn Reson Med 44, 162–167.

Gratton, C., Dworetsky, A., Coalson, R.S., Adeyemo, B., Laumann, T.O., Wig, G.S., Kong, T.S., Gratton, G., Fabiani, M., Barch, D.M., Tranel, D., Miranda-Dominguez, O., Fair, D.A., Dosenbach, N.U.F., Snyder, A.Z., Perlmutter, J.S., Petersen, S.E., Campbell, M.C., 2020. Removal of high frequency contamination from motion estimates in single-band fMRI saves data without biasing functional connectivity. Neuroimage 217, 116866.

Griffanti, L., Salimi-Khorshidi, G., Beckmann, C.F., Auerbach, E.J., Douaud, G., Sexton, C.E., Zsoldos, E., Ebmeier, K.P., Filippini, N., Mackay, C.E., Moeller, S., Xu, J., Yacoub, E., Baselli, G., Ugurbil, K., Miller, K.L., Smith, S.M., 2014. ICA-based artefact removal and accelerated fMRI acquisition for improved resting state network imaging. Neuroimage 95, 232–247.

Hagler, D.J., Jr., Hatton, S., Cornejo, M.D., Makowski, C., Fair, D.A., Dick, A.S., Sutherland, M.T., Casey, B.J., Barch, D.M., Harms, M.P., Watts, R., Bjork, J.M., Garavan, H.P., Hilmer, L., Pung, C.J., Sicat, C.S., Kuperman, J., Bartsch, H., Xue, F., Heitzeg, M.M., Laird, A.R., Trinh, T.T., Gonzalez, R., Tapert, S.F., Riedel, M.C., Squeglia, L.M., Hyde, L.W., Rosenberg, M.D., Earl, E.A., Howlett, K.D., Baker, F.C., Soules, M., Diaz, J., de Leon, O.R., Thompson, W.K., Neale, M.C., Herting, M., Sowell, E.R., Alvarez, R.P., Hawes, S.W., Sanchez, M., Bodurka, J., Breslin, F.J., Morris, A.S., Paulus, M.P., Simmons, W.K., Polimeni, J.R., van der Kouwe, A., Nencka, A.S., Gray, K.M., Pierpaoli, C., Matochik, J.A., Noronha, A., Aklin, W.M., Conway, K., Glantz, M., Hoffman, E., Little, R., Lopez, M., Pariyadath, V., Weiss, S.R., Wolff-Hughes, D.L., DelCarmen-Wiggins, R., Feldstein Ewing, S.W., Miranda-Dominguez, O., Nagel, B.J., Perrone, A.J., Sturgeon, D.T., Goldstone, A., Pfefferbaum, A., Pohl, K.M., Prouty, D., Uban, K., Bookheimer, S.Y., Dapretto, M., Galvan, A., Bagot, K., Giedd, J., Infante, M.A., Jacobus, J., Patrick, K., Shilling, P.D., Desikan, R., Li, Y., Sugrue, L., Banich, M.T., Friedman, N., Hewitt, J.K., Hopfer, C., Sakai, J., Tanabe, J., Cottler, L.B., Nixon, S.J., Chang, L., Cloak, C., Ernst, T., Reeves, G., Kennedy, D.N., Heeringa, S., Peltier, S., Schulenberg, J., Sripada, C., Zucker, R.A., Iacono, W.G., Luciana, M., Calabro, F.J., Clark, D.B., Lewis, D.A., Luna, B., Schirda, C., Brima, T., Foxe, J.J., Freedman, E.G., Mruzek, D.W., Mason, M.J., Huber, R., McGlade, E., Prescot, A., Renshaw, P.F., Yurgelun-Todd, D.A., Allgaier, N.A., Dumas, J.A., Ivanova, M., Potter, A., Florsheim, P., Larson, C., Lisdahl, K., Charness, M.E., Fuemmeler, B., Hettema, J.M., Maes, H.H., Steinberg, J., Anokhin, A.P., Glaser, P., Heath, A.C., Madden, P.A., Baskin-Sommers, A., Constable, R.T., Grant, S.J., Dowling, G.J., Brown, S.A., Jernigan, T.L., Dale, A.M., 2019. Image processing and analysis methods for the Adolescent Brain Cognitive Development Study. Neuroimage, 116091.

Hodge, M.R., Horton, W., Brown, T., Herrick, R., Olsen, T., Hileman, M.E., McKay, M., Archie, K.A., Cler, E., Harms, M.P., Burgess, G.C., Glasser, M.F., Elam, J.S., Curtiss, S.W., Barch, D.M., Oostenveld, R., Larson-Prior, L.J., Ugurbil, K., Van Essen, D.C., Marcus, D.S., 2016. ConnectomeDB--Sharing human brain connectivity data. Neuroimage 124, 1102–1107.

Jernigan, T.L., Brown, S.A., Coordinators, A.C., 2018. Introduction. Dev Cogn Neurosci 32, 1–3.

Kassinopoulos, M., Mitsis, G.D., 2019. Identification of physiological response functions to correct for fluctuations in resting-state fMRI related to heart rate and respiration. Neuroimage 202, 116150.

Lindquist, M.A., Geuter, S., Wager, T.D., Caffo, B.S., 2019. Modular preprocessing pipelines can reintroduce artifacts into fMRI data. Hum Brain Mapp 40, 2358–2376.

Miller, K.L., Alfaro-Almagro, F., Bangerter, N.K., Thomas, D.L., Yacoub, E., Xu, J., Bartsch, A.J., Jbabdi, S., Sotiropoulos, S.N., Andersson, J.L., Griffanti, L., Douaud, G., Okell, T.W., Weale, P., Dragonu, I., Garratt, S., Hudson, S., Collins, R., Jenkinson, M., Matthews, P.M., Smith, S.M., 2016. Multimodal population brain imaging in the UK Biobank prospective epidemiological study. Nat Neurosci 19, 1523–1536.

Moeller, S., Yacoub, E., Olman, C.A., Auerbach, E., Strupp, J., Harel, N., Ugurbil, K., 2010. Multiband multislice GE-EPI at 7 tesla, with 16-fold acceleration using partial parallel imaging with application to high spatial and temporal whole-brain fMRI. Magn Reson Med 63, 1144–1153.

Murphy, K., Birn, R.M., Handwerker, D.A., Jones, T.B., Bandettini, P.A., 2009. The impact of global signal regression on resting state correlations: are anti-correlated networks introduced? Neuroimage 44, 893–905.

Murphy, K., Fox, M.D., 2017. Towards a consensus regarding global signal regression for resting state functional connectivity MRI. Neuroimage 154, 169–173.

Muschelli, J., Nebel, M.B., Caffo, B.S., Barber, A.D., Pekar, J.J., Mostofsky, S.H., 2014. Reduction of motion-related artifacts in resting state fMRI using aCompCor. Neuroimage 96, 22–35.

NIMH, 2020. National Institute of Mental Health Strategic Plan for Research. https://www.nimh.nih.gov/about/strategic-planning-reports/.

Parkes, L., Fulcher, B., Yucel, M., Fornito, A., 2018. An evaluation of the efficacy, reliability, and sensitivty of motion correction strategies for resting-state functional MRI. Neuroimage 171, 415–436.

Power, J.D., Barnes, K.A., Snyder, A.Z., Schlaggar, B.L., Petersen, S.E., 2012. Spurious but systematic correlations in functional connectivity MRI networks arise from subject motion. Neuroimage 59, 2142–2154.

Power, J.D., Cohen, A.L., Nelson, S.M., Wig, G.S., Barnes, K.A., Church, J.A., Vogel, A.C., Laumann, T.O., Miezin, F.M., Schlaggar, B.L., Petersen, S.E., 2011. Functional network organization of the human brain. Neuron 72, 665–678.

Power, J.D., Lynch, C.J., Adeyemo, B., Petersen, S.E., 2020. A Critical, Event-Related Appraisal of Denoising in Resting-State fMRI Studies. Cereb Cortex.

Power, J.D., Lynch, C.J., Silver, B.M., Dubin, M.J., Martin, A., Jones, R.M., 2019a. Distinctions among real and apparent respiratory motions in human fMRI data. Neuroimage 201, 116041.

Power, J.D., Mitra, A., Laumann, T.O., Snyder, A.Z., Schlaggar, B.L., Petersen, S.E., 2014. Methods to detect, characterize, and remove motion artifact in resting state fMRI. Neuroimage 84, 320–341.

Power, J.D., Plitt, M., Laumann, T.O., Martin, A., 2017. Sources and implications of whole-brain fMRI signals in humans. Neuroimage 146, 609–625.

Power, J.D., Schlaggar, B.L., Petersen, S.E., 2015. Recent progress and outstanding issues in motion correction in resting state fMRI. Neuroimage 105, 536–551.

Power, J.D., Silver, B.M., Silverman, M.R., Ajodan, E.L., Bos, D.J., Jones, R.M., 2019b. Customized head molds reduce motion during resting state fMRI scans. Neuroimage 189, 141–149.

Prescott, P., Walden, A.T., 1980. Maximum likelihood estimation of the parameters of the generalized extreme-value distrubtion. Biometrika 67, 723–724.

Pruim, R.H.R., Mennes, M., Buitelaar, J.K., Beckmann, C.F., 2015. Evaluation of ICA-AROMA and alternative strategies for motion artifact removal in resting state fMRI. Neuroimage 112, 278–287.

Saad, Z.S., Gotts, S.J., Murphy, K., Chen, G., Jo, H.J., Martin, A., Cox, R.W., 2012. Trouble at rest: how correlation patterns and group differences become distorted after global signal regression. Brain Connect 2, 25–32.

Satterthwaite, T.D., Elliott, M.A., Gerraty, R.T., Ruparel, K., Loughead, J., Calkins, M.E., Eickhoff, S.B., Hakonarson, H., Gur, R.C., Gur, R.E., Wolf, D.H., 2013. An improved framework for confound regression and filtering for control of motion artifact in the preprocessing of resting-state functional connectivity data. Neuroimage 64, 240–256.

Satterthwaite, T.D., Wolf, D.H., Loughead, J., Ruparel, K., Elliott, M.A., Hakonarson, H., Gur, R.C., Gur, R.E., 2012. Impact of in-scanner head motion on multiple measures of functional connectivity: relevance for studies of neurodevelopment in youth. Neuroimage 60, 623–632.

Siegel, J.S., Mitra, A., Laumann, T.O., Seitzman, B.A., Raichle, M., Corbetta, M., Snyder, A.Z., 2017. Data Quality Influences Observed Links Between Functional Connectivity and Behavior. Cereb Cortex 27, 4492–4502.

Smith, S.M., Beckmann, C.F., Andersson, J., Auerbach, E.J., Bijsterbosch, J., Douaud, G., Duff, E., Feinberg, D.A., Griffanti, L., Harms, M.P., Kelly, M., Laumann, T., Miller, K.L., Moeller, S., Petersen, S., Power, J., Salimi-Khorshidi, G., Snyder, A.Z., Vu, A.T., Woolrich, M.W., Xu, J., Yacoub, E., Ugurbil, K., Van Essen, D.C., Glasser, M.F., Consortium, W.U.-M.H., 2013. Resting-state fMRI in the Human Connectome Project. Neuroimage 80, 144–168.

Smyser, C.D., Snyder, A.Z., Neil, J.J., 2011. Functional connectivity MRI in infants: exploration of the functional organization of the developing brain. Neuroimage 56, 1437–1452.

Tagliazucchi, E., Laufs, H., 2014. Decoding wakefulness levels from typical fMRI resting-state data reveals reliable drifts between wakefulness and sleep. Neuron 82, 695–708.

Ugurbil, K., Xu, J., Auerbach, E.J., Moeller, S., Vu, A.T., Duarte-Carvajalino, J.M., Lenglet, C., Wu, X., Schmitter, S., Van de Moortele, P.F., Strupp, J., Sapiro, G., De Martino, F., Wang, D., Harel, N., Garwood, M., Chen, L., Feinberg, D.A., Smith, S.M., Miller, K.L., Sotiropoulos, S.N., Jbabdi, S., Andersson, J.L., Behrens, T.E., Glasser, M.F., Van Essen, D.C., Yacoub, E., Consortium, W.U.-M.H., 2013. Pushing spatial and temporal resolution for functional and diffusion MRI in the Human Connectome Project. Neuroimage 80, 80–104.

Van Dijk, K.R., Sabuncu, M.R., Buckner, R.L., 2012. The influence of head motion on intrinsic functional connectivity MRI. Neuroimage 59, 431–438.

Van Essen, D.C., Smith, S.M., Barch, D.M., Behrens, T.E., Yacoub, E., Ugurbil, K., Consortium, W.U.-M.H., 2013. The WU-Minn Human Connectome Project: an overview. Neuroimage 80, 62–79.

Volkow, N.D., Koob, G.F., Croyle, R.T., Bianchi, D.W., Gordon, J.A., Koroshetz, W.J., Perez-Stable, E.J., Riley, W.T., Bloch, M.H., Conway, K., Deeds, B.G., Dowling, G.J., Grant, S., Howlett, K.D., Matochik, J.A., Morgan, G.D., Murray, M.M., Noronha, A., Spong, C.Y., Wargo, E.M., Warren, K.R., Weiss, S.R.B., 2018. The conception of the ABCD study: From substance use to a broad NIH collaboration. Dev Cogn Neurosci 32, 4–7.

Wackerly, D.D., Mendenhall, W., Scheaffer, R.L., 2008. Mathematical statistics with applications, 7th ed. Thomson Brooks/Cole, Belmont, CA.

Xifra-Porxas, A., Kassinopoulos, M., Mitsis, G.D., 2021. Physiological and motion signatures in static and time-varying functional connectivity and their subject identifiability. Elife 10.

Yan, C.G., Cheung, B., Kelly, C., Colcombe, S., Craddock, R.C., Di Martino, A., Li, Q., Zuo, X.N., Castellanos, F.X., Milham, M.P., 2013. A comprehensive assessment of regional variation in the impact of head micromovements on functional connectomics. Neuroimage 76, 183–201.

Zeng, L.L., Wang, D., Fox, M.D., Sabuncu, M., Hu, D., Ge, M., Buckner, R.L., Liu, H., 2014. Neurobiological basis of head motion in brain imaging. Proc Natl Acad Sci U S A 111, 6058–6062.

