## Supplementary_Materials for "Advancing motion denoising of multiband resting-state functional connectivity fMRI data"

#### These supplementary materials include:

Supplementary Materials and Methods
Supplementary Results and Discussion
Supplementary Tables S1 – S4
Supplementary Figures S1 – S18
Supplementary References

### Abbreviations

ABCD, Adolescent Brain Cognitive Development;
BOLD, blood-oxygen-level-dependent;
DICOM, Digital Imaging and Communications in Medicine;
DQM, data quality metric;
DSM-IV, Diagnostic and Statistical Manual of Mental Disorders, Fourth Edition;
DSM-IV-TR, Diagnostic and Statistical Manual of Mental Disorders, Fourth Edition, Text Revision;
DV, temporal derivative root-mean-squared over voxels;
ECDF, empirical cumulative distribution function;
FD, framewise displacement;
FH+, positive for a reported family history of psychiatric or neurological disease;
FH-, negative for a reported family history of psychiatric or neurological disease;
fMRI, functional magnetic resonance imaging;
GEV, generalized extreme value;
GEV-DV, generalized extreme value low-pass filtered temporal derivative root-mean-squared over voxels;
GLM, generalized linear model;
GSR, global signal regression;
HCP, Human Connectome Project;
HCP500, Human Connectome Project 500 Subjects Release;
HCP1200, Human Connectome Project 1200 Subjects Release;
LPF, low-pass filtered;
LPF-DV, low-pass filtered temporal derivative root-mean-squared over voxels;
LPF-FD, low-pass filtered framewise displacement;
MAC-RSFC, mean absolute change in resting state functional connectivity;
mFD, mean framewise displacement;
MNI, Montreal Neurological Institute;
MNI152NLin6, Montreal Neurological Institute 152 non-linear 6th-generation space;
MP, motion parameter;
MSE, mean squared error;
MRI, magnetic resonance imaging;
NIfTI, Neuroimaging Informatics Technology Initiative;
NYSPI, New York State Psychiatric Institute;
QC, quality control;
QC-FC, quality control-functional connectivity;
ROI, region of interest
RSFC, resting state functional connectivity;
R-MPS, run-level motion pairing score;
SCID-I, Structured Clinical Interview for the DSM-IV Axis I Disorders;
S-MPS, subject-level motion pairing score;
SU+, tested positive for substance (drug or alcohol) use on the day of a scan session;
SU-, tested negative for substance (drug or alcohol) use on the day of a scan session;
ΔMSE-RSFC, delta mean squared error-resting state functional connectivity.

### S1. Supplementary Methods

*S1.1. Functional magnetic resonance imaging datasets – supplementary details*

*S1.1.1. Supplementary Human Connectome Project datasets*

*S1.1.1.1. Resting-state and working memory task data*

To evaluate the effects of volume censoring on the relationships between task and resting-state data across sessions (see Supplementary *Section S1.6.1*) as a function of the percentage of data removed by volume censoring, we utilized minimally preprocessed (Glasser et al., 2013) resting-state and working memory task data from the Human Connectome Project (Barch et al., 2013; Smith et al., 2013; Ugurbil et al., 2013; Van Essen et al., 2013) 500 Subjects Release (HCP500), comprising four 1200 volume resting-state runs (collected across two sessions) and two 405 volume working memory task runs (collected during session 1) from each participant. Only subjects with complete sets of resting-state data and 2 full runs of working memory data were included in this analysis, totaling 330 subjects. Both resting-state and working memory task data were processed and obtained as described in *Section 2.1.2*. Minimally preprocessed HCP working memory Task data were downloaded for each subject from the "Task Working Memory Task fMRI Preprocessed" package on db.humanconnectome.org (Hodge et al., 2016). working memory Task BOLD data were obtained from the files, *tfMRI_WM_LR.nii.gz* and *tfMRI_WM_RL.nii.gz*; motion parameters were obtained from the file, *Movement_Regressors.txt* corresponding to each run. Minimally preprocessed HCP resting-state data was obtained as described in *Section 2.1.2.*

*S1.1.1.2. Resting-state data*

To assess the impact of censoring on the association between observed intra-session differences in RSFC correlations observed for each subject and the difference in the time of day that each scan session took place (see *Supplementary Section S1.6.2)*, we utilized a subset of the HCP500 minimally preprocessed resting-state dataset produced by selecting only subjects with 4 complete runs of resting-state data (n = 478), collected over 2 sessions of 2 runs each. Minimally preprocessed HCP resting-state data was obtained as described in *Section 2.1.2.*

*S1.1.2. New York State Psychiatric Institute (NYSPI) resting-state dataset*

*S1.1.2.1. Acquisition and preprocessing*

The NYSPI Dataset comprises 4 resting-state functional blood-oxygen-level-dependent (BOLD) T2* runs of 15 minutes each from 26 healthy subjects acquired at the New York State Psychiatric Institute (NYSPI; New York, NY) using a General Electric (Boston, MA) MR750 3T scanner (multiband factor 6, repetition time = 850 ms, echo time = 25 ms, flip angle = 60°, voxel size = 2 mm isotropic). BOLD runs were acquired in alternating anterior-posterior and posterior-anterior phase encode directions. Additionally, for each subject, two T1 weighted and one T2 weighted images were acquired, as well as one $B_0$ field map and two 3-volume spin echo images in opposing phase encode directions (anterior-posterior and posterior-anterior) for susceptibility distortion correction. Structural and functional images were converted from DICOM to NIfTI-1 format using dcm2nii (https://people.cas.sc.edu/rorden/mricron/dcm2nii.html).

Images were preprocessed using the following steps of the HCP Minimal Preprocessing Pipelines v3.20.0 (https://github.com/Washington-University/HCPpipelines; described in detail by Glasser and colleagues (2013): 1) *PreFreeSurfer*, 2) *FreeSurfer*, 3) *PostFreeSurfer* and 4) *fMRIVolume* Surface registration was completed by cortical surface matching (MSMsulc; Robinson et al., 2018; Robinson et al., 2014) and slice timing correction was disabled. Preprocessed structural (T1w) and functional (BOLD) images obtained from *fMRIVolume* in Montreal Neurological Institute (MNI) 152 non-linear 6th-generation space (MNI152NLin6; Grabner et al., 2006) with 2mm isotropic voxels were used for all subsequent processing and analysis.

*S1.1.2.2. Sample characteristics*

Inclusion criteria were: 1) absence of any current Axis-I psychiatric disorder per the Diagnostic and Statistical Manual of Mental Disorders, Fourth Edition, Text Revision (DSM-IV-TR; American Psychiatric Association, 2000), as evaluated through psychiatric evaluation and administration of the Structured Clinical Interview for the DSM-IV Axis I Disorders (SCID-I; First and Gibbon, 2004), 2) age 18-55 years old, and 3) fluent English speaking ability. Exclusion criteria were: 1) any current drug use disorder or major affective disorder, evaluated through psychiatric evaluation and administration if the SCID-I; 2) any medical history of neurological disorders (including head trauma with loss of consciousness, epilepsy, or other clinically significant disorders of the central nervous system), intellectual disability, hearing impairment, or unstable severe medical conditions; 3) claustrophobia; and 4) metal implants or paramagnetic objects contained within the body which may interfere with the MRI scan (Shellock, 1998). Demographics for this sample are shown in **Table S1**.

*S1.2. Standard quality control benchmarks across a full range of censoring thresholds*

*S1.2.1. QC-FC and median absolute QC-FC*

QC-FC (Quality Control – Functional Connectivity) for ROI pair *k* is defined as the Pearson product-moment correlation between Z-transformed ROI pair correlations and a summary measure of subject in-scanner motion (mean FD):

$$QC\text{-}FC_{Z,mFD_k} = corr_{Pearson}\left(Z_{i,k}, mFD_i\right), \quad (1)$$

where $Z_{i,j,k}$ is the RSFC correlation for subject *i* and ROI pair *k* (averaged across all runs), and $mFD_i$ is the mean FD value for subject *i,* averaged across all available runs for that subject. Median absolute QC-FC, shown in **Figure 1A**, is defined as the median (over ROI pairs) absolute value of all QC-FC correlations, as defined by Ciric and associates (2017):

$$Median\ absolute\ QC\text{-}FC = median\left(\left|QC\text{-}FC_{Z,mFD}\right|\right). \quad (2)$$

*S1.2.2. QC-FC distance correlation and null rejection rate*

QC-FC null rejection rate, shown in **Figure 1B**, is defined as the proportion of all QC-FC correlations that are statistically significant (Benjamini and Hochberg, 1995; Ciric et al., 2017), as calculated in **Equation 1**, after false discovery rate (FDR) correction for multiple comparisons (Benjamini and Hochberg, 1995). QC-FC Distance Correlation (Spearman), shown in **Figure 1C**, was calculated as the Spearman's rank-order correlation coefficient between QC-FC calculated in **Equation 1** and the distance between ROI pairs (Ciric et al., 2017; Muschelli et al., 2014; Parkes et al., 2018; Power et al., 2012; Satterthwaite et al., 2013):

$$QC\text{-}FC\ Distance\ Correlation = corr_{Spearman}\left(QC\text{-}FC_{Z,mFD}, dist\right), \quad (3)$$

where $dist$ is the Euclidean distance between ROIs comprising ROI pair $k$.

*S1.2.3. High-low null rejection rate*

To calculate High-Low Null Rejection Rate as shown in **Figure 1D**, participants were first separated into terciles based on mean FD with upper and lower terciles representing high- and low-motion participants, respectively (Burgess et al., 2016; Power et al., 2014; Pruim et al., 2015; Satterthwaite et al., 2013; Van Dijk et al., 2012). A Welch's (unequal variances) two-sample t-test was performed between high- and low-motion participants for each ROI pair *k* and corrected for multiple comparisons with significance threshold $\alpha = 0.000432$, equivalent to 300 times the Bonferroni-corrected significance threshold for 34,716 ROI pairs derived from 264 ROIs, following Burgess and associates (Burgess et al., 2016).

*S1.2.4. Two-step generalized linear model-based DQMs*

Burgess and associates also employed a two-level generalized linear model (GLM) to quantify mean and distance-dependent changes in RSFC and QC-FC correlations resulting from denoising (Burgess et al., 2016). Here, we adapted these methods to quantify the mean and distance-dependence of RSFC correlations (rather than changes therein due to censoring), as well as the relationship of each with mFD, in *HCP Dataset 1a*. Both the first and second level GLMs are simple linear regressions of the form, $Y = \beta_0 + \beta_1 X$, where $Y$ is a specified dependent variable, $X$ is a specified predictor, and $\beta_0$ and $\beta_1$ are the regression coefficients. All predictors are mean-centered before coefficients are estimated, so that $\beta_0$ estimates the mean of $Y$. Coefficients were estimated using first-level GLMs for each subject, where $Y$ is that subject's RSFC correlations, and $X$ is the distance between ROI pairs. In this framework, each $\beta_0$ is each subject's mean ROI pair correlation, and $\beta_1$ is the distance-dependence component of these ROI pair correlations.

We then used second-level GLMs to determine the relationship between these coefficients and a summary measure of subject motion (i.e., mean FD). Both models use (mean-centered) mFD as the $X$ variable, and then each model uses either $\beta_0$ or $\beta_1$ from the first-level model as its $Y$ variable. Thus, in the model using $\beta_0$, the intercept parameter is simply the mean RSFC correlation across all subjects and ROIs **(Figure 1G)**, while the slope parameter is a measure of QC-FC (i.e., the slope of the relationship between each subject's mean RSFC and their mFD; **Figure 1E**). In contrast, in the model using $\beta_1$ from the first-level model, the intercept parameter indicates the average within-subject distance-dependence **(Figure 1H)**, while the slope parameter indicates the association of within-subject distance-dependence with subject motion (mFD; **Figure 1F**).

Where possible, we identified optimal censoring thresholds as determined by each of these DQMs in the range of 0% to 70% volumes censored (total across the dataset), in *HCP Dataset 1a*. We defined these optima as the value which produces the minimum magnitude of the DQM (i.e., closest to zero), or selected the first zero-crossing if multiple optima were possible in that range, for: median absolute QC-FC, QC-FC null rejection rate, QC-FC distance correlation (calculated via Spearman's rank-order correlation), high-low null rejection rate, mean QC-FC (via GLM), QC-FC slope (via GLM), mean RSFC (via GLM), and the distance-dependence of RSFC correlations (via GLM). We also noted, for variance and ΔMSE-RSFC, the point at which censoring achieves the greatest magnitude reduction (i.e., most negative change) in these statistics.

*S1.3 Comparison of high- to low-motion participants after censoring: Monte Carlo resampling procedure*

To evaluate the effect that FH+ and SU+ subjects have on detected differences between high- and low-motion subjects, 1000 Monte Carlo resamplings of *HCP Dataset 1* were generated by randomly removing FH- and SU- subjects (only), with approximately the same mean motion as the FH+ and SU+ subjects that were removed to create *HCP Dataset 1b*. This was done by iteratively restricting the set of participants from which resampling could occur to those with a median FD value across all RSFC runs that was either higher or lower than the "true" excluded sample mean (i.e., of the FH+ and SU+ subjects), depending on whether the current sample mean was higher or lower than the mean of the true sample. That is, for each Monte Carlo iteration, a new set of 187 participants to be removed was randomly selected, one participant at a time. The first participant to be removed was randomly selected from the entire set of FH- and SU- participants. If this participant's median FD value was greater than the mean of the 187 excluded FH+ and SU+ participants, then the second participant was randomly selected from only those participants with a median FD value lower than the mean of the true 187 participants; conversely, if the participant's median FD value was lower than the mean of the true sample, then the second participant had to have a median value that was higher than the mean. The third participant was then selected from those participants with higher or lower median FDs than the true sample mean, based on whether the mean of the first two participants was higher or lower than the true sample mean. This process was repeated iteratively until a full sample of 187 participants had been constructed, for each of the 1,000 Monte Carlo resamplings.

*S1.4 Evaluation of trait confounds in DQMs using datasets balanced by subject motion: Matching algorithms and FH+ status as a nuisance parameter*

This procedure began by dividing subjects into one subset with a positive family history of psychiatric or neurological disease (FH+), and another subset with a negative family history of psychiatric or neurological disease (FH-). Next, we generated a single value (using one of three methods described below) to quantify the similarity between the motion characteristics of any possible subject pairing, herein referred to as a subject-level motion pairing score (S-MPS). Greater S-MPS values indicate greater difference in motion characteristics between a pair of subjects. As each subject's data comprises multiple runs, generating S-MPS values requires first generating run-level motion pairing scores (R-MPS) for each possible pairing of runs between any two subjects. We show results from three separate algorithms for producing R-MPS, which are detailed later in this section.

The resulting 16 R-MPS values (because each subject has 4 runs, producing 16 pairwise combinations) for each subject pairing can be represented as a 4x4 matrix, from which we aimed to select a single set of four run pairings (using each row and column from the 4x4 matrix only once each) that minimizes the sum of R-MPS scores. Using the MATLAB function *matchpairs* (an implementation of the Duff-Koster algorithm to solve the linear assignment problem; Duff and Koster, 2001) we derived a single S-MPS from each 4x4 R-MPS matrix. The four R-MPS from the optimal run pairings were then summed to produce the S-MPS for that subject pair. Next, the S-MPS for each subject pair were entered into a ranking matrix containing a row for each FH+ subject and a column for each FH- subject. The ranking matrix contains a single S-MPS for every possible pairing of an FH+ subject to an FH- subject, such that the value in row *i* and column *j* is the S-MPS quantifying the difference in motion characteristics between FH+ subject *i* and FH- subject *j*. Next, we applied *matchpairs* to the ranking matrix to obtain the optimal subject-to-subject pairings, where optimality is defined as the minimization of the total sum of S-MPS, across all FH+ subjects without replacement.

Motion-matching was performed with three methods used to calculate run-level motion pairing scores (R-MPS). The first method, used to produce *HCP Dataset 2a*, involves the calculation of the two-sample Cramér–von Mises criterion (Anderson, 1962) for each run pairing. The Cramér–von Mises criterion is a score computed via the integration of the squared difference of two empirical cumulative distribution functions (ECDFs). We calculated ECDFs for the magnitude of the derivatives of each of the 6 MPs, separately. The Cramér–von Mises criterion was then calculated once for each MP's ECDF. The R-MPS between those runs was calculated as the average of these criteria. The second method, used to produce *HCP Dataset 2b*, is identical to the first, but additionally involves the calculation of the Cramér–von Mises criterion for the magnitude of the DV vectors for each run pairing. Both the averaged MP R-MPS and the DV R-MPS were then individually z-score normalized (over the distribution of all possible subject pairings) to control for magnitude differences, and averaged (with all 6 MPs having equal weight to DV) to generate S-MPS scores. The third and final method, used to produce *HCP Dataset 2c*, compares runs using their MPs. The absolute value of the derivatives of MPs (by backwards differences) was averaged for each of the two runs being compared. We calculated the magnitude (absolute value) of the difference in the two resulting means for each of the six MPs, which was then averaged over all six MPs to calculate a single R-MPS for that run pairing.

We also calculated five of the DQMs described in section 2.2.1 on these datasets, but while also using a dichotomous nuisance regressor controlling for FH+ status, described in the following. QC-FC Slope (Spearman's ρ), QC-FC Null Rejection Rate, and Median Absolute QC-FC were calculated normally, with the exception that QC-FC correlations were calculated as a partial correlation between ROI pair correlations and mean FD (mean centered), while controlling for FH+ status (also mean centered). Similarly, QC-FC Slope and Mean QC-FC were calculated using the two-level GLM-based approach (Burgess et al., 2016), with the addition of a variable accounting for FH+ status (mean centered) and an interaction term between FH+ status and mean FD. Null distributions were generated for all five DQMs using 10,000 random permutations of FH+ membership, and p-values were obtained for each observed statistic by comparison to the relevant null distribution.

*S1.5. Two-parameter global optimization of parameters for combined LPF-FD and GEV-DV volume censoring*

Optimal parameters for combined LPF-FD and GEV-DV censoring were determined with an optimization procedure minimizing ΔMSE-RSFC in *HCP Dataset 1,* calculated as described in Section *2.3.2.2.* We used a particle swarm global optimization algorithm (*particleswarm* in MATLAB; Kennedy and Eberhart, 1995; Mezura-Montes and Coello Coello, 2011; Shi and Eberhart, 1998), followed by simulated annealing (*simulannealbnd*; Ingber, 1996; Kirkpatrick et al., 1983; Metropolis et al., 1953), each followed by a pattern search local optimization algorithm (*patternsearch* with default settings*;* Abramson et al., 2009; Audet and Dennis, 2002; Conn et al., 1997; Davidon, 1991; Kolda et al., 2003; Lewis et al., 2007) to refine results. Particle swarm optimization used 44 particles, 1000 stall iterations and an initial neighborhood size of 1. 40 particles were randomly generated upon initialization in the interval $[10^{-1}, 10^{4}]$, with a $\log_{10}$-uniform distribution, for both LPF-FD $\Phi_F$ and GEV-DV $d_G$. The 4 remaining particles were placed along the edges of this distribution, i.e. $(10^{-1}, 10^{-1})$, $(10^{-1}, 10^{4})$, $(10^{4}, 10^{-1})$, and $(10^{4}, 10^{4})$. Simulated annealing used *particleswarm* optima (after refinement of local minima using *patternsearch*) as starting values, with an initial temperature of 1000, reannealing interval of 100, and function tolerance of $10^{-6}$, with length temperature and direction chosen uniformly at random (i.e., *'annealingfast'*).

*S1.6. Positive controls for RSFC volume censoring*

As discussed in detail in *Section 3.2.3* of the primary manuscript, one potential concern with denoising approaches (including volume censoring) for addressing motion artifact in RSFC is that they may remove true, non-artifactual, neural signals from the data, which may be difficult to detect without employing a positive control. While Glasser and colleagues (Glasser et al., 2018) have previously employed task-based analysis as a positive control for their temporal ICA denoising approach, such an approach is not suitable for volume censoring methods because of the large amounts of (often sequential) data that can be volume censoring. This is because censoring can remove large amounts, or even all, of the timeseries data necessary for obtaining parameter estimates in some subjects, which leads to dramatic increases in between-run and between-subject variability in task parameter estimates. This effect can be particularly pronounced when the peak of the HRF to a given event is largely censored, while the initial rise or return to baseline are not, which creates a scenario where a very small amount of noise in the measured data can lead to enormous errors in the estimate of the peak, resulting in extreme parameter values. Consequently, we developed and tested two different positive controls, described in *Supplementary Sections S1.6.1* and *S1.6.2*, below.

*S1.6.1. Impact of volume censoring on functional connectivity during resting and task states*

We aimed to determine the effect of volume censoring on functional connectivity correlations during resting- and task-states. First, we divided each subject's data (see *Section S1.1.1.1*) into three subsets: 1) session 1 resting-state data (here denoted $RSFC_{S1}$), 2) session 1 working memory task data ($WM_{S1}$), and 3) session 2 resting-state data ($RSFC_{S2}$). Next, we truncated the two $RSFC_{S1}$ runs, keeping only the first 405 volumes of each run, rendering them equal in length to the two $WM_{S1}$ runs; the duration of the $RSFC_{S2}$ runs were unmodified. Across the full range of volume censoring parameters (i.e., from 0% volumes censored to 100% volumes censored), we then calculate pairwise correlations between all 264 ROIs, as described in *Section 2.1.2*, for each of the 3 subsets (i.e., $RSFC_{S1}$, $WM_{S1}$, and $RSFC_{S2}$). Censoring was performed using the fixed ratio of LPF-FD $\Phi_F$ to GEV-DV $d_G$ determined to be optimal in the HCP500 dataset (see *Section 3.2.2.1*). Note that the $WM_{S1}$ data was subjected to the same exact processing as the RSFC data (see *Section 2.1.2*), and the same free parameters were used to censor all 3 subsets (and thus, the precise % of volumes censored in each dataset at each parameter value naturally differed to some extent).

For each subject, we then calculated an inter-session Pearson correlation (Z-Transformed), that reflects the correlation between sessions in the full connectivity graph (i.e., the extent to which the 34,716 pairwise correlations calculated within each session are correlated with each other), which was then averaged across subjects to obtain a single estimator, $Z_M$. Values of $Z_M$ were obtained 1) between $RSFC_{S1}$ and $RSFC_{S2}$, denoted $Z_{RSFC\text{-}RSFC}$; and 2) between $WM_{S1}$ and $RSFC_{S2}$, denoted $Z_{WM\text{-}RSFC}$. The change in $Z_M$ due to censoring relative to uncensored data, $\Delta Z_M$, was additionally calculated and as a function of percent volumes censored in the corresponding session 1 dataset (i.e., either $RSFC_{S1}$ or $WM_{S1}$ as appropriate), for visualization purposes. Over the course of this analysis, any subject from whom all data within a data subset ($RSFC_{S1}$, $WM_{S1}$, and $RSFC_{S2}$) was censored was excluded from all analyses using that subset's data for that parameter value.

*S1.6.2. Time of day effects on resting-state functional connectivity correlations*

Next, we evaluated the impact of increasingly aggressive volume censoring on time-of-day effects on resting-state correlations, in order to determine whether censoring high-motion frames results in improvement or impairment in the ability to detect this relationship, which could reflect removal of true neural signal by aggressive censoring methods. Across censoring parameters (ranging from 0% censored to 100% censored data, with censoring performed using the fixed ratio of LPF-FD $\Phi_F$ to GEV-DV $d_G$ optimal in the HCP500 dataset, as described in *Section 3.2.2.2*), we first calculated RSFC correlations for each subject separately within each session. We then took the difference between sessions (session 2 - session 1) for each ROI pair. This produced a single value for each ROI pair of each subject, reflecting the difference in RSFC between sessions for that ROI. Next, we calculated the change in scan time (in hours, reflecting the difference in the time of day, but not reflecting the number of days between sessions) for each subject (again, session 2 - session 1). For each ROI pair, we then determined the association between change in scan time and corresponding change in connectivity as the slope in a standard generalized linear model. Finally, we averaged these slopes over ROI pairs. For each set of censoring parameter values, 95% confidence intervals were estimated using a bias-corrected and accelerated bootstrap procedure (Efron and Tibshirani, 1993) using 10,000 bootstrap resamplings. During censoring, any subject from whom both runs was removed for either session was entirely excluded from analysis for that volume censoring parameter value.

### S2. Supplementary Results and Discussion

#### *S2.1. Evaluation of trait confounds in DQMs using datasets balanced by subject motion*

**Figure S11** demonstrates that the high-low null rejection rate is significantly higher in samples with FH+ remaining than in those with all FH+ removed across all P-value thresholds examined. We repeated these analyses with FH- subjects instead matched by 1) averaging the overlap in ECDFs of the derivatives of MPs together with the overlap in the ECDFs of DV traces (**Figure S12**), and 2) the means of the absolute values of the derivates of MP traces (**Figure S13;** see *Section 2.2.2.2* for details of these analyses).

Finally, we also used a matched FH+ and FH- dataset from these analyses and measured the effect of accounting for FH+ group membership in a partial correlation when calculating 1) median absolute QC-FC, 2) proportion of significant QC-FC correlations, and 3) QC-FC slope (Spearman's), and in a parallel analysis where FH+ was a nuisance regressor in the GLM-based method (Burgess et al., 2016) used to determine mean QC-FC and the association between QC-FC and ROI pair distance.

Results for analyses using uncensored data with no nuisance regressors (to show these results do not depend on effects of motion denoising pipelines) are reported in **Figure S14.** We repeated these analyses using 161 FH- participants matched instead by the ECDFs of both MP derivative and DV traces **(Figure S15)**, as well as by the mean absolute value of MP derivatives **(Figure S16)**. Regardless of the method used to match subjects based on motion, significant differences are observed when accounting for group membership in the majority of DQMs, suggesting that these metrics are critically confounded.

### *S2.2. Positive controls for volume censoring-based motion denoising in resting-state functional connectivity analyses*

#### *S2.2.1. Impact of censoring on functional connectivity correlations during resting and task states*

We calculated the inter-session correlations of functional connectivity as a function of data removed by volume censoring between session 2 resting-state runs and 1) session 1 resting-state runs, and 2) session 1 working memory task runs. **Figure S17A-B** shows the results of this analysis; the change from baseline (no censoring) was also calculated and is shown in **Figure S17C-D.** Because task data contains known neural signals that are absent from RSFC data, if volume censoring were removing true neural signal, then as censoring becomes more aggressive one might expect task and RSFC runs to become more similar. That is, as neural signal is removed from the task data by censoring, its similarity to RSFC should *increase*. As **Figure S17** shows, this phenomenon did not occur.

As expected, the correlation between resting-state runs ($Z_{RSFC\text{-}RSFC}$) is significantly greater than that between task runs and resting-state runs ($Z_{WM\text{-}RSFC}$) in uncensored data; this relationship persists regardless of the proportion of data removed by censoring. Notably, in comparison to $Z_{WM\text{-}RSFC}$, $Z_{RSFC\text{-}RSFC}$ declines more rapidly as volumes are censored. The comparative robustness of $Z_{WM\text{-}RSFC}$ to volume censoring could reflect one or more competing interpretations. First, there may be more motion artifact in resting-state data that is elevating correlations; however, comparison of mean FD, LPF-FD, DV, and LPF-DV values between session 1 working memory task runs and session 1 resting-state runs (with resting-state runs truncated to be the same duration as task runs) demonstrate that this is likely not the case, as motion did not differ significantly between task and RSFC runs, and was in fact greater in task runs (mean FD: task = 0.168 mm, rest = 0.159 mm, $p = 0.056$; mean LPF-FD: task = 0.0397 mm, rest = 0.0376 mm, $p = 0.367$; mean DV: task = 59.33, rest = 59.32, $p = 0.94$; LPF-DV: task = 10.70, rest = 10.55, $p = 0.26$; significance evaluated via Mann-Whitney-Wilcoxon rank-sum test). On the other hand, it is possible that RSFC data may contain greater variability in brain states than task data – for instance, individuals may fall asleep in the scanner more often during the resting-state than during a task. If these variable brain states are a diversion from a baseline, and if such a diversion is correlated with motion events, then volume censoring could be removing this distinct brain state information. However, the robustness of $Z_{WM\text{-}RSFC}$ provides strong evidence that, at least in the case of changes in brain activity caused by working memory task blocks, the ability to detect true differences in functional connectivity across conditions is maintained even when strict censoring criteria are applied.

*S2.2.2. Time of day effects on resting-state functional connectivity correlations*

Next, we evaluated the effect of volume censoring on the ability to detect differences in resting-state connectivity that exist between scan sessions that were acquired during a different time of day. It has previously been reported by Orban and associates (using the HCP dataset) acquiring resting-state BOLD data at a later time of day is associated with reductions in measured functional connectivity (across subjects) and global signal amplitude (both within and across subjects; Orban et al., 2020). In the HCP dataset, each subject underwent two scan sessions on different days, during an unstandardized time of day. Thus, the time of day at which session 2 resting-state scans took place relative to session 1 scans varies across subjects. We calculated the average association between the difference in scan time between session 1 and session 2, and the change in observed functional connectivity. The magnitude of this association, along with 95% confidence intervals, is shown in **Figure S18A-B**; the width of the 95% confidence intervals (calculated via bias-corrected and accelerated bootstrap; Efron and Tibshirani, 1993) is shown in **Figure S18C-D**. Because these RSFC differences as a function of scan time presumably result from true differences in neural signal as a function of arousal and circadian rhythm (or any other biological variable with a systematic relationship with time of day including, speculatively, time since caffeine intake), if excessive volume censoring results in removal of true neural signal, we might expect that magnitude of this effect to decrease and eventually be eliminated.

We indeed find a significant association between, within each subject, the difference in the time of day at which each RS scan session took place, and the change in magnitude of the average of RSFC correlations observed across sessions in data without using GSR (**Figure S18A-C**), though this average effect is not apparent when GSR is used (**Figure S18B-D**). On average, a Session 2 scan time that is later than Session 1 tends to produce an observed reduction in RSFC in Session 2 relative to Session 1, and vice versa. This is consistent as was found previously be Orban and associates (Orban et al., 2020), who observed that, without GSR, RSFC is negatively associated with a later scan time of day across subjects when analyzing data from each session separately, and that application of GSR largely removes this association as observed in aggregate (though some associations in specific networks were revealed or strengthened). Further, global signal magnitude was found to decrease with later scan time of day across subjects and within subjects (i.e., across scan sessions). Notably, in the case of data analyzed without GSR, the confidence interval width decreases through 60% frames removed, reflecting increased ability to resolve the true, population-average effect. While the magnitude of association does reduce slightly due to censoring, the change is non-significant and indistinguishable from the effect of removing systematic motion artifact.

*S2.3. Proportion of volumes censored over the course of fMRI runs*

**Figure S10** shows the percent of volumes censored across all runs for all subjects as a function of volume number within each run, separately for each run and each session, using the optimal volume censoring parameters for analyses without GSR in **Table 2**. Motion is minimal at the start of each run, and continues to increase over the course of each run, although the first 50-100 frames of Session A Run 1 (the first frames acquired for all subjects) are mildly elevated. Thus, when considering motion artifact, it appears that the highest-quality data is acquired at the start of each run, and continues to deteriorate due to declining subject adherence over the course of each scan. While the first run acquired in each session is of substantially higher quality until approximately 800 volumes (9.6 minutes of data) have been acquired, a substantial "rebound" towards higher quality data also occurs at the beginning of the second run of each session. Thus, it is advisable to employ a larger number of shorter runs rather than a smaller number of long runs. Specifically, **Figure S10** suggests that employing twice as many runs at half the length employed in the HCP study would have potentially resulted in substantially less participant motion than was observed.

It therefore appears from this analysis that, in order to maximize the quality of RSFC data, and maximize the quantity of data used for analysis after volume censoring, it is optimal to aim to maximize the number of runs within each session, and the number of sessions over which data are acquired. However, there are several practical considerations that must be weighed against this effect. Notably, participants must travel to the place of data acquisition for each session, and an increased number of sessions increases the burden on participants and study personnel. Further, by dividing sessions into multiple runs, "dead time" when the subject is in the fMRI facility but not actively undergoing acquisition procedures is increased. Finally, as volumes are discarded from the start of each run to allow for completion of tissue relaxation effects, and from the start and end of each run after band-pass filtering, increasing the number of runs during each session will increase the number of volumes discarded as part of routine data processing prior to volume censoring and analysis. Thus, in addition to consideration of limited resources for participants, study personnel, and facilities, there is a balance between the quantity of data that is acquired and the proportion of which is suitable for inclusion in analysis after volume censoring. In any case, our methods here do provide optimal methodology for volume censoring that an investigator may utilize either when prospectively designing such study protocols or while analyzing data following their implementation.

### Supplementary Tables

**Table S1.** Demographic information for the New York State Psychiatric Institute (NYSPI) Validation Dataset.

| Sample Size | Age (Mean ± SD) | Sex | Right Handedness | Race | Ethnicity |
|---|---|---|---|---|---|
| 26 | 32.58 ± 10 | F = 8, M = 18 | 92.3% | W = 9, AA = 10, As = 3, O = 4 | H = 7, NH = 19 |

*F = Female; M = Male; W = White; AA = African American; As = Asian; O = Other; H = Hispanic; NH = Non-Hispanic.*

**Table S2.** Optimal percent volumes censored with corresponding Standard Framewise Displacement (FD) and Low-Pass Filtered FD (LPF-FD) Parameter Values for candidate Dataset QC Metrics (DQMs).

| **Dataset QC Metric** | **Parameter** | **Optimal % Volumes Censored (Parameter Value)** | | **Figure** |
|---|---|---|---|---|
| | | **–GSR** | **+GSR** | |
| Median Absolute QC-FC | LPF-FD | 27.1003 % (4.4200) | 1.5604 % (17.5000) | *Figure 1A* |
| | Standard FD | 5.4103% (36.0000) | 0.1397% (107.5189) | |
| QC-FC Null Rejection Rate | LPF-FD | 17.3005 % (5.7200) | 0.874 % (22.0554) | *Figure 1B* |
| | Standard FD | 3.0571% (42.7500) | 0.0529% (154.3877) | |
| QC-FC Distance Correlation (Spearman) | LPF-FD | 21.6968 % (5.0400) | 25.3579 % (4.6000) | *Figure 1C* |
| | Standard FD | 6.8197% (33.5000) | 5.6641% (35.5000) | |
| High-Low Null Rejection Rate | LPF-FD | 29.6465 % (4.1800) | N/A | *Figure 1D* |
| | Standard FD | 5.2940% (36.2500) | 0.0185% (232.1430) | |
| Mean QC-FC (QC = mFD) | LPF-FD | 22.6342 % (4.9200) | 0.0226 % (100.0000) | *Figure 1E* |
| | Standard FD | 9.0972% (30.5000) | 0.0093% (310.0000) | |
| QC-FC Slope (GLM; QC = mFD) | LPF-FD | 26.3412% (4.4962) | 11.8414% (6.9900) | *Figure 1F* |
| | Standard FD | 7.1481% (33.0000) | 2.8294% (43.7500) | |
| Mean RSFC | LPF-FD | N/A | N/A | *Figure 1G* |
| | Standard FD | N/A | N/A | |
| RSFC Distance Dependence | LPF-FD | N/A | N/A | *Figure 1H* |
| | Standard FD | N/A | N/A | |
| Δ Variance | LPF-FD | 47.1728 % (3.0300) | 44.6103 % (3.1600) | *Figure 1I* |
| | Standard FD | 74.6565% (9.4900) | 63.6783% (11.2895) | |
| # Subjects Removed | LPF-FD | N/A | N/A | *Figure 1J* |
| | Standard FD | N/A | N/A | |
| Δ MSE-RSFC | LPF-FD | 47.4193 % (3.0179) | 44.6103 % (3.1600) | *Figure S11* |
| | Standard FD | 63.7587% (11.2773) | 59.6130% (12.0103) | N/A |

*Analysis performed using a subset of 475 subjects from the Human Connectome Project (HCP) 500 Subjects Release (HCP Dataset 1a). GSR = Global Signal Regression; QC-FC = Quality Control - Functional Connectivity; mFD = Mean Framewise Displacement; RSFC = Resting State Functional Connectivity; ICC = Intraclass Correlation; MSE-RSFC = Mean Squared Error-Resting State Functional Connectivity. "GLM" (Generalized Linear Model) denotes statistics generated using the GLM-based DQMs developed by Burgess and associates.*

**Table S3.** Optimal percent volumes censored with corresponding low-pass-filtered DV (LPF-DV) and Standard DV Parameter Values for candidate Dataset QC Metrics (DQMs).

| **Dataset QC Metric** | **Parameter** | **Optimal % Volumes Censored (Parameter Value)** | | **Figure** |
|---|---|---|---|---|
| | | **–GSR** | **+GSR** | |
| Median Absolute QC-FC | LPF-DV | 30.5152% (11.4972) | 53.2315% (10.5075) | *Figure S4A* |
| | Standard DV | 33.5124% (64.4000) | 44.0331% (61.7551) | |
| QC-FC Null Rejection Rate | LPF-DV | 55.0817% (10.4220) | 62.7919% (10.0432) | *Figure S4B* |
| | Standard DV | 56.9220% (58.3000) | 54.9114% (58.8000) | |
| QC-FC Distance Correlation (Spearman) | LPF-DV | 17.2422% (12.1692) | 17.0416% (12.1814) | *Figure S4C* |
| | Standard DV | 10.9608% (71.1000) | 6.3893% (73.5000) | |
| High-Low Null Rejection Rate | LPF-DV | 64.9890% (9.9300) | 60.6225% (10.1532) | *Figure S4D* |
| | Standard DV | 60.2431% (57.3440) | 59.1290% (57.7000) | |
| Mean QC-FC (QC = mFD) | LPF-DV | 19.9518% (12.0103) | 0.0228% (28.6720) | *Figure S4E* |
| | Standard DV | N/A | 0.0088% (155.4905) | |
| QC-FC Slope (GLM; QC = mFD) | LPF-DV | 78.6229% (9.2000) | 5.3269% (13.5000) | *Figure S4F* |
| | Standard DV | 67.5153% (54.8000) | 5.2544% (74.3000) | |
| Mean RSFC | LPF-DV | N/A | N/A | *Figure S4G* |
| | Standard DV | N/A | N/A | |
| RSFC Distance Dependence | LPF-DV | N/A | N/A | *Figure S4H* |
| | Standard DV | N/A | N/A | |
| Δ Variance | LPF-DV | 3.8559% (13.9000) | 3.0771% (14.2000) | *Figure S6A* |
| | Standard DV | 69.3936% (54.2000) | 3.6181% (75.9000) | |
| # Subjects Removed | LPF-DV | N/A | N/A | *Figure S6C* |
| | Standard DV | N/A | N/A | |
| Δ MSE-RSFC | LPF-DV | 4.8922% (13.6000) | 3.0771% (14.2000) | *N/A* |
| | Standard DV | 0.8641% (82.0000) | 0.7269% (82.7077) | |

*Analysis performed using a subset of 475 subjects from the Human Connectome Project (HCP) 500 Subjects Release (HCP Dataset 1a). GSR = Global Signal Regression; QC-FC = Quality Control - Functional Connectivity; mFD = Mean Framewise Displacement; RSFC = Resting State Functional Connectivity; ICC = Intraclass Correlation; MSE-RSFC = Mean Squared Error-Resting State Functional Connectivity. "GLM" (Generalized Linear Model) denotes statistics generated using the GLM-based DQMs developed by Burgess and associates*

**Table S4.** Optimal percent volumes censored with corresponding Generalized-Extreme-Value DV (GEV-DV) Parameter Values for candidate Dataset QC Metrics (DQMs).

| Dataset QC Metric | Optimal % Volumes Censored (GEV-DV Parameter Value) | | Figure |
|---|---|---|---|
| | –GSR | +GSR | |
| Median Absolute QC-FC | 55.2882 % (2.4314) | N/A | Figure S5A |
| QC-FC Null Rejection Rate | 55.2882 % (2.4314) | N/A | Figure S5B |
| QC-FC Distance Correlation (Spearman) | N/A | N/A | Figure S5C |
| High-Low Null Rejection Rate | 55.3262 % (2.4300) | N/A | Figure S5D |
| Mean QC-FC (QC = mFD) | N/A | 2.2638 % ($2.8672 \times 10^4$) | Figure S5E |
| QC-FC Slope (GLM; QC = mFD) | N/A | N/A | Figure S5F |
| Mean RSFC | N/A | N/A | Figure S5G |
| RSFC Distance Dependence | N/A | N/A | Figure S5H |
| Δ Variance | 43.0065 % (3.0000) | 38.1155 % ($1.4943 \times 10^3$) | Figure S6B |
| # Subjects Removed | N/A | N/A | Figure S6D |
| Δ MSE-RSFC | 43.1846 % (2.99) | 37.9486 % ($1.4998 \times 10^3$) | Figure S11 |

Analysis performed using a subset of 475 subjects from the Human Connectome Project (HCP) 500 Subjects Release (HCP Dataset 1a). GSR = Global Signal Regression; QC-FC = Quality Control - Functional Connectivity; mFD = Mean Framewise Displacement; RSFC = Resting State Functional Connectivity; ICC = Intraclass Correlation; MSE-RSFC = Mean Squared Error-Resting State Functional Connectivity. “GLM” (Generalized Linear Model) denotes statistics generated using the GLM-based DQMs developed by Burgess and associates.

### Supplementary Figures

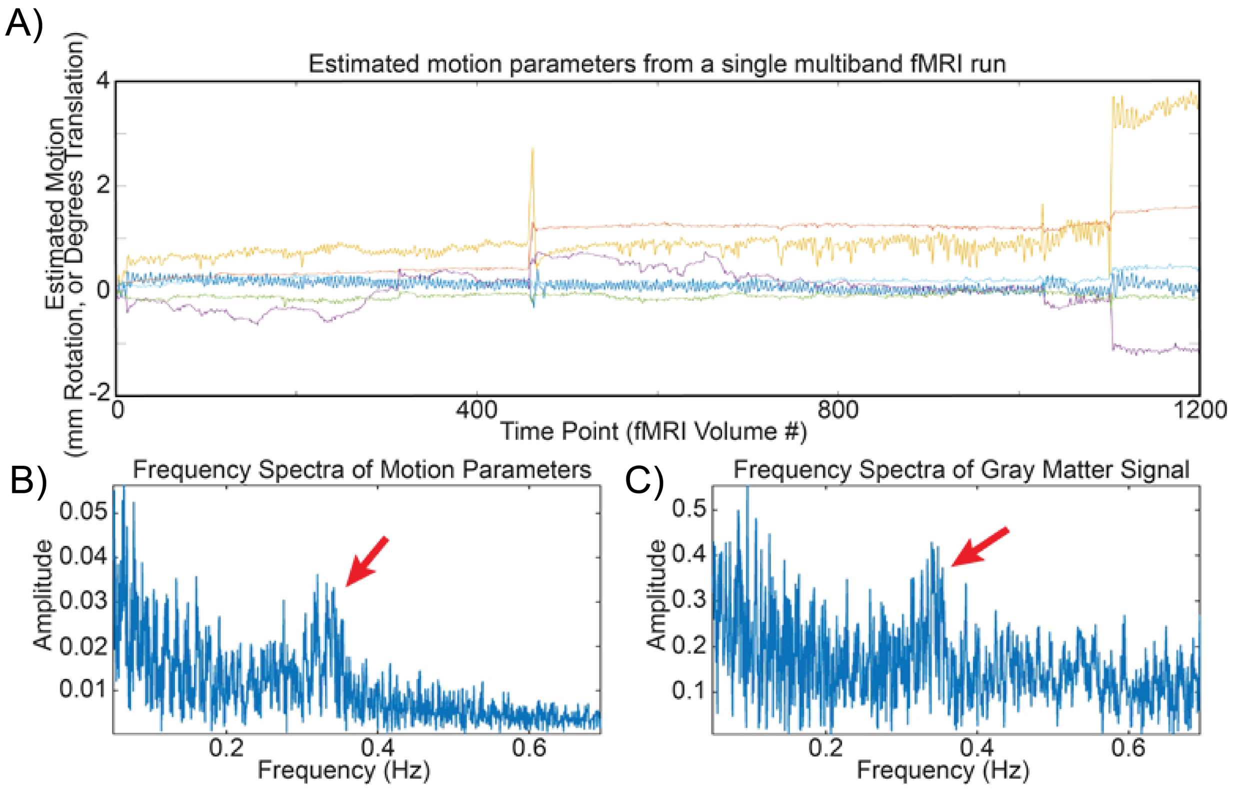


**Figure S1. A)** Estimated motion parameters (MPs) for a single RSFC scan from a participant in the Human Connectome Project (Subject ID 116524), which show a clear high-frequency oscillation throughout the run. **B)** Spectral frequency amplitudes of the MP traces, obtained via Fourier decomposition and summed over all 6 estimated MPs. **C)** Spectral frequency amplitudes of the mean signal extracted from all gray matter voxels after image preprocessing (see *Section 2.1.2*). Red arrows indicate the elevation of spectral amplitude in both MPs and global signal thought to result from participant respiration.

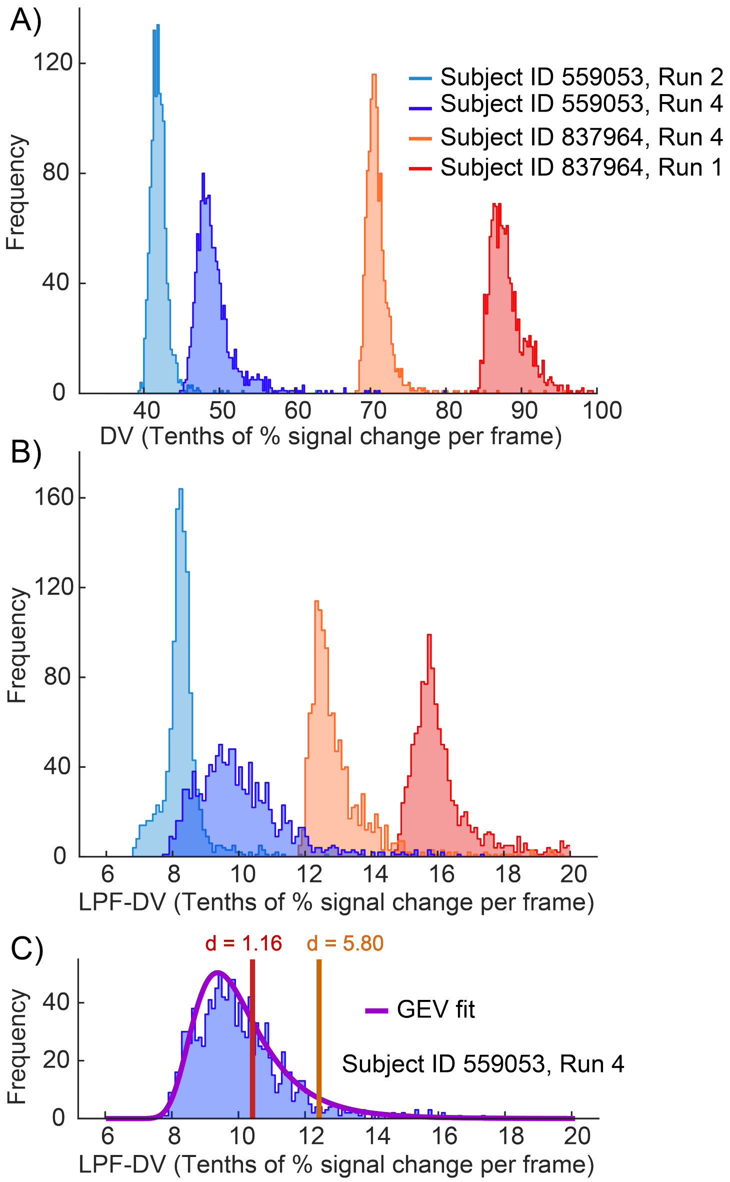


**Figure S2. A)** Histograms of framewise DV values for 4 runs, comprising 2 pairs of runs each from 2 subjects, demonstrating the large variability in central tendency of DV values between runs. **B)** Histograms of framewise LPF-DV for the same 4 runs as in A. **C)** Example generalized extreme value (GEV) distribution probability density function (PDF), determined from a maximum-likelihood fit, for one run (purple). Two adaptive censoring thresholds for this run are shown using (arbitrary unit) free parameter values d = 1.16 and d = 5.80.

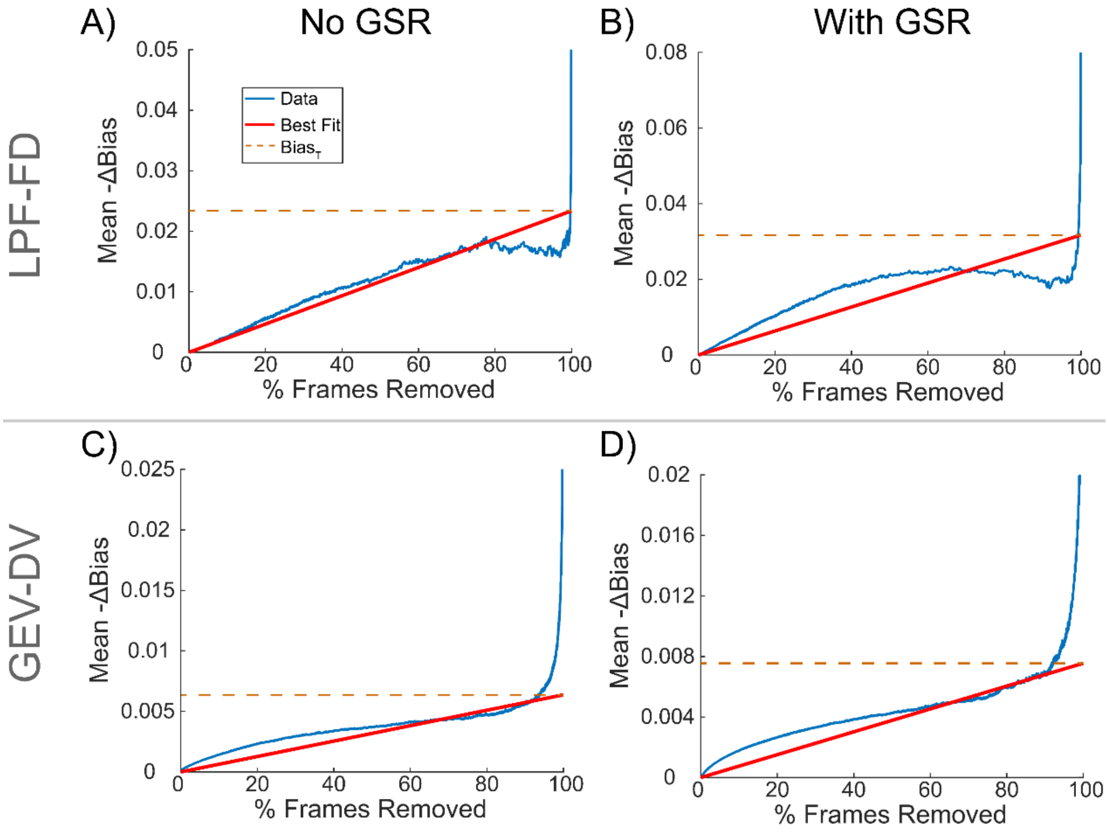


**Figure S3.** Demonstration of the procedure used to estimate the total motion artifact-induced bias in RSFC correlations (see *Section 2.3.2.4*), for LPF-FD **(A,B)** and GEV-DV **(C,D)**, both without GSR (**A,C)** and with GSR **(B,D)**. The blue line (Data) indicates the mean absolute value of the average change in RSFC correlations due to volume censoring, averaged across all ROI pairs. The red line (Best Fit) indicates the best fit using linear interpolation. The dashed line represents the y-value of the intersection of best Fit with 100% frames removed, corresponding to the estimated total motion-induced bias ($Bias_T$) in the sample removable by volume censoring using each method. $Bias(\hat{\theta}_{U_k}, \theta_k)$ was defined independently for analyses with and without GSR as $Bias_T$ using GEV-DV volume censoring.

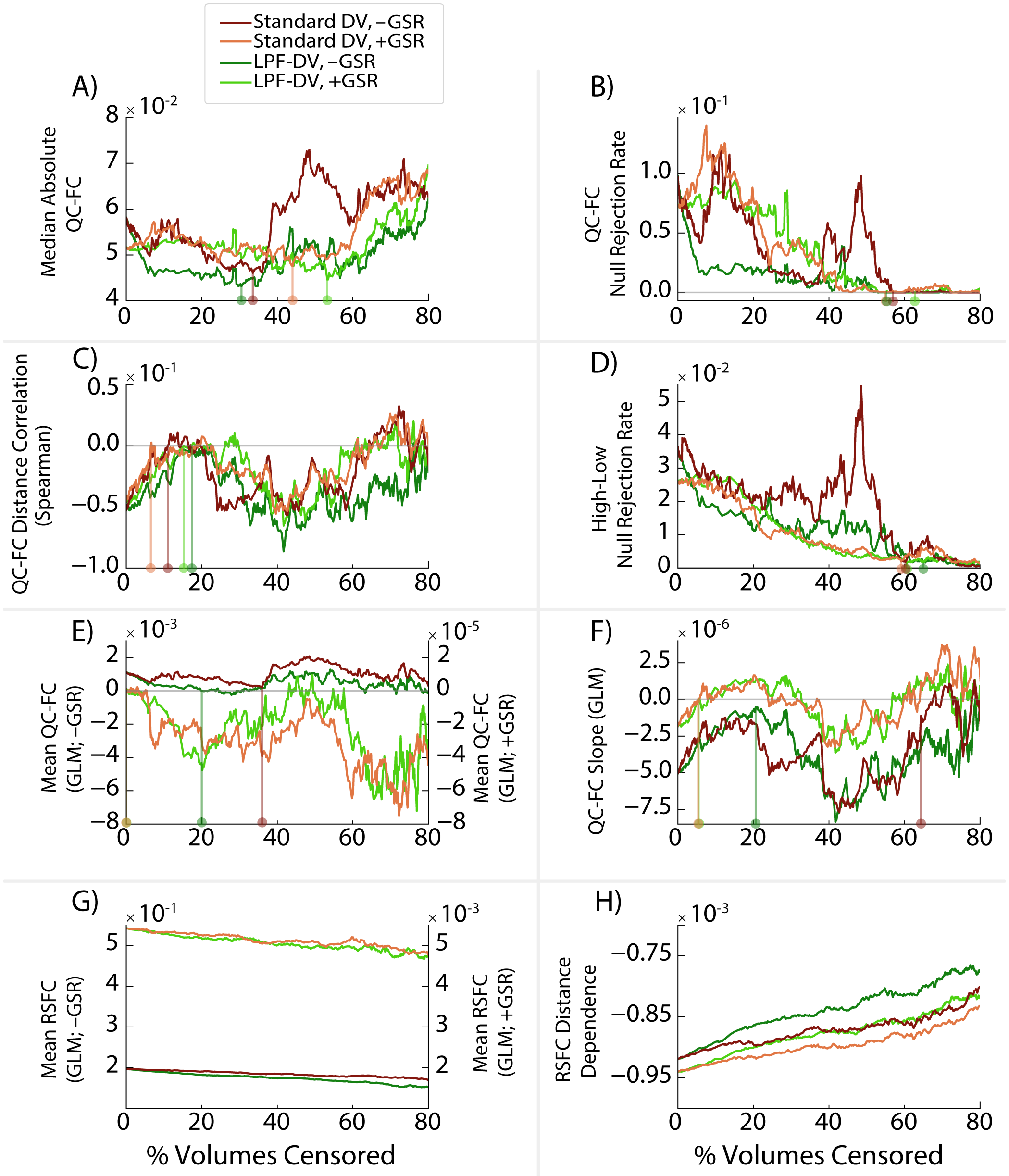


**Figure S4.** Dataset QC Metrics (DQMs) utilized in prior literature, calculated after volume censoring using either standard DV or low-pass-filtered DV (LPF-DV), plotted as a function of the percent of volumes censored in a subset of the HCP500 dataset (n = 475; *HCP Dataset 1a*). **A)** Median absolute QC-FC across ROI pairs, **B)** proportion of ROI pairs with significant QC-FC correlations (null rejection rate), **C)** correlation of QC-FC with the distance between ROI pairs, and **D)** proportion of ROI pairs showing significant differences between high-motion and low-motion participants. Also shown are DQMs calculated using a two-step generalized linear model (GLM), including: **E)** the mean of QC-FC correlations across the dataset, **F)** the distance-dependence (slope) of QC-FC correlations, **G)** the mean of RSFC correlations across the dataset, and **H)** the mean distance-dependence (slope) of RSFC correlations. Vertical lines show potential “optima” according to each DQM when identifiable in the region between 0% - 70% volumes censored. Results from analyses without global signal regression (GSR) are denoted by –GSR; results from analyses with GSR are denoted by +GSR.

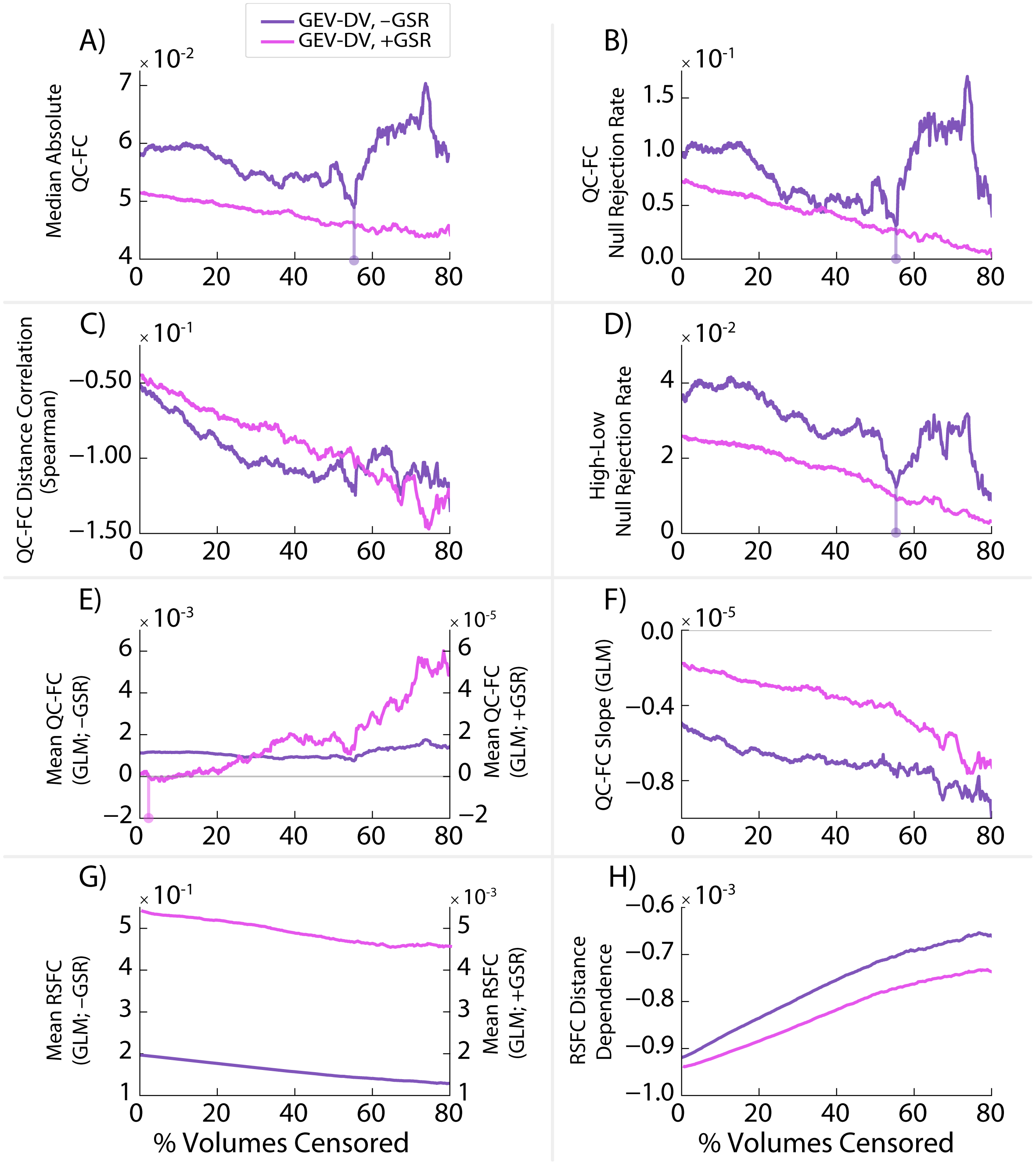


**Figure S5.** Dataset QC Metrics (DQMs) utilized in prior literature, calculated after volume censoring using GEV-DV plotted as a function of the percent of volumes censored in a subset of the HCP500 dataset (n = 475; *HCP Dataset 1a*). **A)** Median absolute QC-FC across ROI pairs, **B)** proportion of ROI pairs with significant QC-FC correlations (null rejection rate), **C)** correlation of QC-FC with the distance between ROI pairs, and **D)** proportion of ROI pairs showing significant differences between high-motion and low-motion participants. Also shown are DQMs calculated using a two-step generalized linear model (GLM), including: **E)** the mean of QC-FC correlations across the dataset, **F)** the distance-dependence (slope) of QC-FC correlations, **G)** the mean of RSFC correlations across the dataset, and **H)** the mean distance-dependence (slope) of RSFC correlations. Vertical lines show potential "optima" according to each DQM when identifiable in the region between 0% - 70% volumes censored. Results from analyses without global signal regression (GSR) are denoted by –GSR; results from analyses with GSR are denoted by +GSR.

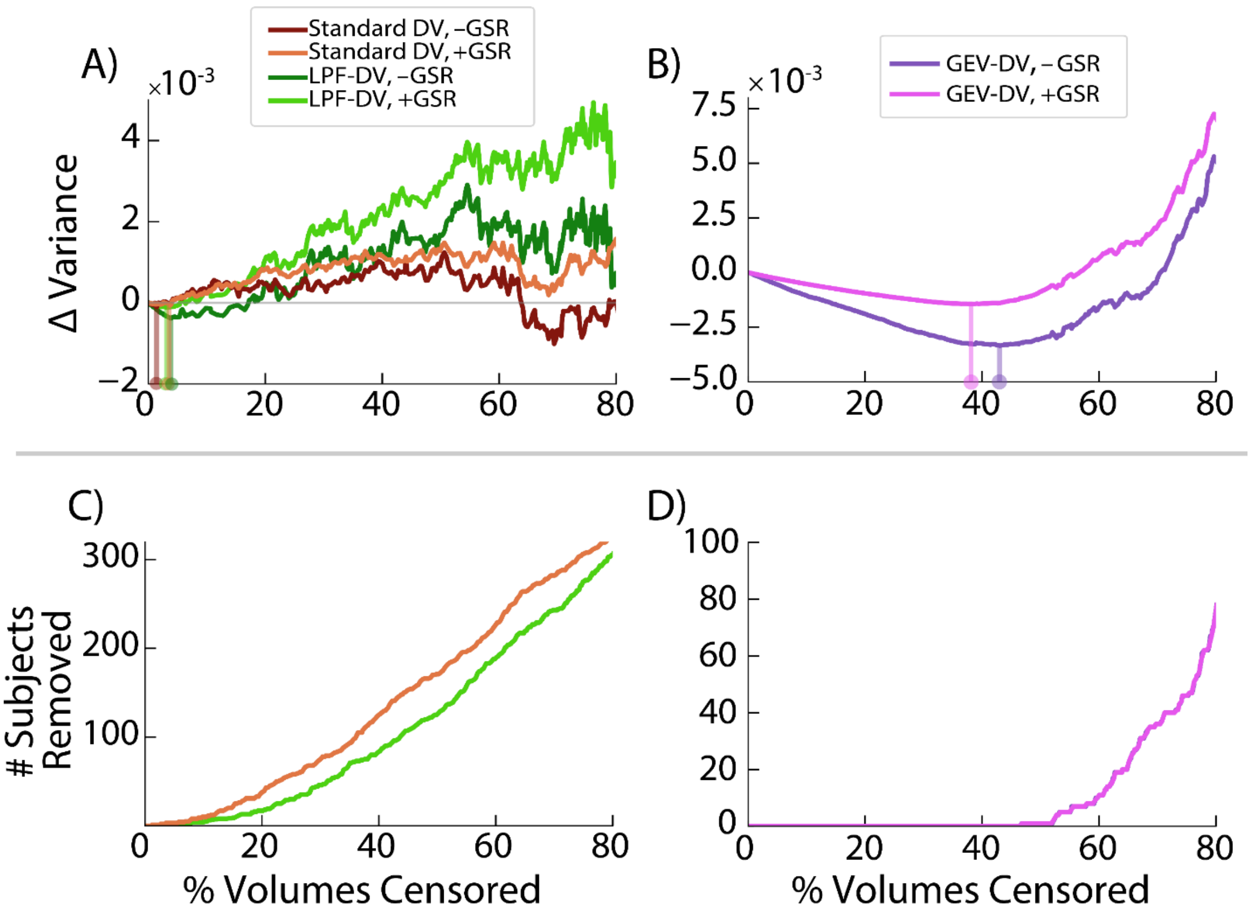


**Figure S6.** Change in observed between-subjects variance (ΔVariance) **(A,B),** and number of subjects removed **(C,D)**, due to standard DV and low-pass-filtered DV (LPF-DV) volume censoring **(A,C)** and GEV-DV volume censoring **(B,D)**, as a function of the percent of volumes censored in a subset of the HCP500 dataset (n = 475; *HCP Dataset 1a*). Vertical lines in Panels **A** and **B** show minima in the region between 0% and 70% volumes censored. Results from analyses without global signal regression (GSR) are denoted by –GSR; results from analyses with GSR are denoted by +GSR. Note: data in Panels **C** and **D** are identical for analyses with and without GSR, as GSR does not change which or how many volumes are censored.

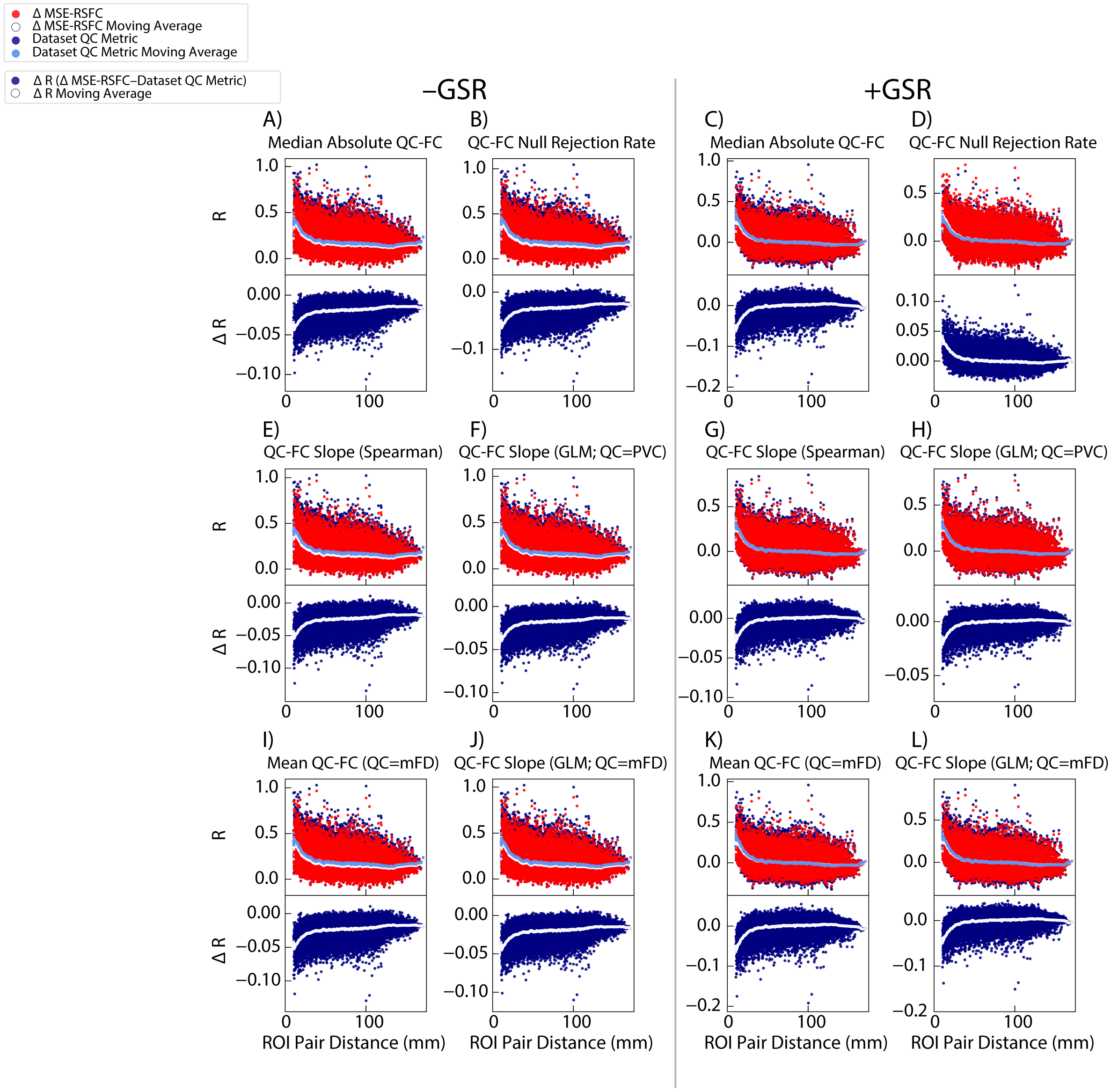


**Figure S7.** Resting-state functional connectivity (RSFC) correlations calculated in a subset of the HCP500 dataset (n = 475; *HCP Dataset 1a*) after volume censoring using an LPF-FD parameter value identified as "optimal" by ΔMSE-RSFC based optimization over and above censoring at LPF-FD parameter values identified as "optimal" by other traditional motion denoising benchmarks. Top panels of each vertical pair show pairwise ROI correlations for both ΔMSE-RSFC based censoring (red dots) with a moving average (white line) and a second metric (navy blue dots) with moving average (blue line). Bottom panels of each vertical pair show the difference in pairwise ROI correlations obtained between ΔMSE-RSFC based censoring and censoring based on the second metric (navy blue dots) with moving average (white line).

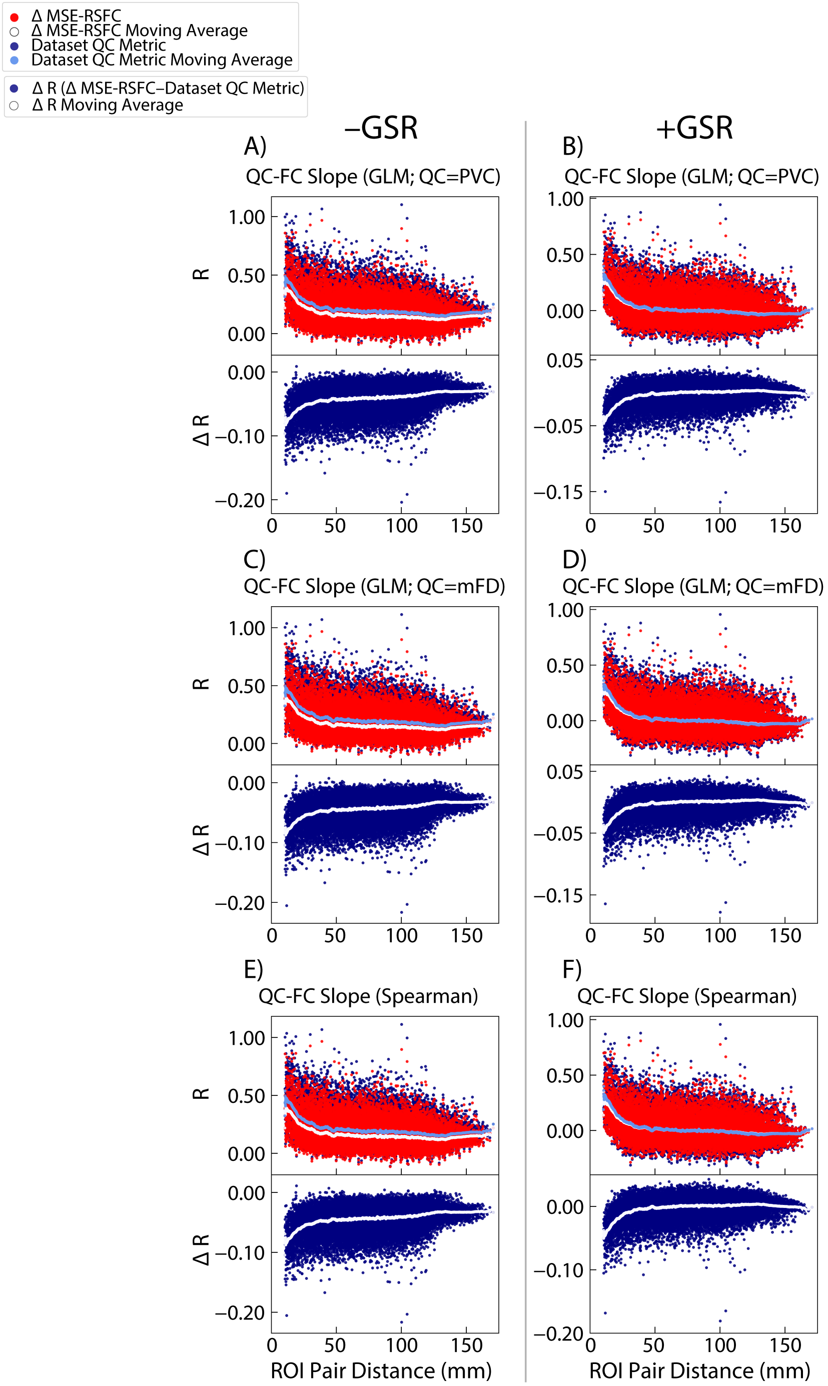


**Figure S8.** Resting-state functional connectivity (RSFC) correlations calculated in a subset of the HCP500 dataset (n = 475; *HCP Dataset 1a*) after volume censoring using a GEV-DV parameter value identified as "optimal" by ΔMSE-RSFC based optimization over and above censoring at GEV-DV parameter values identified as "optimal" by other traditional motion denoising benchmarks. Top panels of each vertical pair show pairwise ROI correlations for both ΔMSE-RSFC based censoring (red dots) with a moving average (white line) and a second metric (navy blue dots) with moving average (blue line). Bottom panels of each vertical pair show the difference in pairwise ROI correlations obtained between ΔMSE-RSFC based censoring and censoring based on the second metric (navy blue dots) with moving average (white line).

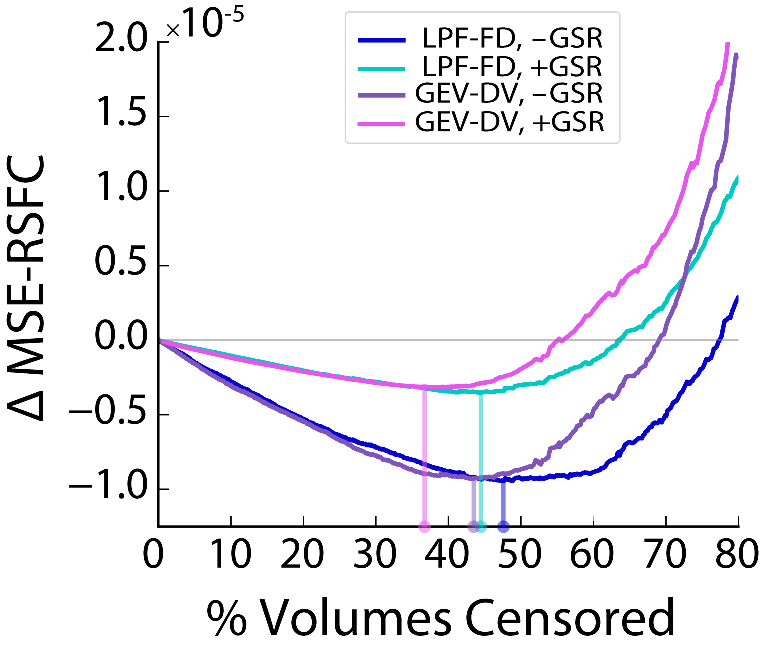


**Figure S9.** Change ΔMSE-RSFC due to volume censoring using either low-pass-filtered framewise displacement (LPF-FD) or GEV-DV, plotted as a function of the percent of volumes censored in a subset of the HCP500 dataset (*HCP Dataset 1a;* n = 475). Vertical lines show minima.

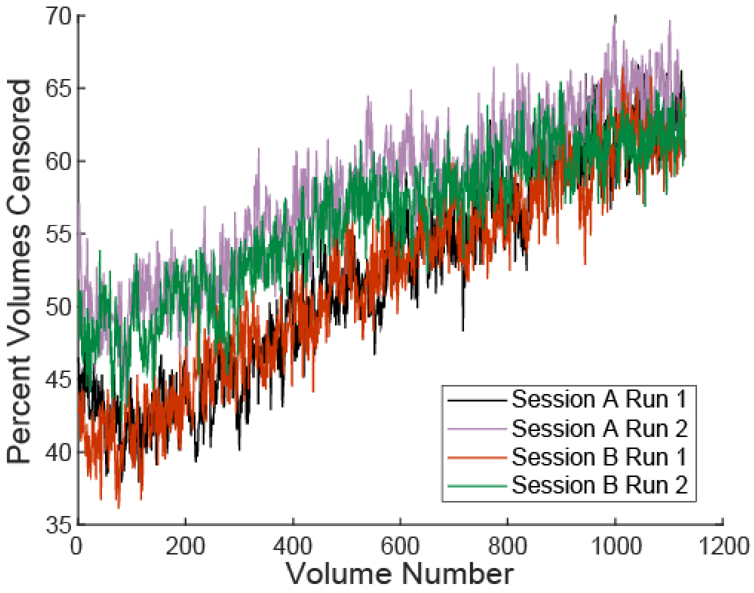


**Figure S10.** Percentage of times each given volume was censored across all subjects in the HCP500 dataset (n = 501; *HCP Dataset 1*) shown separately for each of the 4 runs, which were acquired over 2 sessions.

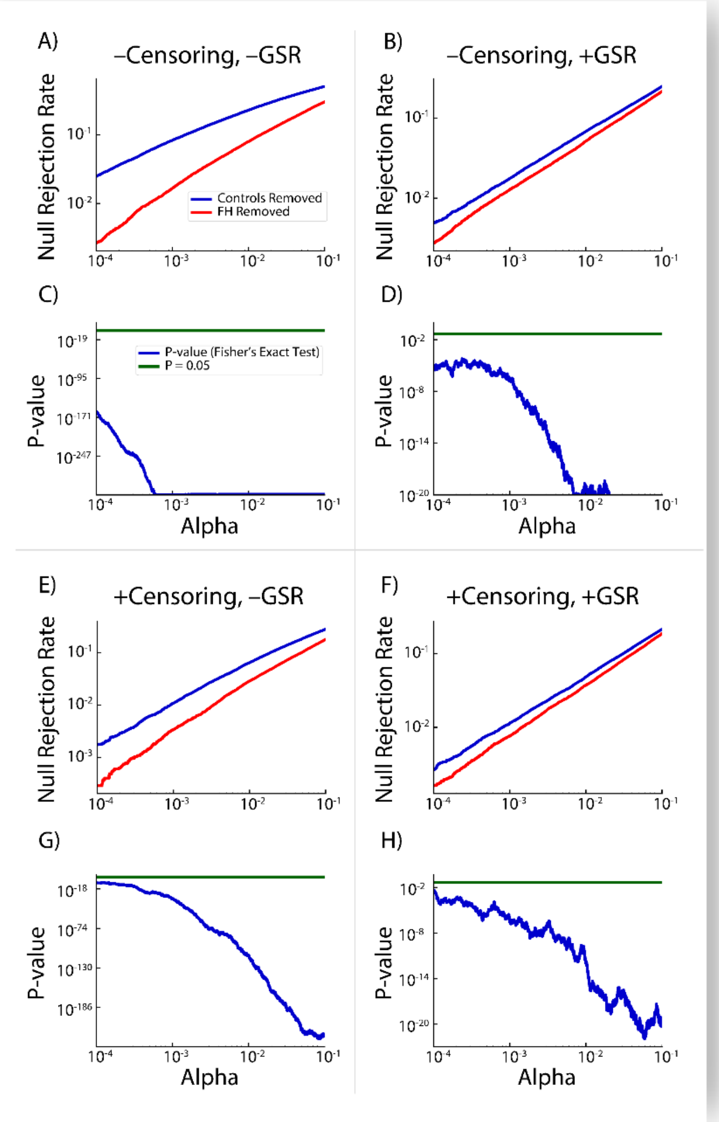


**Figure S11.** Proportion of RSFC correlations showing a significant difference between high- and low-motion subjects (Panels **A**, **B**, **E**, and **F**), as well as their p-values (Panels **C**, **D**, **G**, and **H**), in a subsample of unrelated subjects from the HCP 1200 Subjects Release (total n = 403; *HCP Dataset 2*), after removing either all subjects with a family history of a psychiatric or neurological disorder (FH+ removed; red lines), or FH- subjects (blue lines in C, D, G, and H) matched by the empirical cumulative density functions (ECDFs) of the derivatives of their motion parameter (MP) traces (in both groups, n = 242). Significance established using Fisher's exact test (blue lines in A, B, E, and F) with a nominal alpha level of p = 0.05 (green lines). Analyses without GSR are denoted by –GSR (left), and those with GSR are denoted by +GSR (right); analyses with volume censoring are denoted by –Censoring (top), and those with volume censoring are denoted by +Censoring (bottom).

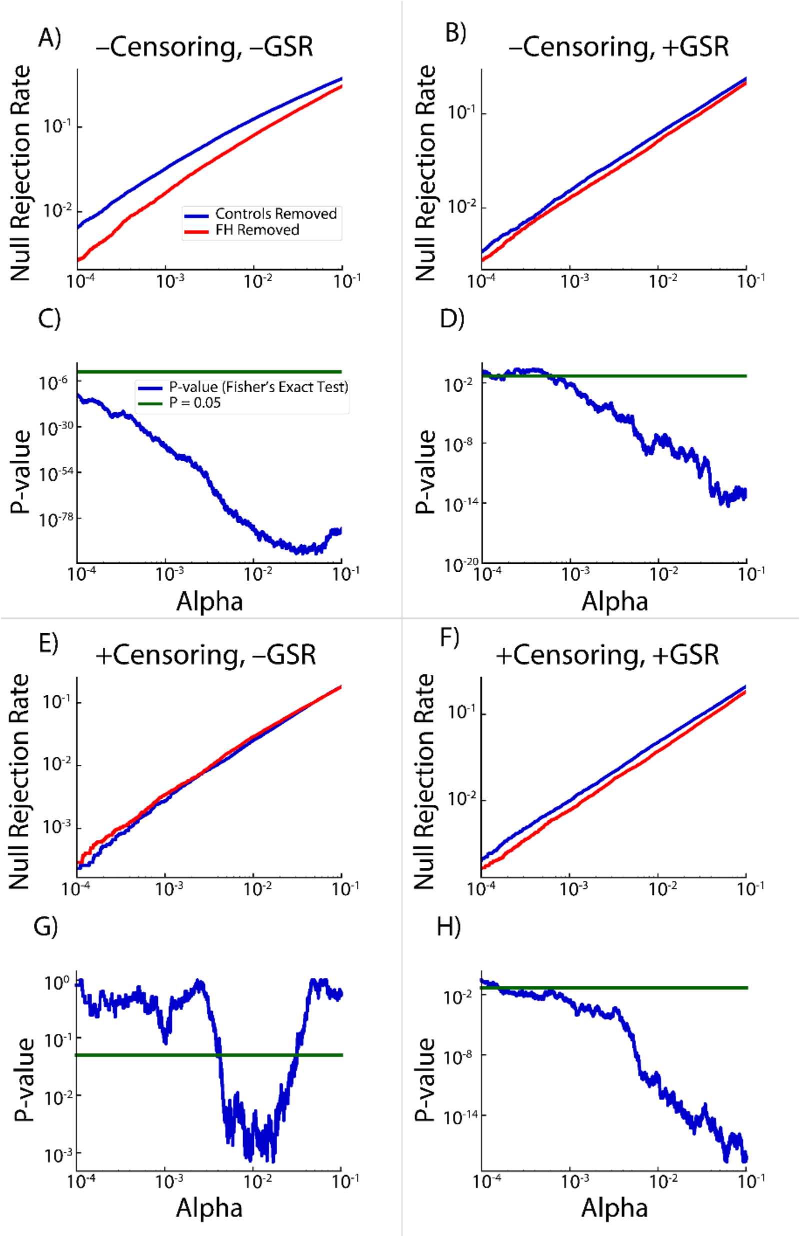


**Figure S12.** Proportion of RSFC correlations showing a significant difference between high- and low-motion subjects (**A**, **B**, **E**, and **F**), in a subsample of unrelated subjects from the HCP 1200 Subjects Release (total n = 403; *HCP Dataset 2*), after removing either all 161 subjects with a family history of psychiatric or neurological disease (FH+ Removed; red lines), or 161 control subjects optimally matched by their motion characteristics to the FH+ subjects (Controls Removed; blue lines). Panels **C**, **D**, **G**, and **H** show the significance level of the difference in proportion between FH+ and FH- groups as established using Fisher's exact test (blue lines) against the nominal alpha of 0.05 (green lines). Subjects were motion-matched using the empirical cumulative density functions (ECDFs) of the derivatives of their motion parameter (MP) traces together with the ECDFs of their DV traces (in both groups, n = 242). Analyses without GSR are denoted by –GSR (left), and those with GSR are denoted by +GSR (right). Analyses without volume censoring are denoted by –Censoring (top), and those with volume censoring are denoted by +Censoring (bottom).

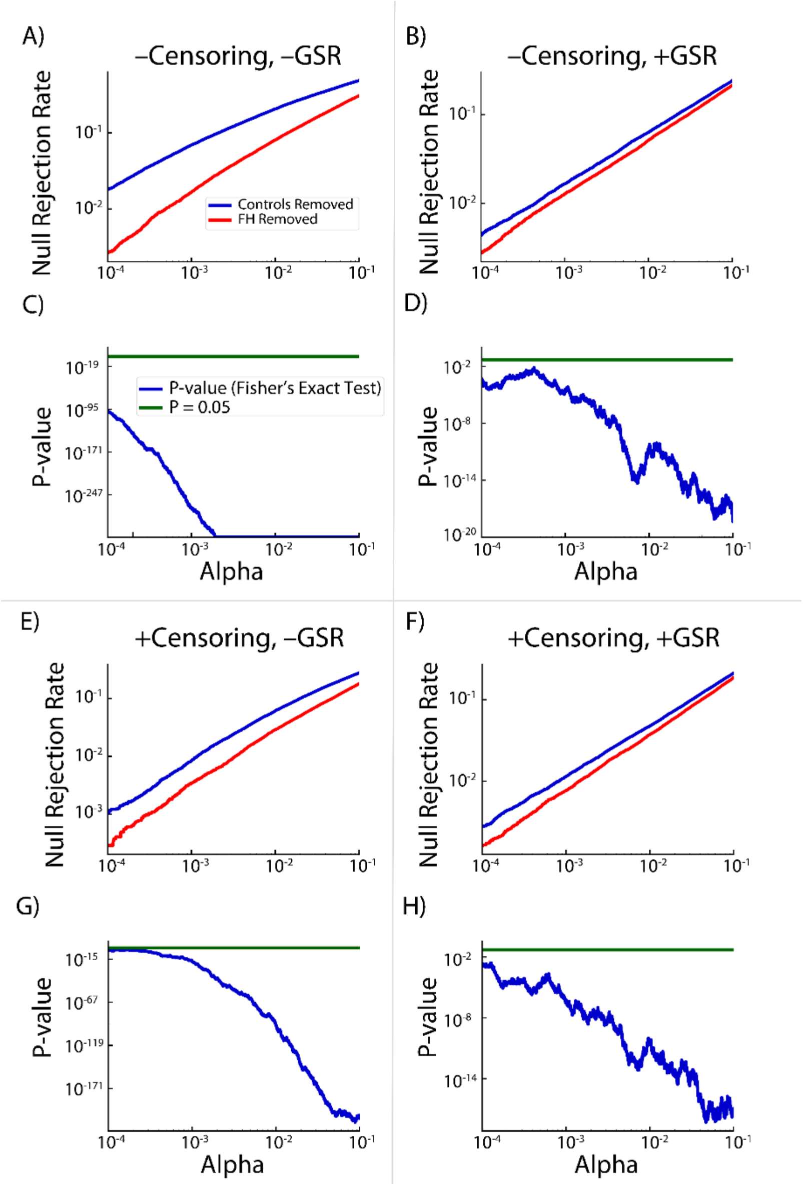


**Figure S13.** Proportion of RSFC correlations showing a significant difference between high- and low-motion subjects (**A**, **B**, **E**, and **F**), in a subsample of unrelated subjects from the HCP 1200 Subjects Release (total n = 403; *HCP Dataset 2*), after removing either all 161 subjects with a family history of psychiatric or neurological disease (FH+ Removed; red lines), or 161 control subjects optimally matched by their motion characteristics to the FH+ subjects (Controls Removed; blue lines). Panels **C**, **D**, **G**, and **H** show the significance level of the difference in proportion between FH+ and FH- groups as established using Fisher's exact test (blue lines) against the nominal alpha of 0.05 (green lines). Subjects were motion-matched using the means of the absolute values of their six motion parameter (MP) derivatives (in both groups, n = 242). Analyses without GSR are denoted by –GSR (left), and those with GSR are denoted by +GSR (right). Analyses without volume censoring are denoted by –Censoring (top), and those with volume censoring are denoted by +Censoring (bottom).

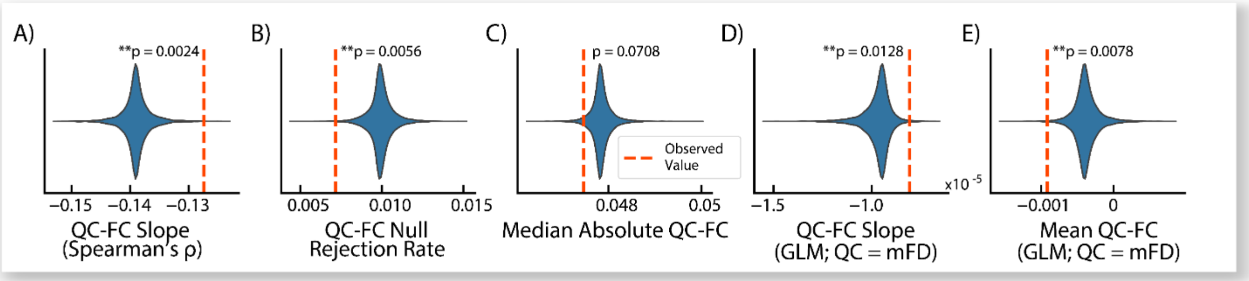


**Figure S14. Impact of modeling family history status on dataset quality metrics (DQMs) with subjects matched on motion parameter (MP) derivative distributions.** Violin plots show null distributions using 10,000 random permutations of family history (FH+) group membership assignments; dashed red lines show observed data values. Double asterisks (**) denote significance after false discovery rate (FDR) correction with α = 0.05. "GLM" (Generalized Linear Model) denotes statistics generated using the GLM-based DQMs developed by Burgess and associates, with mean FD used as the subject quality control (QC) metric. Results shown from a subset of the HCP1200 dataset (n = 322; *HCP Dataset 2a*) with 161 FH+ and 161 FH- subjects, matched by the empirical cumulative density functions (ECDFs) of the derivatives of their motion parameter traces.

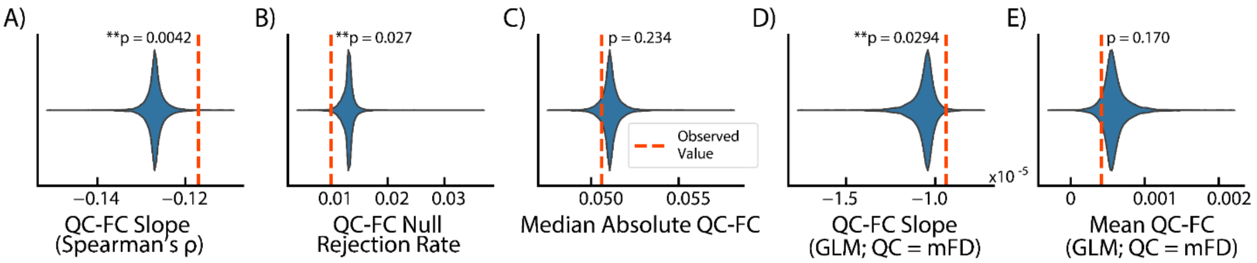


**Figure S15. Impact of modeling family history status on dataset quality metrics (DQMs) with subjects matched on both motion parameter (MP) derivative and DV trace distributions.** Violin plots show null distributions using 10,000 random permutations of family history (FH+) group membership assignments; dashed red lines show observed data values. Double asterisks (**) denote significance after false discovery rate (FDR) correction with α = 0.05. "GLM" (Generalized Linear Model) denotes statistics generated using the GLM-based DQMs developed by Burgess and associates, with mean FD used as the subject quality control (QC) metric. Results shown from a subset of the HCP1200 dataset (n = 322; *HCP Dataset 2b*) with 161 FH+ and 161 FH- subjects, matched by the empirical cumulative density functions (ECDFs) of the derivatives of their motion parameter traces.

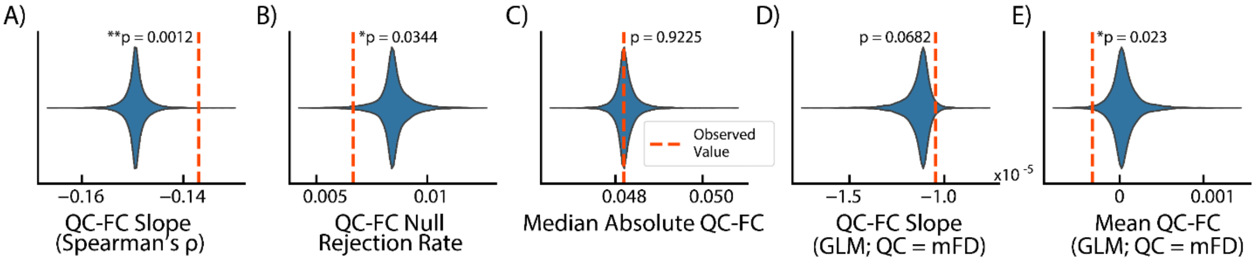


**Figure S16. Impact of modeling family history status on dataset quality metrics (DQMs) with subjects matched on the mean absolute value of their motion parameter (MP) derivatives.** Violin plots show null distributions using 10,000 random permutations of family history (FH+) group membership assignments; dashed red lines show observed data values. Double asterisks (**) denote significance after false discovery rate (FDR) correction with α = 0.05. "GLM" (Generalized Linear Model) denotes statistics generated using the GLM-based DQMs developed by Burgess and associates, with mean FD used as the subject quality control (QC) metric. Results shown from a subset of the HCP1200 dataset (n = 322; *HCP Dataset 2c*) with 161 FH+ and 161 FH- subjects, matched by the empirical cumulative density functions (ECDFs) of the derivatives of their motion parameter traces.

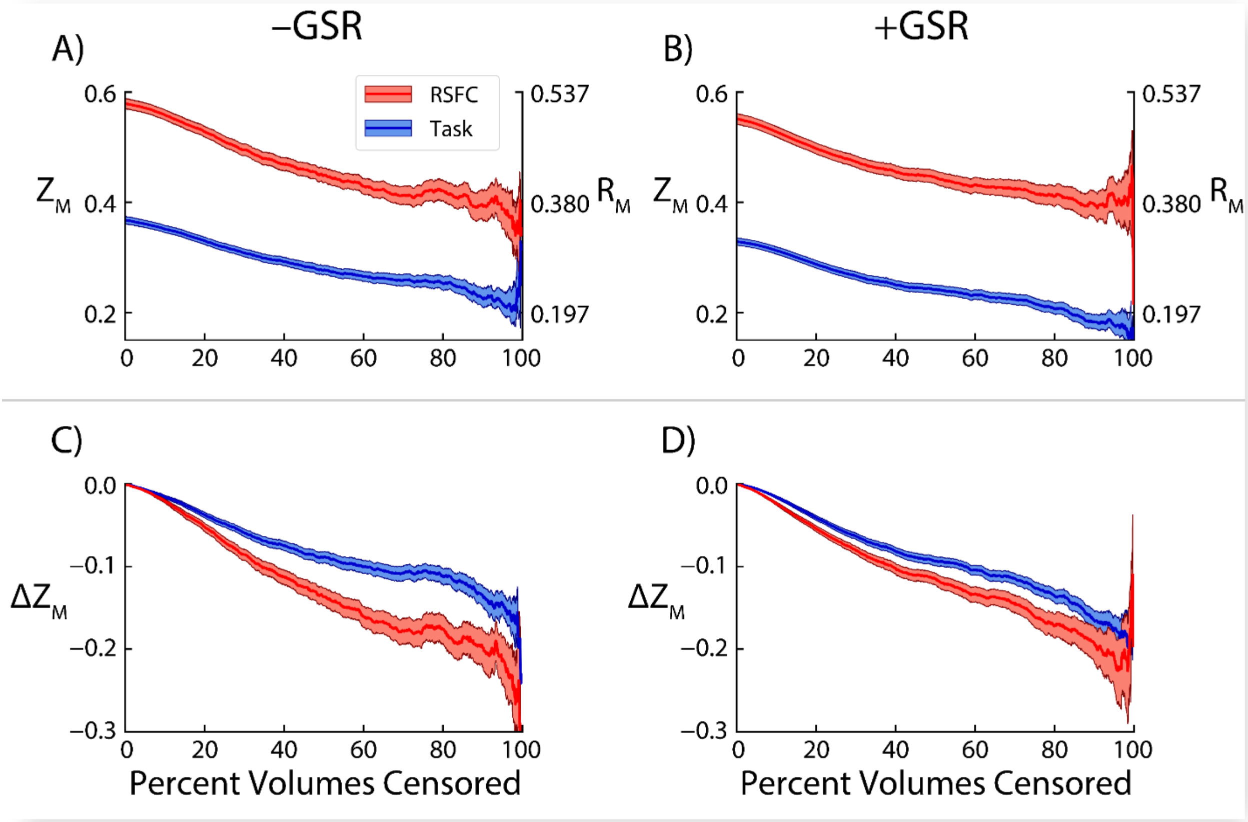


**Figure S17.** Comparison of the average effect of increasingly strict volume censoring on the correlation between working memory task (task) and resting state (RS) correlation matrices, in a subset of the HCP500 dataset (n = 330). Correlations were calculated within subjects, Fisher's Z-transformed, and then averaged across subjects. Panels **A** and **B** show the correlations between task (session 1) and RS (session 2) data (task-RS correlations; blue lines), as well as correlations between RS (session 1) and RS (session 2) data (RS-RS correlations; red lines), both before and after inverse Fisher's Z-transformation ($Z_M$ and $R_M$, respectively). Panels **C** and **D** show the change in (Z-transformed) task-RS and RS-RS correlations relative to baseline uncensored data ($\Delta Z_M$). Results from analyses without global signal regression (GSR) are denoted -GSR (left); results from analyses without GSR are denoted +GSR (right).

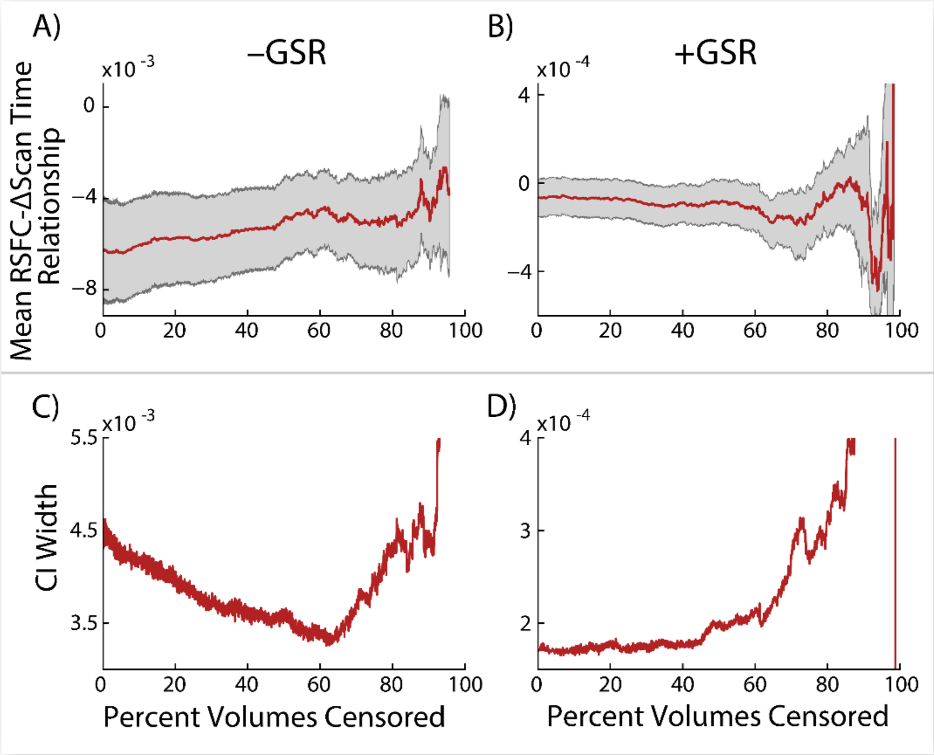


**Figure S18.** Average magnitude of association between the change in time between scan sessions and the resulting change in RSFC correlations in a subset of the HCP500 dataset (n = 478) are shown in panels **A** and **B**, along with associated 95% confidence intervals estimated via bootstrapping. Panels **C** and **D** show the corresponding width of the CI interval width.

### Supplementary References

Abramson, M., Audet, C., Dennis, J., Digabel, S., 2009. OrthoMADS: A Deterministic MADS Instance with Orthogonal Directions. SIAM Journal on Optimization 20, 948-966.

Anderson, T.W., 1962. On the Distribution of the Two-Sample Cramér-von Mises Criterion. The Annals of Mathematical Statistics 33, 1148-1159.

Association, A.P., 2000. Diagnostic and statistical manual of mental disorders : DSM-IV-TR, 4th ed., text revision. ed. American Psychiatric Association, Washington, DC.

Audet, C., Dennis, J., 2002. Analysis of Generalized Pattern Searches. SIAM Journal on Optimization 13, 889-903.

Barch, D.M., Burgess, G.C., Harms, M.P., Petersen, S.E., Schlaggar, B.L., Corbetta, M., Glasser, M.F., Curtiss, S., Dixit, S., Feldt, C., Nolan, D., Bryant, E., Hartley, T., Footer, O., Bjork, J.M., Poldrack, R., Smith, S., Johansen-Berg, H., Snyder, A.Z., Van Essen, D.C., Consortium, W.U.-M.H., 2013. Function in the human connectome: task-fMRI and individual differences in behavior. Neuroimage 80, 169-189.

Benjamini, Y., Hochberg, Y., 1995. Controlling the False Discovery Rate: A Practical and Powerful Approach to Multiple Testing. Journal of the Royal Statistical Society: Series B (Methodological) 57, 289-300.

Burgess, G.C., Kandala, S., Nolan, D., Laumann, T.O., Power, J.D., Adeyemo, B., Harms, M.P., Petersen, S.E., Barch, D.M., 2016. Evaluation of Denoising Strategies to Address Motion-Correlated Artifacts in Resting-State Functional Magnetic Resonance Imaging Data from the Human Connectome Project. Brain Connect 6, 669-680.

Ciric, R., Wolf, D.H., Power, J.D., Roalf, D.R., Baum, G.L., Ruparel, K., Shinohara, R.T., Elliott, M.A., Eickhoff, S.B., Davatzikos, C., Gur, R.C., Gur, R.E., Bassett, D.S., Satterthwaite, T.D., 2017. Benchmarking of participant-level confound regression strategies for the control of motion artifact in studies of functional connectivity. Neuroimage 154, 174-187.

Conn, A.R., Gould, N., Toint, P.L., 1997. A globally convergent Lagrangian barrier algorithm for optimization with general inequality constraints and simple bounds. Math. Comput. 66, 261-288.

Davidon, W., 1991. Variable Metric Method for Minimization. SIAM Journal on Optimization 1, 1-17.

Duff, I., Koster, J., 2001. On Algorithms For Permuting Large Entries to the Diagonal of a Sparse Matrix. SIAM J. Matrix Anal. Appl. 22, 973-996.

Efron, B., Tibshirani, R., 1993. An introduction to the bootstrap. Chapman & Hall, New York.

First, M.B., Gibbon, M., 2004. The Structured Clinical Interview for DSM-IV Axis I Disorders (SCID-I) and the Structured Clinical Interview for DSM-IV Axis II Disorders (SCID-II). Comprehensive handbook of psychological assessment, Vol. 2: Personality assessment. John Wiley & Sons, Inc., Hoboken, NJ, US, pp. 134-143.

Glasser, M.F., Coalson, T.S., Bijsterbosch, J.D., Harrison, S.J., Harms, M.P., Anticevic, A., Van Essen, D.C., Smith, S.M., 2018. Using temporal ICA to selectively remove global noise while preserving global signal in functional MRI data. Neuroimage 181, 692-717.

Glasser, M.F., Sotiropoulos, S.N., Wilson, J.A., Coalson, T.S., Fischl, B., Andersson, J.L., Xu, J., Jbabdi, S., Webster, M., Polimeni, J.R., Van Essen, D.C., Jenkinson, M., Consortium, W.U.-M.H., 2013. The minimal preprocessing pipelines for the Human Connectome Project. Neuroimage 80, 105-124.

Grabner, G., Janke, A.L., Budge, M.M., Smith, D., Pruessner, J., Collins, D.L., 2006. Symmetric atlasing and model based segmentation: an application to the hippocampus in older adults. Med Image Comput Comput Assist Interv 9, 58-66.

Hodge, M.R., Horton, W., Brown, T., Herrick, R., Olsen, T., Hileman, M.E., McKay, M., Archie, K.A., Cler, E., Harms, M.P., Burgess, G.C., Glasser, M.F., Elam, J.S., Curtiss, S.W., Barch, D.M., Oostenveld, R., Larson-Prior, L.J., Ugurbil, K., Van Essen, D.C., Marcus, D.S., 2016. ConnectomeDB--Sharing human brain connectivity data. Neuroimage 124, 1102-1107.

Ingber, L., 1996. Adaptive simulated annealing (ASA): Lessons learned. Control Cybernetics 25, 33-54.

Kennedy, J., Eberhart, R., 1995. Particle swarm optimization. Proceedings of ICNN'95 - International Conference on Neural Networks, pp. 1942-1948 vol.1944.

Kirkpatrick, S., Gelatt, C.D., Jr., Vecchi, M.P., 1983. Optimization by simulated annealing. Science 220, 671-680.

Kolda, T., Lewis, R., Torczon, V., 2003. Optimization by Direct Search: New Perspectives on Some Classical and Modern Methods. SIAM Review 45, 385-482.

Lewis, R., Shepherd, A., Torczon, V., 2007. Implementing Generating Set Search Methods for Linearly Constrained Minimization. SIAM Journal on Scientific Computing 29, 2507-2530.

Metropolis, N., Rosenbluth, A.W., Rosenbluth, M.N., Teller, A.H., Teller, E., 1953. Equation of State Calculations by Fast Computing Machines. The Journal of Chemical Physics 21, 1087-1092.

Mezura-Montes, E., Coello Coello, C.A., 2011. Constraint-handling in nature-inspired numerical optimization: Past, present and future. Swarm and Evolutionary Computation 1, 173-194.

Muschelli, J., Nebel, M.B., Caffo, B.S., Barber, A.D., Pekar, J.J., Mostofsky, S.H., 2014. Reduction of motion-related artifacts in resting state fMRI using aCompCor. Neuroimage 96, 22-35.

Orban, C., Kong, R., Li, J., Chee, M.W.L., Yeo, B.T.T., 2020. Time of day is associated with paradoxical reductions in global signal fluctuation and functional connectivity. PLoS Biol 18, e3000602.

Parkes, L., Fulcher, B., Yucel, M., Fornito, A., 2018. An evaluation of the efficacy, reliability, and sensitivty of motion correction strategies for resting-state functional MRI. Neuroimage 171, 415-436.

Power, J.D., Barnes, K.A., Snyder, A.Z., Schlaggar, B.L., Petersen, S.E., 2012. Spurious but systematic correlations in functional connectivity MRI networks arise from subject motion. Neuroimage 59, 2142-2154.

Power, J.D., Mitra, A., Laumann, T.O., Snyder, A.Z., Schlaggar, B.L., Petersen, S.E., 2014. Methods to detect, characterize, and remove motion artifact in resting state fMRI. Neuroimage 84, 320-341.

Pruim, R.H.R., Mennes, M., Buitelaar, J.K., Beckmann, C.F., 2015. Evaluation of ICA-AROMA and alternative strategies for motion artifact removal in resting state fMRI. Neuroimage 112, 278-287.

Robinson, E.C., Garcia, K., Glasser, M.F., Chen, Z., Coalson, T.S., Makropoulos, A., Bozek, J., Wright, R., Schuh, A., Webster, M., Hutter, J., Price, A., Cordero Grande, L., Hughes, E., Tusor, N., Bayly, P.V., Van Essen, D.C., Smith, S.M., Edwards, A.D., Hajnal, J., Jenkinson, M., Glocker, B., Rueckert, D., 2018. Multimodal surface matching with higher-order smoothness constraints. Neuroimage 167, 453-465.

Robinson, E.C., Jbabdi, S., Glasser, M.F., Andersson, J., Burgess, G.C., Harms, M.P., Smith, S.M., Van Essen, D.C., Jenkinson, M., 2014. MSM: a new flexible framework for Multimodal Surface Matching. Neuroimage 100, 414-426.

Satterthwaite, T.D., Elliott, M.A., Gerraty, R.T., Ruparel, K., Loughead, J., Calkins, M.E., Eickhoff, S.B., Hakonarson, H., Gur, R.C., Gur, R.E., Wolf, D.H., 2013. An improved framework for confound regression and filtering for control of motion artifact in the preprocessing of resting-state functional connectivity data. Neuroimage 64, 240-256.

Shellock, F.G., 1998. Pocket guide to MR procedures and metallic objects: update 1998. Lippincott-Raven, Philadelphia.

Shi, Y., Eberhart, R., 1998. A modified particle swarm optimizer. 1998 IEEE International Conference on Evolutionary Computation Proceedings. IEEE World Congress on Computational Intelligence (Cat. No.98TH8360), pp. 69-73.

Smith, S.M., Beckmann, C.F., Andersson, J., Auerbach, E.J., Bijsterbosch, J., Douaud, G., Duff, E., Feinberg, D.A., Griffanti, L., Harms, M.P., Kelly, M., Laumann, T., Miller, K.L., Moeller, S., Petersen, S., Power, J., Salimi-Khorshidi, G., Snyder, A.Z., Vu, A.T., Woolrich, M.W., Xu, J., Yacoub, E., Ugurbil, K., Van Essen, D.C., Glasser, M.F., Consortium, W.U.-M.H., 2013. Resting-state fMRI in the Human Connectome Project. Neuroimage 80, 144-168.

Ugurbil, K., Xu, J., Auerbach, E.J., Moeller, S., Vu, A.T., Duarte-Carvajalino, J.M., Lenglet, C., Wu, X., Schmitter, S., Van de Moortele, P.F., Strupp, J., Sapiro, G., De Martino, F., Wang, D., Harel, N., Garwood, M., Chen, L., Feinberg, D.A., Smith, S.M., Miller, K.L., Sotiropoulos, S.N., Jbabdi, S., Andersson, J.L., Behrens, T.E., Glasser, M.F., Van Essen, D.C., Yacoub, E., Consortium, W.U.-M.H., 2013. Pushing spatial and temporal resolution for functional and diffusion MRI in the Human Connectome Project. Neuroimage 80, 80-104.

Van Dijk, K.R., Sabuncu, M.R., Buckner, R.L., 2012. The influence of head motion on intrinsic functional connectivity MRI. Neuroimage 59, 431-438.

Van Essen, D.C., Smith, S.M., Barch, D.M., Behrens, T.E., Yacoub, E., Ugurbil, K., Consortium, W.U.-M.H., 2013. The WU-Minn Human Connectome Project: an overview. Neuroimage 80, 62-79.
